# Designing and interpreting 4D tumour spheroid experiments

**DOI:** 10.1101/2021.08.18.456910

**Authors:** Ryan J. Murphy, Alexander P. Browning, Gency Gunasingh, Nikolas K. Haass, Matthew J. Simpson

**Affiliations:** Mathematical Sciences, Queensland University of Technology, Brisbane, Australia; The University of Queensland Diamantina Institute, The University of Queensland, Brisbane, Australia

## Abstract

Tumour spheroid experiments are routinely used to study cancer progression and treatment. Various and inconsistent experimental designs are used, leading to challenges in interpretation and reproducibility. Using multiple experimental designs, live-dead cell staining, and real-time cell cycle imaging, we measure necrotic and proliferation-inhibited regions in over 1000 4D tumour spheroids (3D space plus cell cycle status). By intentionally varying the initial spheroid size and temporal sampling frequencies across multiple cell lines, we collect an abundance of measurements of internal spheroid structure. These data are difficult to compare and interpret. However, using an objective mathematical modelling framework and statistical identifiability analysis we quantitatively compare experimental designs and identify design choices that produce reliable biological insight. Measurements of internal spheroid structure provide the most insight, whereas varying initial spheroid size and temporal measurement frequency is less important. Our general framework applies to spheroids grown in different conditions and with different cell types.

## 1 Introduction

Tumour spheroid experiments are an important *in vitro* tool routinely used since the 1970s to understand avascular tumour growth, cancer progression, develop cancer treatments, and reduce animal experimentation [1–10]. However, a vast range of experimental designs are employed, leading to inconsistencies in: i) the times when measurements are taken; ii) experimental durations, ranging from a few days to over a month [11–16]; iii) the initial number of cells used to form spheroids [11–18], commonly between 300 to 20, 000 cells [16, 17]; and, iv) the type of experimental measurements that are taken [11–18]. This variability in experimental protocols makes comparing different studies very difficult, and introduces challenges in both interpretation and reproducibility of these experiments.

Mathematical modelling provides a powerful tool to provide such intepretation through model calibration and mechanism deduction. Simple mathematical models calibrated to outer radius measurements, such as Gompertzian growth models, have been used for decades to predict the growth of tumours [19, 20]. However, these simple mathematical models do not provide information about the internal spheroid structure over time. In response, many mathematical models of varying complexity have been developed to explore the internal structure of spheroids [21–42]. Here, we revisit the seminal Greenspan mathematical model for avascular tumour spheroid growth [21] and quantitatively directly connect it to data for the first time. Greenspan’s mathematical model was the first to describe the three phases of avascular tumour spheroid growth: in phase (i) cells throughout the spheroid can proliferate; in phase (ii) cells near the periphery proliferate while a central region of living cells cannot proliferate, referred to as the inhibited region; and in phase (iii) there is an outer region of proliferative cells, an intermediate region of living inhibited cells, and a central necrotic region composed of dead cells and cellular material in various stages of disintegration and dissolution (Figure 1a-d, Methods 4.1). These various regions of cellular behaviour are thought to arise as a result of nutrient availability, such as oxygen, that is driven by diffusion and uptake. Using Greenspan’s mathematical model has great advantages. All parameters are physically interpretable and biologically relevant, as opposed to more complicated mathematical models that may have parameters that cannot be interpreted physically and cannot be identified with the data in this study [27]. In addition, without using mathematical modelling and uncertainty quantification such that we employ here it is not obvious how to quantify the value of different experimental designs, and therefore impossible to interpret and determine the uncertainty of biologically relevant parameters.

**Figure 1:**
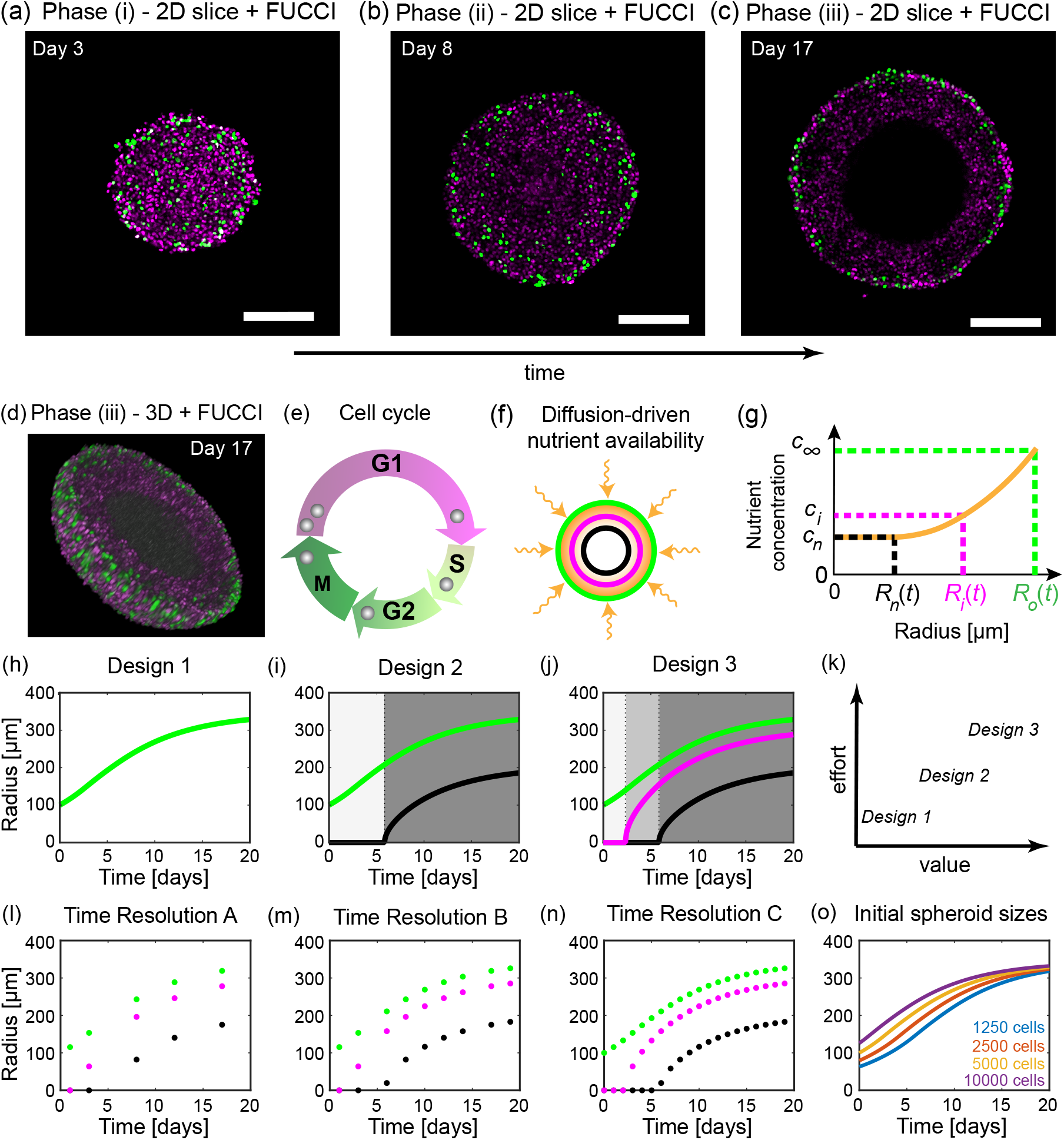
Tumour spheroid growth and the Greenspan mathematical model. Tumour spheroids experience three phases of growth. (a)-(d) Confocal microscopy reveals different phases of tumour growth. Fluorescent ubiquitination-based cell cycle indicator (FUCCI) transduced cells allow visualisation of each cell’s stage in the cell cycle. (a)-(c) 2D equatorial plane images of WM793b human melanoma tumour spheroids, formed with 5000 cells per spheroid, on days 3, 8, and 17 after formation. Scale bar 200 µm. (d) 3D representation of half of a WM793b human melanoma tumour spheroid on day 17 after formation, additional 3D representations are shown in supplementary material C.3.4. (e) Cell cycle schematic coloured with respect to FUCCI signal. (f) Schematic for Greenspan mathematical model. Nutrient diffuses within the tumour spheroid and is consumed by living cells. (g) Snapshot of nutrient concentration, *c*(*r, t*) for 0 < *r* < *R*_o_(*t*), for a tumour spheroid in phase (iii) and where *R*_o_(*t*) is the tumour spheroids outer radius. External nutrient concentration is *c*_∞_. Inhibited radius, *R*_i_(*t*), and necrotic radius, *R*_n_(*t*), are defined as the radius where the nutrient concentration first reaches the thresholds *c*_i_ and *c*_n_, respectively. (h)-(j) Three experimental designs varying by measurement type. Design 1 considers only the outer radius (green). Design 2 considers the outer (green) and necrotic radius (black). Design 3 considers the outer (green), necrotic (black), and inhibited (magenta) radius. (k) Comparison of experimental designs with respect to their value and experimental effort required. (l)-(n) Three experimental designs that vary due to the time resolution at which measurements are taken. (o) Four experimental designs that vary the number of cells used to form each spheroid.

In this study, we systematically explore a range of experimental designs and measurements. The first and simplest measurements we obtain are of the outer radius of the spheroid. Next, using live-dead cell staining we obtain measurements of the necrotic region. Measurements of the inhibited region are harder to obtain using traditional techniques. We use fluorescent ubiquitination-based cell cycle indicator (FUCCI) transduced cell lines [43–48]. The nuclei of these cells fluoresce red when cells are in the gap 1 (G1) phase of the cell cycle and green when cells are in the synthesis (S), gap 2 (G2) and mitotic (M) phases of the cell cycle (Figure 1e). For clarity, we choose to show cells in the gap 1 (G1) phase in magenta instead of red. These data are collected for human melanoma cell lines established from primary (WM793b) and metastatic cancer sites (WM983b, WM164) [44, 49–51], with endogenously low (WM793b) and high (WM983b, WM164) microphthalmia-associated transcription factor which is a master regulator of melanocyte biology [52]. Analysing these data provides real-time visualisation of the cell cycle throughout tumour spheroids and powerfully reveals the time evolution of the inhibited region (Figure 1a-d). This additional dimension of information that we capture in our experiments, namely the cell cycle status which can be thought of as a measure of time since a freely cycling cell enters the cell cycle, together with the three-spatial dimensions of the tumour spheroid give rise to the term 4D tumour spheroid experiments. Given an abundance of measurements of the outer radius, inhibited radius, and necrotic radius for tumour spheroids across multiple initial spheroid sizes, time points, and cell lines, we calculate maximum likelihood estimates (MLE) and form approximate 95% confidence intervals for the parameters of the Greenspan model. This allows us to quantitatively elucidate how modifying experimental designs can extract more information from experiments. Furthermore, this approach identifies the experimental design choices that are important and lead to reliable biological insight.

## 2 Results

The results in this main document are for spheroids formed with the WM793b human melanoma cell line [44, 49–51]. Additional results in Supplementary Material F and G show results for two other cell lines.

### 2.1 Outer radius measurements are not sufficient to predict inhibited and necrotic radii

Tumour outer radius measurements are simple to obtain and have been used for decades to quantify tumour growth [19, 20]. Modern technology enables these measurements to be obtained more frequently, easily, and accurately. For example, the IncuCyte S3 live cell imaging system (Sartorius, Goettingen, Germany) enables automated image acquisition and processing to measure spheroids every minute throughout an experiment providing a large number of measurements with ease. However, it is unclear whether these measurements provide sufficient information to understand and probe the internal structure of tumour spheroids and accurately predict tumour growth. Furthermore, it is unclear when measurements should be taken and the frequency of measurement. Performing experiments with WM793b spheroids formed with 5000 cells per spheroid, a typical choice in many experiments [7, 11, 52, 53], 24 spheroids are imaged every six hours. We monitor the time evolution of the outer radius to determine when spheroid formation ends and growth begins, which we call day 0 and occurs four days after seeding (Supplementary Material C.1.1), and to decide when to terminate the experiment, which we choose to be day 20. These measurements, supplemented with additional outer radius measurements from spheroids harvested for confocal imaging (Supplementary Material C.2-C.3), provide an abundance of data. We now compare three experimental designs with increasing temporal resolution: (i) Resolution A, using measurements from days 1, 3, 8, 12, 17 (Figure 1l, 2a); (ii) Resolution B, using measurements from days 1, 3, 6, 8, 10, 12, 14, 17, 19 (Figure 1m, 2b); and, (iii) Resolution C, using daily measurements from day 0 to day 19 (Figure 1n, 2c). Excluding the final day(s) of measurements from these temporal resolutions allows a predictive check to be performed. Note that all these temporal resolutions are low relative to the capability of the automated imaging system but are high relative to the number of measurements typically taken in standard experiments [11, 12, 14, 44, 54].

To understand the influence of the choice of temporal resolution we now qualitatively and quantitatively compare the results. Across the three temporal resolutions in Figures 2d-f we observe excellent agreement between the full set of outer radius measurements, collected every six hours, and the predicted outer radius from the Greenspan model simulated with the MLE (Methods 4.1-4.2). However, it is clear that the prediction of the inhibited and necrotic radius is poor with Resolutions A and B (Figures 2d-e). With Resolution C, the prediction of the inhibited and necrotic radius appears to have improved (Figure 2f) but we will show that it is misleading to suggest that increasing the temporal resolution is always beneficial. While MLE point estimates are insightful, it is unclear whether a similarly excellent match to the outer radius measurements could be obtained with different parameter values in the mathematical model. To answer this question we undertake a profile likelihood analysis of the five parameters that govern the behaviour of the mathematical model (Methods 4.1):

**Figure 2:**
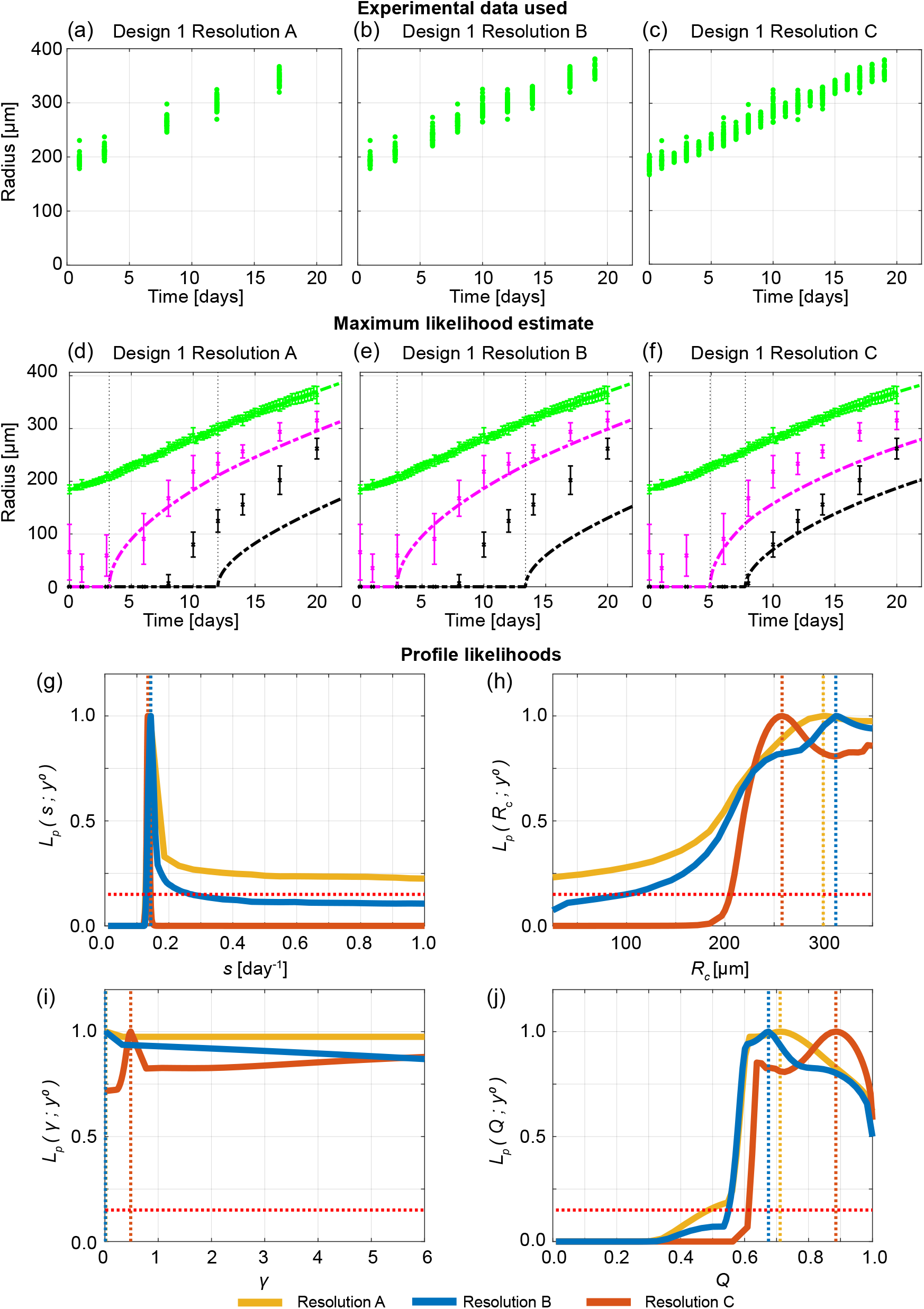
Increasing the temporal resolution when the outer radius is measured is not sufficient to predict necrotic and inhibited radii. (a)-(c) Experimental data used in Design 1 with temporal resolutions A, B, and C. (d)-(f) Comparison of Greenspan model simulated with maximum likelihood estimate compared to full experimental data set, where error bars show standard deviation of the experimental data. Profile likelihoods for (g) *s*, (h) *R*_c_, (i) *γ*, (j) *Q*. Yellow, blue, orange lines in (g)-(j) represent profile likelihoods from Design 1 with temporal resolutions A, B, and C, respectively, and the red-dashed line shows the approximate 95% confidence interval threshold. Results shown for WM793b spheroids formed with 5000 cells per spheroid.

1. *s* [day^−1^], the rate at which cell volume is produced by mitosis per unit volume of living cells (Figure 2g),
2. *R*_c_ [µm], the outer radius when the necrotic region first forms (Figure 2h),
3. *γ* = *λ/s* [-], the proportionality constant given by the rate at which cell volume is lost from the necrotic core, *λ*, divided by the rate at which cell volume is produced by mitosis per unit volume of living cells, *s*, (Figure 2i),
4. *Q*^2^ = (*c*_∞_ − *c*_i_) */* (*c*_∞_ − *c*_n_) [-], the ratio of the difference between the inhibited nutrient concentration threshold, *c*_i_, and external nutrient threshold, *c*_∞_ to the difference between the necrotic nutrient concentration threshold, *c*_n_, and external nutrient threshold, *c*_∞_ (Figure 2j),
5. *R*_o_(0) [µm], the initial outer radius (Supplementary Material D.3).

Profile likelihoods are a powerful tool to visualise and analyse how many parameter values give a similar match to the experimental data in comparison to the MLE. Furthermore, we use profile likelihoods to compute approximate 95% confidence intervals for each parameter (Supplementary Material D.1). Narrow approximate 95% confidence intervals indicate parameters are identifiable and that few parameters give a similar match to the data as the MLE. In contrast, wide approximate 95% confidence intervals suggest that parameters are not identifiable, that many parameters give a similar match to the experimental data, and that additional information is required to confidently estimate the parameters.

The profile likelihoods for *s* across all three temporal resolutions (Figure 2g) lead to a peak that is close to *s* = 0.14 [day^−1^]. These peaks correspond to the MLEs. While there is a wide 95% approximate confidence interval for *s* with Resolution A, there are narrow approximate 95% confidence intervals for *s* with Resolutions B and C. The profile likelihoods for the other parameters, *R*_c_, *γ*, and *Q*, are wide and do not change significantly using different temporal resolutions (Figures 2h-j). For example, the profile likelihoods for *γ* across all three temporal resolutions (Figure 2i) are approximately flat and equal to one. These profile likelihoods for *R*_c_, *γ*, and *Q* suggest that increasing the temporal resolution does not provide significant additional information. These results are consistent with additional results using synthetic data (Supplementary Material E). Additional results for different initial spheroid sizes (Supplementary Material D) and results for the WM983b cell line (Supplementary Material F.1) also clearly show that increasing the temporal resolution while using Design 1 may result in a worse prediction from the MLE for the time evolution of the internal structure. These results do not mean that the mathematical model is incorrect. Our interpretation of these results is that the experimental data are insufficient to correctly identify the parameters in the mathematical model. Overall, these results suggest that Design 1 (Figure 1h) is not a reliable design to identify the true parameter values and cannot be used to determine details of the internal structure of tumour spheroids. This is important because this is the most standard measurement [5, 14–16, 19, 54].

### 2.2 Cell cycle and necrotic core measurements reveal time evolution of internal spheroid structure

Given that measuring the outer radius of tumour spheroids alone (Design 1) is insufficient to determine details of the internal spheroid structure, we now examine which measurements are required to provide reliable estimates. The next simplest measurements to obtain are both the outer radius and necrotic core radius, which we refer to as Design 2 (Figure 1i). However, Design 2 requires far more experimental effort since necrotic core measurements are more time-consuming involving harvesting, fixing, staining procedures, confocal microscopy or cryosectioning, and image processing. In addition, necrotic core measurements are end point measurements only, meaning that many spheroids are required to collect many data points. While intuitively we may anticipate that more experimental effort leads to more insight, it is impossible to quantify the value of this additional effort without a mathematical modelling and uncertainty quantification framework such that we employ here.

Using Design 2 for spheroids formed with 5000 cells per spheroid, we do not observe a necrotic core until approximately day 8 (Figure 3a, Supplementary Material C.3.1). The Greenspan model simulated with the MLE obtained using temporal resolution A, since results obtained using temporal resolutions B and C are very similar (Supplementary Material D.2), excellently matches the growth of the outer radius, as before, and now captures the formation and growth of the necrotic core (Figure 3c). Interestingly, the MLE suggests that the inhibited region is very small, so *R*_i_(*t*) is very close to *R*_n_(*t*). However, experimental measurements of the inhibited radius not only suggest that an inhibited region exists, but that it forms prior to the formation of the necrotic core (Supplementary Material C.3.1). Profile likelihoods for each parameter are relatively narrow, and because the profile for *Q* is peaked and close to *Q* = 1, these profiles are consistent with either the absence of an inhibited region or a very small inhibited region (Figures 3e-h). Therefore, these data do not identify the true parameter values since the calibrated mathematical model is inconsistent with the experimental observations that clearly show the formation of an inhibited region. This inconsistency does not mean that the mathematical model is incorrect. Our interpretation of this inconsistency is that this experimental data are insufficient to identify the parameters in the mathematical model.

**Figure 3:**
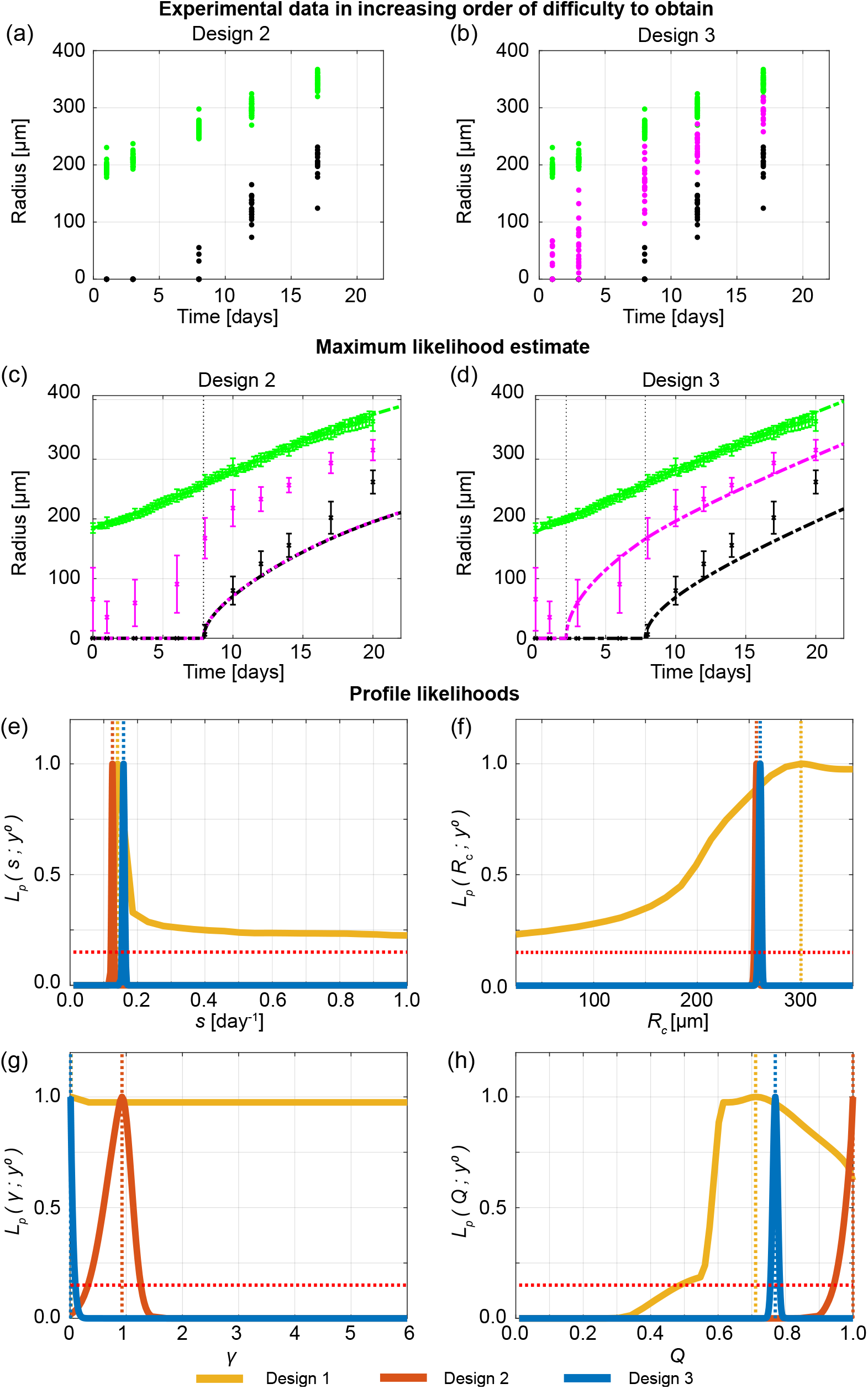
Measuring the necrotic and inhibited radius provides valuable information. (a)-(b) Experimental data used in Designs 2 and 3 with temporal resolution A. (c)-(d) Comparison of Greenspan model simulated with maximum likelihood estimate compared to full experimental data set for Designs 2 and 3, where error bars show standard deviation. Profile likelihoods for (e) *s*, (f) *R*_c_, (g) *γ*, (h) *Q*. Yellow, orange, blue lines in (e)-(h) represent profile likelihoods from Designs 1, 2, and 3, respectively, and the red-dashed line shows the approximate 95% confidence interval threshold. Results shown for WM793b spheroids formed with 5000 cells per spheroid.

We now explore Design 3, where we measure the outer, necrotic, and inhibited radius of multiple tumour spheroids (Figure 1j, 3b). This design is considered third because measuring the inhibited radius is more difficult and requires substantial additional experimental effort. Using FUCCI transduced cell lines in combination with optical clearing procedures and confocal microscopy powerfully reveals intratumoral spatiotemporal differences with respect to the cycle. This method also requires semi-automated image processing and expert guidance to minimise subjectivity and accurately identify the inhibited region boundary (Supplementary Material C.2) [53]. Simulating the Greenspan model with the MLE from Design 3 matches the evolution of the outer radius and captures the evolution of the necrotic and inhibited regions very accurately (Figure 3d). Furthermore, the profile likelihoods for all parameters are well formed, with a single narrow peak, suggesting that Design 3 identifies the true parameter values (Figure 3e-h). Comparing experimental Designs 1, 2, and 3, we observe that the profile likelihoods for *s* are consistent across all designs (Figure 3e) and the profile likelihoods for *R*_c_ (Figure 3f) are consistent for Designs 2 and 3. However, the profile likelihoods for *γ* (Figure 3g) and *Q* (Figure 3h) emphasise the power of measuring the inhibited radius and using Design 3 in comparison to Designs 1 and 2. These observations are consistent with additional results obtained using synthetic data (Supplementary Material E), different cell lines (Supplementary Material F), and initial spheroid sizes (Supplementary Material D.5). In Supplementary Material D.2, we also consider Design 3 with different temporal resolutions and experimental durations. As with Design 2, results obtained using Design 3 with temporal resolutions A, B, and C are very similar. Experiments performed for 4 or 10 days after spheroids form do not accurately predict late time behaviour. Designs that use the first days 10 to 20 or days 16 to 19 of measurements do not always accurately predict early time behaviour. Most insight is gained with Resolutions A, B, and C that cover the full experimental duration.

### 2.3 Information gained using spheroids of different sizes is consistent

In the literature tumour spheroids are initialised with a wide range of cell numbers, leading to inconsistent results that are difficult to meaningfully compare across different protocols [11–18]. Furthermore, it is unclear what the impact of this variability is when tumour spheroids are used to study fine-grained molecular-level interventions or potential drug designs. To quantitatively compare how information gained across experimental designs differs with respect to the initial number of cells in a spheroid we consider four initial spheroid sizes: 1250, 2500, 5000, 10000 cells per spheroid (Figure 1o). To proceed we use Design 3, and measure outer, necrotic, and inhibited radius, with time resolution A. Profile likelihoods for *R*_o_(0) show four distinct narrow peaks corresponding to each initial spheroid size as expected (Figure 4a). Profile likelihoods for *s, R*_c_, and *Q* are consistent across the four initial spheroid sizes, allowing us to compare profile likelihoods on narrower intervals in Figures 4b,c,e. The profile likelihoods for *γ* (Figures 4d) are more variable due to the differing number of measurements collected in phase (iii). These results suggest that the initial spheroid size does not play a significant role in determining information from experiments, provided sufficient measurements are obtained in phase (iii). To support these results, we show along the diagonal of Figure 4f the solution of the mathematical model evaulated at the MLE associated with each initial spheroid size compared to the experimental measurements. Next, on the off-diagonals of Figure 4f, we compare how the Greenspan model simulated with the MLE from one initial spheroid size predicts data from different initial spheroid sizes by only changing the initial radius. For example, in the top right of Figure 4f we show that the Greenspan model simulated with the MLE obtained formed with 10000 cells per spheroid agrees well with data from spheroids formed with 1250 cells per spheroid, when the initial radius is set to be the initial radius of the 1250 MLE. Results in Figure 4f also show the inhibited and necrotic regions form earlier when considering spheroids formed with more cells, and results for spheroids formed with 10000 cells per spheroid suggest that these spheroids form in phase (ii) rather than phase (i). These observations are consistent with additional results from synthetic data (Supplementary Material E) and the WM983b cell line (Supplementary Material F).

**Figure 4:**
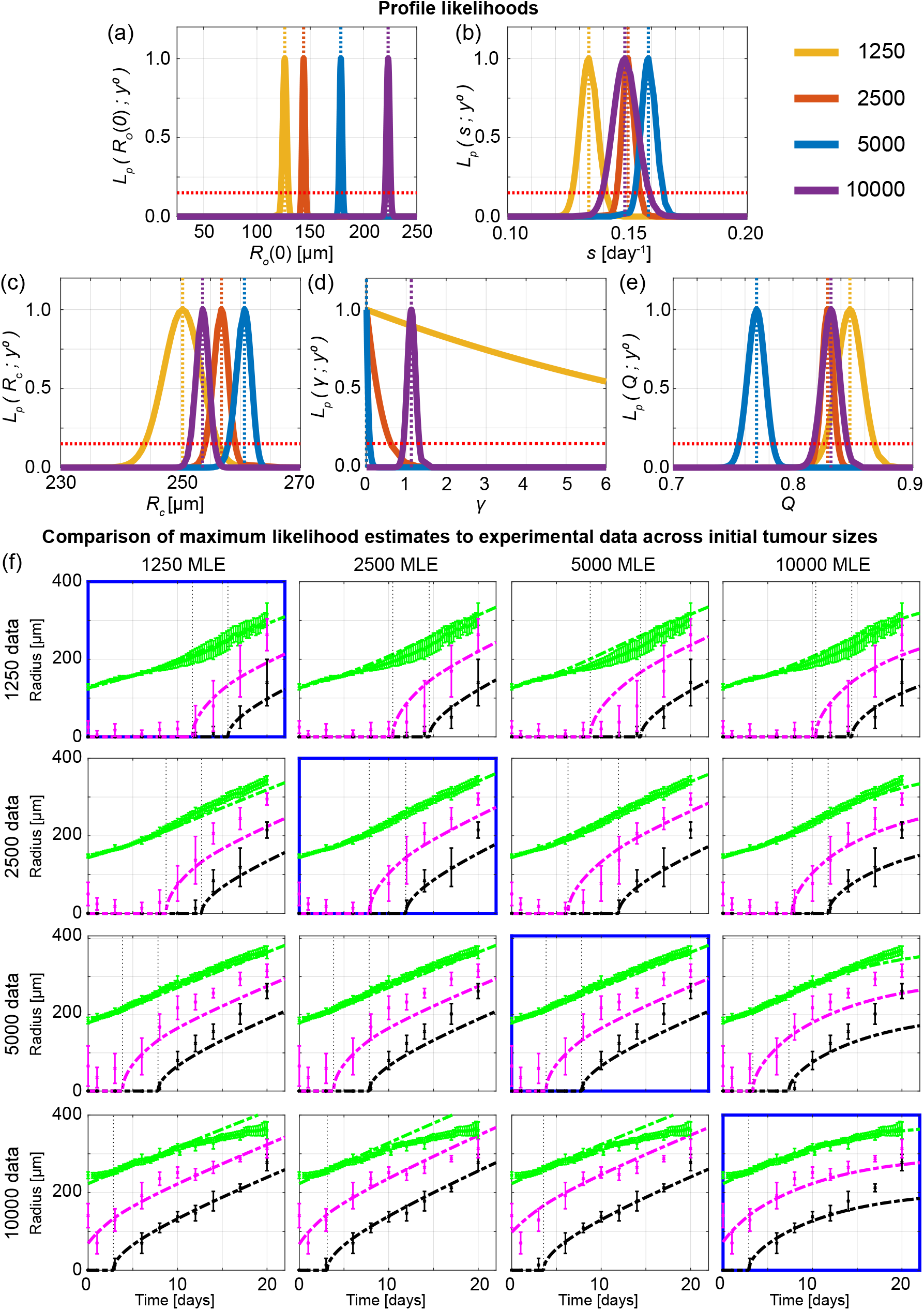
Information gained from experiments across different initial tumour spheroid sizes is mostly consistent. Profile likelihoods for (a) *R*_o_(0), (b) *s*, (c) *R*_c_, (d) *γ*, (e) *Q*. Yellow, orange, blue, and purple lines in (a)-(e) represent profile likelihoods from WM793b spheroids formed with 1250, 2500, 5000, 10000 cells per spheroid, respectively, and the red-dashed line shows the approximate 95% confidence interval threshold. (f) Comparison of Greenspan model simulated with maximum likelihood estimates compared to full experimental data sets across initial tumour spheroid size, where error bars show standard deviation.

## 3 Discussion

In this work we present an objective theoretical framework to quantitatively compare tumour spheroid experiments across a range of experimental designs using the seminal Greenspan mathematical model and statistical profile likelihood analysis. By considering different temporal data resolutions, experiment durations, types of measurements, and initial spheroid sizes we identify the experimental design choices that lead to reliable biological insights and allow us to obtain reliable estimates with low uncertainty of parameters in Greenspan’s model. This approach enables us to quantify the time evolution of the structure of growing spheroids and, since all parameters in Greenspan’s model are physically interpretable and biologically relevant, we also gain insights to the contribution of underlying biological mechanisms.

While it is common in spheroid experiments to measure the outer radius, it is less common to measure the necrotic core and not standard protocol to measure the cell cycle. However, we find that although Design 3, where we obtain outer, necrotic, and inhibited radius measurements, requires most experimental effort it is essential to determine the dynamics of tumour spheroid structure and growth. While we may have anticipated that more experimental effort leads to more insight, it is impossible to quantify the value of this additional effort without a mathematical modelling and uncertainty quantification framework such that we employ here. Alternative experimental designs may also provide useful insights but such approaches likely require more experimental effort and expense than the approach we use here of extending a typical experimental protocol to include necrotic core and cell cycle measurements. We note that should these alternative experimental designs be developed our framework will be relevant to their assessment.

We also show that temporal resolution and initial spheroid sizes are less important experimental design choices. Therefore, we recommend that for future studies, where tumour spheroid structure is important, that cell cycle data are essential and that some measurements using Design 3 are more valuable than many measurements using Designs 1 or 2. Furthermore, as information from tumour spheroids across varying initial spheroid sizes is relatively consistent, provided sufficient measurements in phase (iii) are obtained, we recommend that performing experiments with larger tumour spheroids can be beneficial to obtain useful information in a shorter experimental duration (Supplementary Material E.3). However, we also note that this may lead to large tumour spheroids that begin growth in phase (ii) rather than phase (i).

To perform this analysis we use Greenspan’s seminal mathematical model, where all parameters have a relatively straightforward biological interpretation. We find that Greenspan’s model performs remarkably well across cell lines and initial spheroid sizes, and provides powerful insights into experimental design. Even though Greenspan’s model is relatively simple, and may not capture all of the biological details of tumour spheroid growth, the fact that results for experimental data are consistent with those from synthetic data enhances our confidence that key biological features are captured in Greenspan’s model (Supplementary Material E). Future modelling may wish to explore potential model misspecifications, for example WM983b spheroids appear to reduce in size at very late time suggesting a fourth phase in these *in vitro* experiments (Supplementary Material F); and, WM164 spheroids, possibly due to their lack of spherical symmetry [52], are more challenging to interpret as information gained using spheroids of different sizes is not consistent (Supplementary Material G).

The general framework presented in this work can be applied to other cell types, for example FUCCI-transduced lung, stomach and breast cancer cells [48, 55–57], to extract more information from existing experimental data across experimental designs, and is suitable to be extended to consider tumour spheroids grown in different conditions and to more complex mathematical models. Given that cell cycle data is demonstrated to be informative in this study, we suggest that it may be beneficial for FUCCI technology to be further developed and more widely used, for example in avascular patient-derived organoids [58], and our framework be extended to these heterogeneous populations accordingly. Furthermore, the insights of this study provide a platform for future studies with spheroids that develop, test, and quantify the effectiveness of cancer treatments, possibly across different experimental designs. In such future studies cell cycle data will be informative since cytotoxic or cytostatic drugs may result in similar changes in the outer radius but due to different causes, that can be measured by cell death and cell cycle imaging (Haass laboratory unpublished observations). Therefore, understanding the time evolution of the outer radius, inhibited radius, and necrotic radius will be of great value in such future experiments, which is another advantage and contributing factor of developing a framework with Greenspan’s model in this study.

## 4 Methods

### 4.1 Mathematical model

Greenspan’s mathematical model describes the three phases of avascular tumour spheroid growth [21]. Spherical symmetry is assumed at all times and maintained by adhesion and surface tension. Under these minimal assumptions, the only independent variables are time, *t* [days], and radial position, *r* [µm]. Tumour growth is governed by the evolution of the outer radius, *R*_o_(*t*) [µm], the inhibited radius, *R*_i_(*t*) [µm], and the necrotic radius, *R*_n_(*t*) [µm]. Nutrient diffuses within the spheroid with diffusivity *k* [µm^2^ day^−1^] and is consumed by living cells at a constant rate per unit volume *α* [mol µm^−3^ day^−1^]. The external nutrient concentration is *c*_∞_ [mol µm^−3^]. The nutrient concentration at a distance *r* from the centre of the spheroid and time *t*, denoted *c*(*r, t*) [mol µm^−3^], is assumed to be at diffusive equilibrium. Therefore, at any instant in time we have *c*(*r, t*) = *c*(*r*) due to fast diffusion of nutrient. However, as *R*_o_(*t*) is growing, nutrient diffusion occurs on a growing domain and we write *c*(*r*) = *c*(*r*(*t*)). The inhibited and necrotic regions form when the nutrient concentration at the centre of the spheroid reaches *c*_i_ [mol µm^−3^] and *c*_n_ [mol µm^−3^], respectively. For *c*(*r*(*t*)) > *c*_n_ the rate at which cell volume is produced by mitosis per unit volume of living cells is *s* [day^−1^]. In the necrotic core cellular debris disintegrates into simpler chemical compounds that are freely permeable through cell membranes. The mass lost in the necrotic region is replaced by cells pushed inwards by forces of adhesion and surface tension. The necrotic core loses cell volume at a rate proportional to the necrotic core volume with proportionality constant 3*λ* [day^−1^], where the three is included for mathematical convenience.

Conservation of mass is written in words as *A* = *B* + *C* − *D* − *E* where *A* is the total volume of living cells at any time, *t*; *B* is the initial volume of living cells at time *t* = 0; *C* is the total volume of cells produced in *t* ≥ 0; *D* is the total volume of necrotic debris at time *t*; *E* is the total volume lost in the necrotic core in *t* ≥ 0. Writing *A, B, C, D, E* in their mathematical form gives the conservation of mass equation and also writing the nutrient diffusion equation gives,

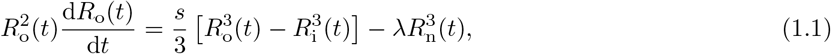

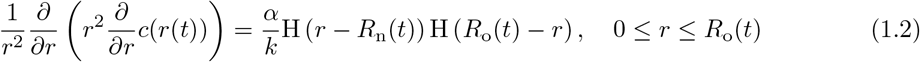

where *R*_i_(*t*), *R*_n_(*t*) are the radii implicitly defined by *c*(*R*_i_(*t*), *t*) = *c*_i_, and *c*(*R*_n_(*t*), *t*) = *c*_n_, respectively, if the nutrient concentration inside the spheroid is sufficiently small otherwise *R*_i_(*t*) = 0 or *R*_n_(*t*) = 0, and H(·) is the Heaviside step function. There are eight unknowns: Θ = (*s, λ, α, k, c*_∞_, *c*_i_, *c*_n_, *R*_o_(0)). Note this includes *R*_o_(0) which we treat as a parameter since we also need to estimate this quantity. Rescaling reduces the number of parameters to five: *θ* = (*R*_o_(0), *R*_c_, *s, γ, Q*). The new dimensionless parameters are: the outer radius when the necrotic region first forms defined as 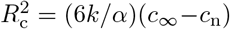; *Q*^2^ = (*c*_∞_ −*c*_i_)*/*(*c*_∞_ −*c*_n_); and *γ* = *s/λ*. Further details, and a formal demonstration that this model is equivalent to a model where nutrient determines the necrotic region and waste produced from live cells determines the inhibited region, are provided in Supplementary Material A.

### 4.2 Practical parameter identifiability analysis

To determine the maximum likelihood estimate (MLE) and approximate 95% confidence intervals for the parameters *θ* = (*R*_o_(0), *R*_c_, *s, γ, Q*) we use profile likelihood identifiability analysis [59–63]. We first choose simple parameter bounds and then compare the width of these simple parameter bounds to realised interval estimates for the parameters. Initial parameter bounds are chosen to be the same across all experimental designs analysed in this study. Outer radius data suggests we choose 0 < *R*_o_(0) < 350 [µm] and 0 < *R*_c_ < 250 [µm]. Assuming a cell doubling time of at least 12 hours and performing preliminary data exploration, we set 0 < *s* < 1 [day^−1^] (Supplementary Material B.1). Limited information exists for the parameter *γ* so bounds are determined by preliminary data exploration to be 0 < *γ* < 6. By definition of *Q* and experimental results that show the inhibited region forms before the necrotic core, we set 0 < *Q* ≤ 1. Note that the time evolution of *R*_o_(*t*) and *R*_n_(*t*) are the same for *Q* = 1 and *Q* > 1. The difference arises for *R*_i_(*t*), where it is equal to *R*_n_(*t*) for *Q* = 1 and equal to zero for *Q* > 1.

To determine the interval estimates for the parameters we treat the mathematical model as having two components. The first is the deterministic mathematical model governing the evolution of *R*_o_(*t*), *R*_n_(*t*), and *R*_i_(*t*) and the second is a probabilistic observation model accounting for experimental variability and measurement error. Specifically, we assume that experimental measurements are noisy observations of the deterministic mathematical model [63, 64]. For each of the three measurement types *R*_o_(*t*), *R*_n_(*t*), and *R*_i_(*t*) we assume that the observation error is independent and identically distributed and that the noise is additive and normally distributed with zero mean and variance 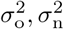 and 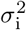, respectively [64, 65] (Supplementary Material B.3). We approximate 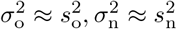, and 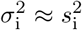 where 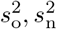, and 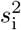 are pooled sample variances of the outer, necrotic, and inhibited radius measurements, respectively [66].

The likelihood function *p*(*y*°|*θ*) is the likelihood of the observations *y*° given the parameter *θ*. This corresponds to the probabilistic observation model evaluated at the observed data. The maximum likelihood estimate is 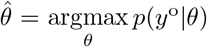. We present results in terms of the normalised likelihood function 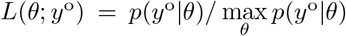 which we consider a function of *θ* for fixed *y*°. Profile likelihoods for each parameter are obtained by assuming the full parameter *θ* can be partitioned into a scalar interest parameter, *ψ*, and vector nuisance parameter, *ϕ*, so that *θ* = (*ψ, ϕ*). The profile likelihood for *ψ* is then 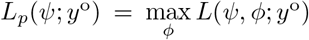. Approximate 95% confidence intervals are then calculated using a profile likelihood threshold value of 0.15 (Supplementary Material D.1) [61]. Prediction intervals are not shown since confidence intervals are narrow in many cases. Further details, including exact forms of the likelihood function and the use of log-likelihoods for calculations, and numerical methods are provided in Supplementary Material B.

### 4.3 Experimental methods

#### Cell culture

The human melanoma cell lines WM793b, WM983b, and WM164 were genotypically characterised [44, 49–51], grown as described in [7], and authenticated by short tandem repeat fingerprinting (QIMR Berghofer Medical Research Institute, Herston, Australia). All cell lines were transduced with fluorescent ubiquitination-based cell cycle indicator (FUCCI) constructs as described in [7, 44].

#### Spheroid generation, culture, and experiments

Spheroids were generated in 96-well cell culture flat-bottomed plates (3599, Corning), with four different seeding densities (1250, 2500, 5000, 10000 total cells/well), using 50 µL total/well non-adherent 1.5% agarose to promote formation of a single spheroid per well [53]. For all spheroid experiments, after a formation phase of 4, 3 and 2 days for WM793b, WM164 and WM983b, respectively (Supplementary Material C.1.1), and then every 3-4 days for the duration of the experiment, 50% of the medium in each well was replaced with fresh medium (200 µL total/well). Incubation and culture conditions were as described in *Cell culture*.

To estimate the outer radius, one plate for each cell line, containing 24 spheroids for each initial spheroid size, was placed inside the IncuCyte S3 live cell imaging system (Sartorius, Goettingen, Germany) incubator (37 °C, 5% CO_2_) immediately after seeding the plates. IncuCyte S3 settings were chosen to image every 6 hours for the duration of the experiment with the 4*×* objective. To estimate the radius of the inhibited and necrotic region and additional outer radius measurements, spheroids maintained in the incubator were harvested, fixed with 4% paraformaldehyde (PFA), and stored in phosphate buffered saline solution, sodium azide (0.02%), Tween-20 (0.1%), and DAPI (1:2500) at 4 °C, on days 3, 4, 5, 7, 10, 12, 14, 16, 18, 21 and 24 after seeding. For necrotic core measurements, 12 hours prior to harvesting 1 µmol total/well DRAQ7™ dye (Abcam, Cambridge, United Kingdom. ab109202) was added to each well [53, 67]. Fixed spheroids were set in place using low melting 2% agarose and optically cleared in 500 µL total/well high refractive index mounting solution (Quadrol 9 % wt/wt, Urea 22 % wt/wt, Sucrose 44 % wt/wt, Triton X-100 0.1 % wt/wt, water) for 2 days in a 24-well glass bottom plate (Cellvis, P24-1.5H-N) before imaging to ensure accurate measurements [68, 69]. Images were then captured using an Olympus FV3000 confocal microscope with the 10*×* objective focused on the equatorial plane of each spheroid.

#### Image processing

Images captured with the IncuCyte S3 were processed using the accompanying IncuCyte 2020C Rev1 software (spheroid analysis type, red image channel, largest red object area per well). Area masks were visually compared with IncuCyte brightfield images to confirm accuracy. Area was converted to an equivalent radius (*r*^2^ = *A/π*). Confocal microscopy images were converted to TIFF files in ImageJ and then processed with custom MATLAB scripts that use standard MATLAB image processing toolbox functions. These scripts are freely available on Zenodo with DOI:10.5281/zenodo.5121093 [70].

#### Statistics and Reproducibility

Details of practical parameter identifiability analysis are presented in Section 4.2. Data points in Figures 2a-c and 3a-b represent measurements of individual spheroids. Data points in Figures 2d-f, 3c-d, 4f represent the mean of the measurements of either the outer radius, inhibited radius, or necrotic radius at that specific time, while error bars represent the standard deviation of those measurements. Supplementary Material C details the number of measurements at each time point for each cell line and experimental data analysed during the study are available on a GitHub repository. We note that some measurements could not be obtained primarily due to blurring of the automated imaging, spheroids not forming properly, or spheroids losing their structural integrity at very late time. Data for these spheroids was excluded. As part of the study we compare results across different experimental designs and determine when our sample size is sufficient. Randomisation and blinding was not possible.

## Data Availability

The datasets generated during and analysed during the current study are available on a GitHub repository and are summarised in the electronic supplementary material.

## Code Availability

Key computer code and all experimental data used to generate computational results are available on a GitHub repository. The computer code for the mathematical modelling and statistical identifiability analysis was written in MATLAB R2021b (v9.11) with the Image Processing Toolbox (v11.4), Optimization Toolbox (v9.2), Global Optimization Toolbox (v4.6), and the Statistics and Machine Learning Toolbox (v12.2).

## Author Contributions

All authors conceived and designed the study. R.J.M. performed the research and drafted the article. R.J.M., A.P.B., and G.G. performed experimental work. All authors provided comments and approved the final version of the manuscript. N.K.H. and M.J.S. contributed equally.

## Acknowledgements

We thank Dr Pascal Buenzli and Dr Patrick Thomas for helpful discussions, John Blake for guidance using IncuCyte, and Dr David Warne for guidance using the high performance computing resource at QUT. This research was carried out at the Translational Research Institute (TRI), Woolloongabba, QLD. TRI is supported by a grant from the Australian Government. We thank the staff in the microscopy core facility at TRI for their outstanding technical support. We thank Prof. Atsushi Miyawaki, RIKEN, Wako-city, Japan, for providing the FUCCI constructs, Prof. Meenhard Herlyn and Ms. Patricia Brafford, The Wistar Institute, Philadelphia, PA, for providing the cell lines.

