## Supplementary Material for "Designing and interpreting 4D tumour spheroid experiments"

### Supplementary Material:
### Designing and interpreting 4D tumour spheroid experiments

Ryan J. Murphy[1], Alexander P. Browning[1], Gency Gunasingh[2], Nikolas K. Haass[2,†], and Matthew J. Simpson[1,†,*]

[1] *Mathematical Sciences, Queensland University of Technology, Brisbane, Australia*

[2] *The University of Queensland Diamantina Institute, The University of Queensland, Brisbane, Australia*

#### Supplementary Material

| Section | Page No. |
|---|---|
| A. Mathematical model | 3 |
| A.1 Nutrient only | 3 |
| A.2 Nutrient and waste | 9 |
| B. Profile likelihood further details | 11 |
| B.1 Numerical method | 11 |
| B.2 Parameter bounds | 14 |
| B.3 Pooled sample variances | 15 |
| C. Experimental data | 16 |
| C.1 Outer radius experimental measurements and images | 16 |
| C.1.1 Spheroid formation duration | 20 |
| C.2 Confocal microscopy | 23 |
| C.1.1 Measurements | 23 |
| C.3 Confocal microscopy supplementary experimental images | 24 |
| C.3.1 WM793b | 24 |
| C.3.2 WM983b | 28 |
| C.3.3 WM164 | 31 |
| C.3.4 3D renderings | 32 |
| D. Additional results for WM793b | 33 |
| D.1 Results in tables | 33 |
| D.2 Measurement times and experimental duration | 35 |
| D.3 Profile likelihoods for $R_o(0)$ | 40 |
| D.4 Outer radius measurements are not sufficient to predict inhibited and necrotic radii | 41 |
| D.5 Cell cycle and necrotic core measurements reveal time evolution of spheroid structure | 45 |
| E. Synthetic data based on WM793b | 49 |
| E.1 Outer radius measurements are not sufficient to predict inhibited and necrotic radii | 49 |
| E.2 Cell cycle and necrotic core measurements reveal time evolution of spheroid structure | 51 |
| E.3 Role of initial spheroid sizes and experimental duration | 52 |
| E.4 Increasing number of measurements | 54 |
| F. Parameter identifiability for WM983b | 56 |
| F.1 Outer radius measurements are not sufficient to predict inhibited and necrotic radii | 57 |
| F.2 Cell cycle and necrotic core measurements reveal time evolution of spheroid structure | 60 |
| F.3 Information gained across spheroid sizes is consistent | 63 |
| G. Parameter identifiability for WM164 | 64 |

### A Mathematical model

#### A.1 Nutrient only

Here we recall Greenspan's mathematical model governing the evolution of $R_o(t)$, $R_i(t)$, and $R_n(t)$ (Figures 1f-g, S1a-c). We consider conservation of mass to govern the evolution of $R_o(t)$. Assuming: i) all living cells are identical and an incompressible mass of constant volume; ii) cell division occurs instantaneously relative to the growth time of the tumour, and each daughter cell occupies the same volume as any other cell; iii) the proliferation rate is a constant, $s$, for cells which have sufficient nutrient; and, iv) the mass density of living cells is constant and equal to density of necrotic debris; then conservation of mass is equivalent to conservation of volume, giving,

$$A = B + C - D - E, \tag{S.1}$$

where $A$ is the total volume of living cells at any time, $t$; $B$ is the initial volume of living cells at time $t = 0$; $C$ is the total volume of cells produced in $t \geq 0$; $D$ is the total volume of necrotic debris at time $t$; and, $E$ is the total volume lost in the necrotic core in $t \geq 0$.

Writing $A, B, C, D, E$ in their mathematical forms and recalling that the volume and surface area

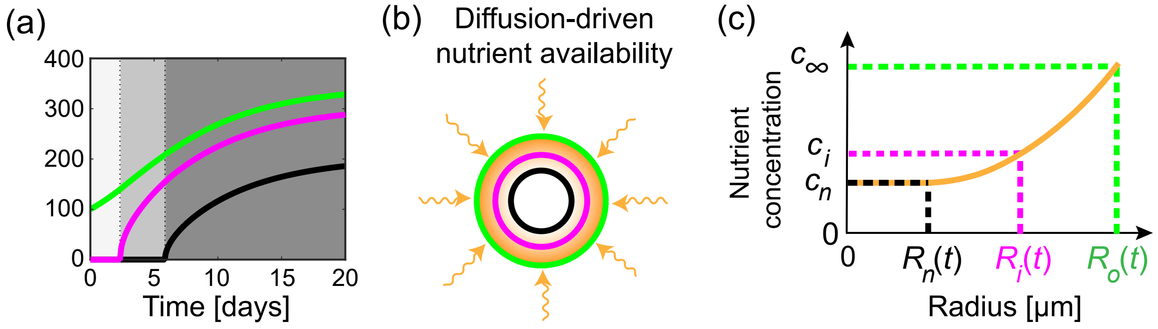


Figure S1: Greenspan's mathematical model. (a) Greenspan's mathematical model describes the three phases of growth and time evolution of the outer radius, $R_o(t)$ (green), inhibited radius, $R_i(t)$ (magenta), and necrotic radius, $R_n(t)$ (black). (b) Schematic for Greenspan mathematical model. Nutrient diffuses within the tumour spheroid and is consumed by living cells. (c) Snapshot of nutrient concentration, $c(r(t))$ for $0 < r < R_o(t)$, for a tumour spheroid in phase (iii). External nutrient concentration is $c_\infty$. Inhibited radius, $R_i(t)$, and necrotic radius, $R_n(t)$, are defined as the radii where the nutrient concentration first reaches the thresholds $c_i$ and $c_n$, respectively.

of a sphere with radius $r$ is $4\pi r^3/3$ and $4\pi r^2$, respectively, gives

$$A = \frac{4\pi}{3}\left(R_\mathrm{o}^3(t) - R_\mathrm{n}^3(t)\right), \tag{S.2.1}$$

$$B = \frac{4\pi}{3}R_\mathrm{o}^3(0), \tag{S.2.2}$$

$$C = 4\pi\int_0^t \int_{R_\mathrm{i}(t)}^{R_\mathrm{o}(t)} sr^2 \ \mathrm{d}r \ \mathrm{d}t, \tag{S.2.3}$$

$$D = \frac{4\pi}{3}R_\mathrm{n}^3(t), \tag{S.2.4}$$

$$E = \frac{4\pi}{3}\int_0^t 3\lambda R_\mathrm{n}^3(t) \ \mathrm{d}t, \tag{S.2.5}$$

where the three inside the integral of equation (S.2.5) is included for mathematical convenience. Substituting equations (S.2.1)-(S.2.5) into equation (S.1) and simplifying gives,

$$R_\mathrm{o}^3(t) = R_\mathrm{o}^3(0) + 3\int_0^t \int_{R_\mathrm{i}(t)}^{R_\mathrm{o}(t)} sr^2 \ \mathrm{d}r \ \mathrm{d}t - \int_0^t 3\lambda R_\mathrm{n}^3(t) \ \mathrm{d}t. \tag{S.3}$$

Differentiating equation (S.3) with respect to time and simplifying gives the more convenient form,

$$R_\mathrm{o}^2(t)\frac{\mathrm{d}R_\mathrm{o}(t)}{\mathrm{d}t} = \underbrace{\frac{s}{3}\left(R_\mathrm{o}^3(t) - R_\mathrm{i}^3(t)\right)}_{\text{proliferation of living cells}} - \underbrace{\lambda R_\mathrm{n}^3(t).}_{\text{mass lost in necrotic core}} \tag{S.4}$$

The other important equation concerns the evolution of nutrient within the spheroid. Rewriting equation (1.2) gives

$$\frac{1}{r^2}\frac{\partial}{\partial r}\left(r^2\frac{\partial}{\partial r}c(r(t))\right) = \frac{\alpha}{k}\mathrm{H}\left(r - R_\mathrm{n}(t)\right)\mathrm{H}\left(R_\mathrm{o}(t) - r\right), \quad 0 \le r \le R_\mathrm{o}(t). \tag{S.5}$$

where $\mathrm{H}(\cdot)$ is the Heaviside step function.

To determine the full evolution of the system we solve equations (S.4) and (S.5) together with the nutrient threshold for inhibition, $c_\mathrm{i}$, and the nutrient threshold for necrosis, $c_\mathrm{n}$, which implicitly define the radius of the inhibited region, $R_\mathrm{i}(t)$, and the radius of the necrotic region, $R_\mathrm{n}(t)$, respectively, through (Figures 1g,

$$c(R_\mathrm{i}(t), t) = c_\mathrm{i}, \tag{S.6.1}$$

$$c(R_\mathrm{n}(t), t) = c_\mathrm{n}, \tag{S.6.2}$$

if the nutrient concentration inside the spheroid is sufficiently small otherwise $R_\mathrm{i}(t) = 0$ or $R_\mathrm{n}(t) = 0$. Note that the equation (S.4) for nutrient does not involve any temporal derivative so the only initial condition required to solve the full system of equations (S.4) and (S.5) is the initial outer radius, $R_\mathrm{o}(0)$.

The solution of equation (S.5) is,

$$c(r(t)) = \begin{cases} c_\infty - \dfrac{\alpha}{6k}\left(R_\mathrm{o}^2(t) - r^2\right) + \dfrac{\alpha R_\mathrm{n}^3(t)}{3k}\left(\dfrac{1}{r} - \dfrac{1}{R_\mathrm{o}(t)}\right), & R_\mathrm{n}(t) \le r \le R_\mathrm{o}(t), \\ c_\mathrm{n}, & 0 \le r \le R_\mathrm{n}(t), \end{cases} \tag{S.7}$$

where

$$c_\infty - c_\mathrm{n} = \frac{\alpha}{3k}\left[\frac{1}{2}\left(R_\mathrm{o}^2(t) - R_\mathrm{n}^2(t)\right) - \frac{R_\mathrm{n}^2(t)}{R_\mathrm{o}(t)}\left(R_\mathrm{o}(t) - R_\mathrm{n}(t)\right)\right]. \tag{S.8}$$

The necrotic region first forms when the nutrient concentration reaches $c_\mathrm{n}$ at the centre, which occurs when $R_\mathrm{n}(t) = 0$ and $r = 0$ in equation (S.8), which gives a critical outer radius,

$$R_\mathrm{c}^2 = \frac{6k}{\alpha}\left(c_\infty - c_\mathrm{n}\right). \tag{S.9}$$

Also recall that $R_\mathrm{i}(t)$ corresponds to $c(R_\mathrm{i}(t), t) = c_\mathrm{i}$ which we can substitute into equation (S.7) to give,

$$c_\infty - c_\mathrm{i} = \frac{\alpha}{3k}\left[\frac{1}{2}\left(R_\mathrm{o}^2(t) - R_\mathrm{i}^2(t)\right) - R_\mathrm{n}^3(t)\left(\frac{1}{R_\mathrm{i}(t)} - \frac{1}{R_\mathrm{o}(t)}\right)\right]. \tag{S.10}$$

Since the inhibited region first forms when the nutrient concentration reaches $c_\mathrm{i}$ at the centre of the spheroid and the necrotic region forms after the inhibited region, setting $R_\mathrm{n}(t) = 0$ and $r = 0$ on right-hand-side of equation (S.10) gives the outer radius when the inhibited region first forms

$$R_\mathrm{d}^2 = \frac{6k}{\alpha}\left(c_\infty - c_\mathrm{i}\right). \tag{S.11}$$

We can then define a useful dimensionless quantity, $Q^2 = R_\mathrm{d}^2/R_\mathrm{c}^2 = \left(c_\infty - c_\mathrm{i}\right)/\left(c_\infty - c_\mathrm{n}\right)$, which is related to the time when phase (ii) begins.

Equations (S.4), (S.5), and (S.6) can now be solved in each of phase (i), (ii), and (iii). To provide valuable insights into the structure of the solutions to the Greenspan model it helps to consider the non-dimensional form of the equations and their solutions. To non-dimensionalise we rescale time with $s$ to give $\tau = st$ and rescale lengths with $R_\mathrm{c}$ via $\xi_\mathrm{o}(t) = R_\mathrm{o}(t)/R_\mathrm{c}$, $\xi_\mathrm{i}(t) = R_\mathrm{i}(t)/R_\mathrm{c}$, and $\xi_\mathrm{n}(t) = R_\mathrm{n}(t)/R_\mathrm{c}$. Then phase (ii) starts when $\xi_\mathrm{o}(t) = Q$ and phase (iii) starts when $\xi_\mathrm{o}(t) = 1$. We now consider each phase in turn.

##### Phase (i)

In phase (i), all cells are free to proliferate and the nutrient concentration is sufficiently high, i.e. $c(r,t) > c_\mathrm{i}$ for $0 \le r \le R_\mathrm{o}(t)$, such that there is no inhibited or necrotic region (Figure 1(a)). Phase (i) ends when the nutrient concentration at the centre of the spheroid equals the inhibited threshold, when $c(0,t) = c_\mathrm{i}$ and $R_\mathrm{o}(t) = R_\mathrm{d}$.

Since $R_\mathrm{i}(t) = 0$ and $R_\mathrm{n}(t) = 0$, equation (S.4) becomes

$$R_\mathrm{o}^2(t)\frac{\mathrm{d}R_\mathrm{o}(t)}{\mathrm{d}t} = \frac{s}{3}R_\mathrm{o}^3(t), \tag{S.12}$$

giving,

$$R_\mathrm{o}(t) = R_\mathrm{o}(0)\exp\left(\frac{st}{3}\right). \tag{S.13}$$

Non-dimensionalising gives,

$$\xi_\mathrm{o}(\tau) = \xi_\mathrm{o}(0)\exp\left(\frac{\tau}{3}\right), \quad \text{for} \quad 0 \leq \tau \leq 3\log\left(\frac{Q}{\xi_\mathrm{o}(0)}\right) = \tau_1. \tag{S.14}$$

Given the solution in equation (S.14) we determine $R_\mathrm{o}(t)$ by reintroducing $s$ and $R_\mathrm{c}$,

$$R_\mathrm{o}(t) = \xi_\mathrm{o}(st)R_\mathrm{c}, \quad \text{for} \quad 0 \leq t \leq \frac{\tau_1}{s}. \tag{S.15}$$

Note that $R_\mathrm{i}(t) = 0$ and $R_\mathrm{n}(t) = 0$ throughout phase (i). Hence, we have obtained $R_\mathrm{o}(t), R_\mathrm{i}(t), R_\mathrm{n}(t)$ throughout this phase.

**Phase (ii)**

In phase (ii) the spheroid experiences inhibited growth due to a core of inhibited cells and outer region of freely proliferating cells (Figure 1(b)). Phase (ii) ends when the necrotic core forms. Since $R_\mathrm{i}(t) > 0$ and $R_\mathrm{n}(t) = 0$, equation (S.4) becomes

$$R_\mathrm{o}^2(t)\frac{\mathrm{d}R_\mathrm{o}(t)}{\mathrm{d}t} = \frac{s}{3}\left(R_\mathrm{o}^3(t) - R_\mathrm{i}^3(t)\right). \tag{S.16}$$

Non-dimensionalising equation (S.16) gives,

$$\xi_\mathrm{o}^2(\tau)\frac{\mathrm{d}\xi_\mathrm{o}(\tau)}{\mathrm{d}\tau} = \frac{1}{3}\left[\xi_\mathrm{o}^3(\tau) - \xi_\mathrm{i}^3(\tau)\right]. \tag{S.17}$$

Equation (S.17) is a function of two variables, $\xi_\mathrm{o}(\tau)$ and $\xi_\mathrm{i}(\tau)$, which we can simplify to a function of one variable by introducing a change of variables $y_\mathrm{i}(\tau) = \xi_\mathrm{i}(\tau)/\xi_\mathrm{o}(\tau)$, and by using the constraint $Q^2/\xi_\mathrm{o}^2(\tau) = 1 - y_\mathrm{i}^2(\tau)$, to give

$$\frac{3y_\mathrm{i}(\tau)}{\left(1 - y_\mathrm{i}(\tau)^2\right)\left(1 - y_\mathrm{i}(\tau)^3\right)}\frac{\mathrm{d}y_\mathrm{i}(\tau)}{\mathrm{d}\tau} = 1, \tag{S.18}$$

with initial condition $y_\mathrm{i}(\tau) = 0$ at $\tau = \tau_1$ and terminating condition $y_\mathrm{i}(\tau)^2 = 1 - Q^2$. The constraint used to derive equation (S.18) and the termination condition for phase (ii) are obtained with the following argument. In phase (ii) equation (S.10) is

$$R_{\rm o}^2(t) - R_{\rm i}^2(t) = R_{\rm d}^2. \tag{S.19}$$

Non-dimensionalising equations (S.8) and (S.19), using definitions of $\xi_{\rm o}(\tau), \xi_{\rm i}(\tau)$, $Q$, and combining the resulting expressions gives $Q^2 = \xi_{\rm o}^2(\tau) - \xi_{\rm i}^2(\tau)$. Rewriting in terms of $y_{\rm i}(\tau)$ gives $Q^2/\xi_{\rm o}^2(\tau) = 1 - y_{\rm i}^2(\tau)$, which gives the constraint used to derive equation (S.18). Using the fact that phase (ii) ends when $\xi_{\rm o}(\tau) = 1$ and rearranging gives the termination condition for $y_{\rm i}(\tau)$.

Numerically solving equation (S.18), using MATLAB's in-built *ode15s* differential equation solver [1] with absolute and relative tolerances set to $1 \times 10^{-8}$, we obtain $y_{\rm i}(\tau)$ for phase (ii). To obtain $R_{\rm o}(t)$ we use the constraint $Q^2/\xi_{\rm o}^2(\tau) = 1 - y_{\rm i}(\tau)^2$, and definitions of $\xi_{\rm o}(\tau)$ and $\xi_{\rm i}(\tau)$ to obtain $R_{\rm o}(t) = R_{\rm c}Q\left[1 - y_{\rm i}^2(st)\right]^{-1/2}$. Similarly using the constraints we obtain $R_{\rm i}(t) = R_{\rm c}Q\left[1/\left(1 - y_{\rm i}^2(st)\right) - 1\right]^{1/2}$. Recall $R_{\rm n}(t) = 0$ throughout phase (ii). Hence, we have obtained $R_{\rm o}(t), R_{\rm i}(t), R_{\rm n}(t)$ throughout this phase.

**Phase (iii)**

In phase (iii) the spheroid experiences inhibited growth due to an outer proliferating region, an intermediate region of inhibited cells, and a necrotic core (Figure 1(a)). At steady state there is a balance between the number of cells that are proliferating in the outer region and mass lost from the necrotic core.

Since $R_{\rm i}(t) > 0$ and $R_{\rm n}(t) > 0$, all terms in equation (S.4) are non-zero. Non-dimensionalising equation (S.4) gives

$$\xi_{\rm o}^2(\tau)\frac{{\rm d}\xi_{\rm o}(\tau)}{{\rm d}\tau} = \frac{1}{3}\left[\xi_{\rm o}^3(\tau) - \xi_{\rm i}^3(\tau)\right] - \gamma\xi_{\rm n}^3(\tau), \tag{S.20}$$

where $\gamma = \lambda/s$. Equation (S.20) is a function of three variables $\xi_{\rm o}(\tau), \xi_{\rm i}(\tau), \xi_{\rm n}(\tau)$. Introducing $y_{\rm i}(\tau) = \xi_{\rm i}(\tau)/\xi_{\rm o}(\tau)$ and $y_{\rm n}(\tau) = \xi_{\rm n}(\tau)/\xi_{\rm o}(\tau)$ we rewrite equation (S.20) and the non-dimensionalised forms of equations (S.8) and (S.10) as

$$\frac{9y_{\rm n}(\tau)}{(1+2y_{\rm n}(\tau))\,(1-y_{\rm n}(\tau))}\frac{{\rm d}y_{\rm n}(\tau)}{{\rm d}\tau} = 1 - y_{\rm i}^3(\tau) - 3\gamma y_{\rm n}^3(\tau), \tag{S.21.1}$$

$$\xi_{\rm o}^{-2}(\tau) = (1 - y_{\rm n}(\tau))^2\,(1 + 2y_{\rm n}(\tau)), \tag{S.21.2}$$

$$\frac{Q^2}{\xi_{\rm o}^2(\tau)} = 1 - y_{\rm i}^2(\tau) - 2y_{\rm n}^3(\tau)\left(\frac{1 - y_{\rm i}(\tau)}{y_{\rm i}(\tau)}\right), \tag{S.21.3}$$

noting that equation (S.21.1) is obtained using equation (S.21.2). Then we numerically solve equations (S.21.1)-(S.21.3) to obtain $y_{\rm i}(\tau)$ and $y_{\rm n}(\tau)$ using the the following approach. First, we substitute equation (S.21.2) into equation (S.21.3) to eliminate $\xi_{\rm o}(\tau)$ and rearrange which gives

$$0 = -Q^2\left[(1 - y_{\rm n}(\tau))^2\,(1 + 2y_{\rm n}(\tau))\right] + 1 - y_{\rm i}^2(\tau) - 2y_{\rm n}^3(\tau)\left(\frac{1 - y_{\rm i}(\tau)}{y_{\rm i}(\tau)}\right). \tag{S.22}$$

Equations (S.21.1) and (S.22) form a system of differential-algebraic equations which we numerically solve using MATLAB's in-built *ode15s* solver with relative and absolute tolerances set to $1 \times 10^{-8}$. Given the solution for $y_{\mathrm{n}}(\tau)$ and $y_{\mathrm{i}}(\tau)$ we obtain $R_{\mathrm{o}}(t), R_{\mathrm{i}}(t)$ and $R_{\mathrm{n}}(t)$ using the following approach. Given $y_{\mathrm{n}}(\tau)$ we obtain $\xi_{\mathrm{o}}(\tau)$ using equation (S.21.2). Then $R_{\mathrm{o}}(t) = R_{\mathrm{c}}\xi_{\mathrm{o}}(st)$. Using the definition of $y_{\mathrm{i}}(\tau)$, $y_{\mathrm{n}}(\tau)$ and $\xi_{\mathrm{o}}(\tau)$ we obtain $R_{\mathrm{i}}(t) = R_{\mathrm{c}} y_{\mathrm{i}}(st)\xi_{\mathrm{o}}(st)$ and $R_{\mathrm{n}}(t) = R_{\mathrm{c}} y_{\mathrm{n}}(st)\xi_{\mathrm{o}}(st)$. Hence, we have obtained $R_{\mathrm{o}}(t), R_{\mathrm{i}}(t), R_{\mathrm{n}}(t)$ throughout this phase. Key software, including precise details of the implementation of this section, is freely available on a GitHub repository.

Greenspan's mathematical model assumes that tumour spheroids experience three phases of growth [2]. While we find experimental evidence confirming that many tumour spheroids experience three phases of growth (Figures 1, S10, S11, S12, S14, and S15), we also find experimental evidence suggesting tumour spheroids seeded with a higher number of cells may form in phase (ii) (Figures S13 and S16). Here, we now describe how to initialise Greenspan's mathematical model in phase (ii) and in phase (iii). While we do not experimentally observe spheroids forming in phase (iii), here we include how to initialise Greenspan's model in phase (iii) for completeness and since calculations used for statistical identifiability analysis (Supplementary Material B.1) may evaluate the likelihood of starting in phase (iii).

To initialise Greenspan's model with a spheroid in phase (ii) we first prescribe $R_{\mathrm{o}}(0)$ and recall that in phase (ii) there is no necrotic core, so $R_{\mathrm{n}}(0) = 0$. Then from equation (S.10) with $R_{\mathrm{n}}(0) = 0$, the corresponding inhibited radius is $R_{\mathrm{i}}(0) = \left(R_{\mathrm{o}}(0)^2 - Q^2 R_{\mathrm{c}}^2\right)^{1/2}$. To initialise Greenspan's model with a spheroid in phase (iii) we first prescribe $R_{\mathrm{o}}(0)$. Given $R_{\mathrm{o}}(0)$ we rewrite equation (S.8) as the following cubic polynomial $(2/R_{\mathrm{o}}(0))R_{\mathrm{n}}(0)^3 - 3R_{\mathrm{n}}(0)^2 + R_{\mathrm{o}}(0)^2 - R_{\mathrm{c}}^2 = 0$, where $R_{\mathrm{n}}(0)$ is the unknown variable. We determine the three solutions of this polynomial using the MATLAB function *roots* [3] and define $R_{\mathrm{n}}(0)$ as the only physically realistic real-valued solution which satisfies $0 < R_{\mathrm{n}}(0) < R_{\mathrm{o}}(0)$. Similarly, to obtain $R_{\mathrm{i}}(0)$, we rewrite equation (S.10) as the following cubic polynomial $R_{\mathrm{i}}(0)^3 + (Q^2R_{\mathrm{c}}^2 - R_{\mathrm{o}}(0)^2 - 2R_{\mathrm{n}}(0)^3/R_{\mathrm{o}}(0))R_{\mathrm{i}}(0) + 2R_{\mathrm{n}}(0)^3 = 0$, where $R_{\mathrm{i}}(0)$ is the unknown variable and $R_{\mathrm{o}}(0)$ and $R_{\mathrm{n}}(0)$ are known. We then define $R_{\mathrm{i}}(0)$ as the only physically realistic real-valued solution which satisfies $R_{\mathrm{n}}(0) < R_{\mathrm{i}}(0) < R_{\mathrm{o}}(0)$. For statistical identifiability analysis we assume spheroids may form in phase (i), phase (ii), or phase (iii).

The approach taken in phases (i), (ii), and (iii) means that we do not require knowledge of the values of the parameters $c_\infty, c_{\mathrm{n}}, c_{\mathrm{i}}$, $k$ and $\alpha$ but instead only the value of $Q = [(c_\infty - c_{\mathrm{i}})/(c_\infty - c_{\mathrm{n}})]^{1/2}$. This reduces the number of parameters describing the evolution of the spheroid from eight to five. The three pieces of information no longer consider regard the nutrient concentration which we do not directly measure in this study and has been explored in other studies [4, 5]. Furthermore, equations (S.8)-(S.10) show that there are constraints on the relationships between $R_{\mathrm{o}}(t)$, $R_{\mathrm{i}}(t)$, $R_{\mathrm{n}}(t)$ which can be explored further.

#### A.2 Nutrient and waste

The model presented in methods section 4.1 and supplementary material A.1 is a special case of Greenspan's model [2]. The general Greenspan model proposes the inhibited region is a result of a build up of waste produced from live or dead cells and the necrotic region forms due to a lack of nutrient. Here, we consider the alternative case where waste is produced from live cells only and show that, for the measurements we obtain, it is equivalent to the nutrient only case we consider in the main manuscript (Figure S2). We do not consider waste produced only from dead cells in this study since with that model the necrotic core must form before the inhibited region which is not what we observe in these experiments (Figure 1(b)).

In comparison to the nutrient only model in supplementary material A.1, the model with nutrient and waste requires an additional equation for the evolution of waste concentration, $\beta(r,t)$. The full system of governing equations are,

$$R_{\mathrm{o}}^2(t)\frac{\mathrm{d}R_{\mathrm{o}}(t)}{\mathrm{d}t} = \frac{s}{3}\left[R_{\mathrm{o}}^3(t) - \max(R_{\mathrm{n}}^3(t), R_{\mathrm{i}}^3(t))\right] - \lambda R_{\mathrm{n}}^3(t), \tag{S.23.1}$$

$$\frac{1}{r^2}\frac{\partial}{\partial r}\left(r^2\frac{\partial}{\partial r}c(r(t))\right) = \frac{\alpha}{k}H\left(r - R_{\mathrm{n}}(t)\right)H\left(R_{\mathrm{o}}(t) - r\right), \quad 0 \leq r \leq R_{\mathrm{o}}(t), \tag{S.23.2}$$

$$\frac{1}{r^2}\frac{\partial}{\partial r}\left(r^2\frac{\partial}{\partial r}\beta(r(t))\right) = -\frac{P}{\kappa}H\left(r - R_{\mathrm{n}}(t)\right)H\left(R_{\mathrm{o}}(t) - r\right), \quad 0 \leq r \leq R_{\mathrm{o}}(t), \tag{S.23.3}$$

where equations (S.23.1) and (S.23.2) are unchanged, by restricting our attention to the case when the inhibited region forms before the necrotic region, and equation (S.23.3) is new. In equation (S.23.3), the term on the right-hand-side corresponds to production of waste by live cells at a constant rate per unit volume $P$ that diffuses with diffusivity $\kappa$. Furthermore, $R_{\mathrm{i}}(t)$ is defined as the solution of $\beta(R_{\mathrm{i}}(t),t) = \beta_{\mathrm{i}}$ if a solution exists and $R_{\mathrm{i}}(t) = 0$ otherwise.

This model, with nutrient and waste, is equivalent to the nutrient only model when we focus on the five key parameters $R_{\mathrm{o}}(0), R_{\mathrm{c}}, s, \gamma, Q$ governing the dynamics. The only difference is a new definition of $Q$,

$$Q^2 = \frac{\beta_{\mathrm{i}}\kappa}{P} \times \frac{\alpha}{k(c_\infty - c_{\mathrm{n}})}, \tag{S.24}$$

where $\beta_{\mathrm{i}}\kappa/P$ are the parameters related to nutrient and $\alpha/k(c_\infty - c_{\mathrm{n}})$ are the parameters related to waste. This new definition of $Q$ provides a different interpretation of the data since $Q$ now represents a combination of waste and nutrient parameters. Importantly, with this new definition of $Q$ there are two cases to consider: i) $Q \leq 1$, and ii) $Q > 1$. Previously, we only considered case (i). In case (ii) where $Q > 1$ the necrotic core forms before the inhibited region. We do not observe this scenario in the experiments that we perform and therefore we restrict the attention of this study to case (i) with $Q \leq 1$.

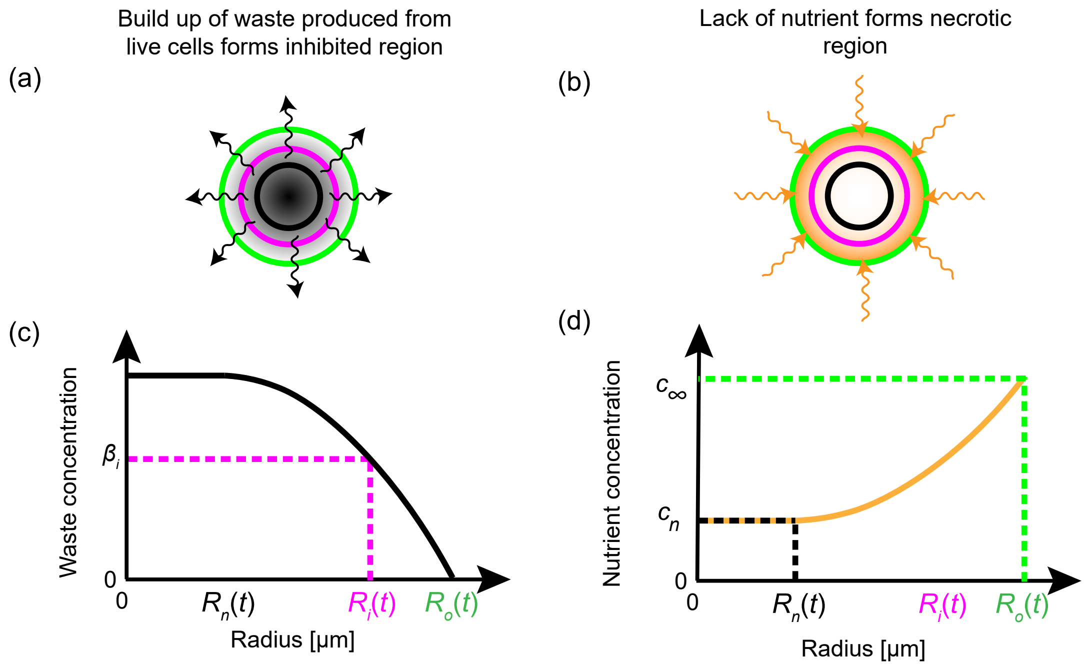


Figure S2: Greenspan's model with waste and nutrient. (a),(c) Build up of waste from live cells results in the formation of an inhibited region. (a) Schematic of waste produced from live cells and diffusing to the external environment. (c) Snapshot of waste concentration against spheroid radius for a spheroid in phase (iii). $R_\mathrm{i}(t)$ is determined by the inhibited waste threshold $\beta_\mathrm{i}$. (b),(d) Lack of nutrient forms the necrotic region. (b) Nutrient, shown in orange, diffusing into the spheroid. (d) Snapshot of nutrient concentration against spheroid radius for a spheroid in phase (iii). $R_\mathrm{n}(t)$ is determined by the necrotic nutrient threshold $c_\mathrm{n}$. External nutrient concentration is $c_\infty$.

### B Profile likelihood further details

#### B.1 Numerical method

Parameter identifiability using statistical profile likelihood analysis is outlined in the methods section 4.2. We now provide further details.

We partition the full set of observations $y^{\mathrm{o}}$ into sets of observations $y^{\mathrm{o}}_{\mathrm{o}}, y^{\mathrm{o}}_{\mathrm{n}}$, and $y^{\mathrm{o}}_{\mathrm{i}}$ corresponding to experimental measurements of $R_{\mathrm{o}}(t), R_{\mathrm{n}}(t)$, and $R_{\mathrm{i}}(t)$. For computational accuracy, we perform calculations using the log-likelihood which is, assuming data independence,

$$\begin{aligned} l\left(\theta; y^{\mathrm{o}}\right) = &\sum_{j=1}^{N_{\mathrm{o}}} \log\left[f(y^{\mathrm{o}}_{\mathrm{o},j}; y_{\mathrm{o},j}(\theta), \sigma^2_{\mathrm{o}})\right] + \sum_{j=1}^{N_{\mathrm{n}}} \log\left[f(y^{\mathrm{o}}_{\mathrm{n},j}; y_{\mathrm{n},j}(\theta), \sigma^2_{\mathrm{n}})\right] \\ &+ \sum_{j=1}^{N_{\mathrm{i}}} \log\left[f(y^{\mathrm{o}}_{\mathrm{i},j}; y_{\mathrm{i},j}(\theta), \sigma^2_{\mathrm{i}})\right], \end{aligned} \tag{S.25}$$

where $y_{\mathrm{o},j}(\theta), y_{\mathrm{n},j}(\theta)$, and $y_{\mathrm{i},j}(\theta)$ are values of $R_{\mathrm{o}}(t), R_{\mathrm{n}}(t)$, and $R_{\mathrm{i}}(t)$ generated from Greenspan's deterministic mathematical model and evaluated at time points corresponding to the experimental observations $y^{\mathrm{o}}_{\mathrm{o},j}, y^{\mathrm{o}}_{\mathrm{n},j}$, and $y^{\mathrm{o}}_{\mathrm{i},j}$, respectively; $f(x; \mu, \sigma^2)$ denotes a Gaussian probability density function with mean $\mu$ and variance $\sigma^2$, calculated using MATLAB's `normpdf` function [6]; $N_{\mathrm{o}}, N_{\mathrm{n}}$, and $N_{\mathrm{i}}$ denote the total number of experimental observations of $R_{\mathrm{o}}(t), R_{\mathrm{n}}(t)$, and $R_{\mathrm{i}}(t)$, respectively; and, $\sigma^2_{\mathrm{o}}, \sigma^2_{\mathrm{n}}$, and $\sigma^2_{\mathrm{i}}$ correspond to pooled variances of the three measurement types $R_{\mathrm{o}}(t), R_{\mathrm{n}}(t)$, and $R_{\mathrm{i}}(t)$, respectively [7,8]. We approximate $\sigma^2_{\mathrm{o}} \approx s^2_{\mathrm{o}}, \sigma^2_{\mathrm{n}} \approx s^2_{\mathrm{n}}$, and $\sigma^2_{\mathrm{i}} \approx s^2_{\mathrm{i}}$, where $s^2_{\mathrm{o}}, s^2_{\mathrm{n}}, s^2_{\mathrm{i}}$ are pooled sample variances of the outer, necrotic, and inhibited radius measurements, respectively [9]. The pooled sample variance for the outer radius is defined as

$$s^2_{\mathrm{o}} = \frac{1}{N_{\mathrm{o}} - 1} \sum_{j=1}^{N_{\mathrm{o}}} \left(y^{\mathrm{o}}_{\mathrm{o},j} - \overline{y^{\mathrm{o}}_{\mathrm{o},j}}\right)^2, \tag{S.26}$$

where $y^{\mathrm{o}}_{\mathrm{o},j}$ is the $j^{\mathrm{th}}$ observation in $y^{\mathrm{o}}_{\mathrm{o}}$ and $\overline{y^{\mathrm{o}}_{\mathrm{o},j}}$ is the sample mean of $y^{\mathrm{o}}_{\mathrm{o}}$ corresponding to the time at which the $j^{\mathrm{th}}$ measurement was observed. We define $s^2_{\mathrm{n}}$ and $s^2_{\mathrm{i}}$ similarly.

The maximum likelihood esimate (MLE), $\hat{\theta}$, is defined as,

$$\hat{\theta} = \underset{\theta}{\operatorname{argmax}} \left[l\left(\theta; y^0\right)\right], \tag{S.27}$$

which we determine by numerically solving the equivalent minimisation problem,

$$\hat{\theta} = \underset{\theta}{\operatorname{argmin}} \left[-l\left(\theta; y^0\right)\right]. \tag{S.28}$$

By assuming the full parameter $\theta$ can be partitioned into an interest scalar parameter, $\psi$, and a nuisance vector parameter, $\phi$, the profile log-likeilihood is

$$l_p\left(\psi; y^0\right) = \max_{\phi}\Bigg[\sum_{j=1}^{N_\mathrm{o}} \log\left[f(y^\mathrm{o}_{\mathrm{o},j}; y_{\mathrm{o},j}(\psi,\phi), \sigma_\mathrm{o}^2)\right] + \sum_{j=1}^{N_\mathrm{n}} \log\left[f(y^\mathrm{o}_{\mathrm{n},j}; y_{\mathrm{n},j}(\psi,\phi), \sigma_\mathrm{n}^2)\right] + \sum_{j=1}^{N_\mathrm{i}} \log\left[f(y^\mathrm{o}_{\mathrm{i},j}; y_{\mathrm{i},j}(\psi,\phi), \sigma_\mathrm{i}^2)\right]\Bigg]. \tag{S.29}$$

Given the five-dimensional parameter space that we are searching to find the maximum likelihood estimate and the four-dimensional parameter space we search to find profile likelihoods, we sequentially determine the maximum likelihood estimate (MLE) and profile likelihoods. All subsequent minimisation optimisations are performed using functions in MATLABs global optimisation toolbox. Specifically, we use the *GlobalSearch* function [10] where we create the following optimisation problem structure. We set the local solver to be the *fmincon* function using the sequential quadratic programming (`sqp`) algorithm, `MaxIterations` = 2500 and `MaxFunctionEvaluations` = 5000. The objective function is defined as the argument of the minimisation of the right hand side of equation (S.28). Other non-default settings that we vary, include `NumTrialPoints`, `MaxTime`, `FirstGuess`, `lowerbounds`, `upperbounds`, along with the method we use to find the MLE and approximate 95% confidence intervals are now discussed.

1. Firstly, we search for MLE. We set the `lowerbounds` and `upperbounds` in agreement with the simple parameter bounds defined in the methods section 4.2. By setting `NumTrialPoints` = 5000 and `MaxTime` = 7200 [seconds], we search for the maximum likelihood estimate for 2 hours with the `FirstGuess` as $(Q, \gamma, s, R_\mathrm{c}, R_\mathrm{o}(0)) = (0.9, 3, 0.5, 175, 125)$. This gives the first estimate for the maximum likelihood estimate $\hat{\theta}_1$. However, numerical experimentation indicates this first estimate is not always an accurate estimate of the true MLE.

2. Secondly, we partition the simple parameter bounds into two sets: [`lowerbounds`, $\hat{\theta}_1$] which we refer to as the lower set, and [$\hat{\theta}_1$, `upperbounds`] which we refer to as the upper set. We then discretise each lower and upper set uniformly using 20 grid points, including the end points. Starting at the grid point associated with $\hat{\theta}_1$ we set `FirstGuess` = $\hat{\theta}_1$, `NumTrialPoints` = 2000 and `MaxTime` = 900 [seconds] in the *GlobalSearch* function. We then move to the next closest grid point and adjust `FirstGuess`. If we are at the closest grid point to $\hat{\theta}_1$ we set `FirstGuess` to be the solution at the previous gridpoint. If we are at any other grid point we make a first order approximation of the first guess by linear extrapolation of the values obtained from the two previous grid points. Before using the first order approximation as a first guess we also check that the value remains within the parameter bounds and if it does not we set `FirstGuess` to be the solution at the previous gridpoint. After calculating the likelihood at each point in the lower and upper set we combine these together to form the first approximation for the profile likelihood.

3. Thirdly, we calculate an estimate of the confidence intervals using a profile likelihood threshold

value of 0.15, which can be approximately calibrated via simulation or the $\chi^2-$distribution [11]. Specifically, we start at either end of the simple parameter bounds until we determine the first grid point where the normalised profile likelihood, $L_p\left(\psi; y^0\right) = \exp\left(l_p\left(\psi; y^0\right) - \max_\theta\left[l\left(\theta; y^0\right)\right]\right)$, is greater than 0.15. We then set new lower and upper bounds as being two grid points to the left or right of that location, respectively. Note that a more sophisticated approach to determine the approximate 95% confidence intervals is applied in step seven to compute the results shown in Table S3, which is not required here.

4. Fourth, we repeat the search for the maximum likelihood estimate using the new lower and upper bounds with the same settings as we first used.

5. Fifth, we repeat the calculations for the profile likelihoods using the new lower and upper bounds.

6. Sixth, we determine the maximum likelihood estimate to be the value across all calculations which maximises the likelihood. We form the final profile likelihood from steps two and four and present the normalised likelihood function, $L_p\left(\psi; y^0\right) = \exp\left(l_p\left(\psi; y^0\right) - \max_\theta\left[l\left(\theta; y^0\right)\right]\right)$, in figures.

7. To compute approximate 95% confidence intervals for each parameter, as shown in Table S3, we form the profile likelihood from steps two and four. Next, we start at either end of the simple parmater bounds until we determine the first grid point where the normalised profile likelihood, $L_p\left(\psi; y^0\right) = \exp\left(l_p\left(\psi; y^0\right) - \max_\theta\left[l\left(\theta; y^0\right)\right]\right)$, is greater than profile likelihood threshold value of 0.15 [11]. These two grid points are a first approximation of the lower and upper 95% confidence interval boundaries. Finally, to obtain a more accurate estimate of the approximate 95% confidence interval boundaries, we consider each of these two grid points in turn as the `FirstGuess` for the MATLAB function *fsolve* [12], and use linear interpolation, with the MATLAB function *interp1* [13].

Key software, including precise details of the implementation of this section, is freely available on a GitHub repository.

#### B.2 Parameter bounds

To interpret $s$ we consider the evolution of the tumour spheroid in phase (i). Equation (S.13) can be written in terms of volume $V(t)$, recalling that the volume of a sphere is $4\pi R_{\mathrm{o}}^3(t)/3$, as

$$V(t) = V(0)\exp(st), \tag{S.30}$$

where $V(0) = 4\pi R_{\mathrm{o}}^3(0)/3$. Then by letting $T$ define the time when $V(T) = 2V(0)$, we relate $s$ to the doubling time of the cells through

$$s = \frac{1}{T}\log_e(2). \tag{S.31}$$

Then assuming that the doubling time is greater than 12 hours (= 1/2 day) we obtain an upper bound of $s$ as $2\log_e(2)$ [day$^{-1}$] $\approx 1.39$ [day$^{-1}$]. Preliminary exploration cofnirms this estimate is very conservative so we set the upper bound to unity.

#### B.3 Pooled sample variances

To identify parameters we assume distinct pooled sample variances for the outer, necrotic, and inhibited radius measurements, as opposed to a single pooled variance for all measurements. In Figure S3 we plot the pooled sample variances for different experimental designs which justify the use of a variance for each measurement type.

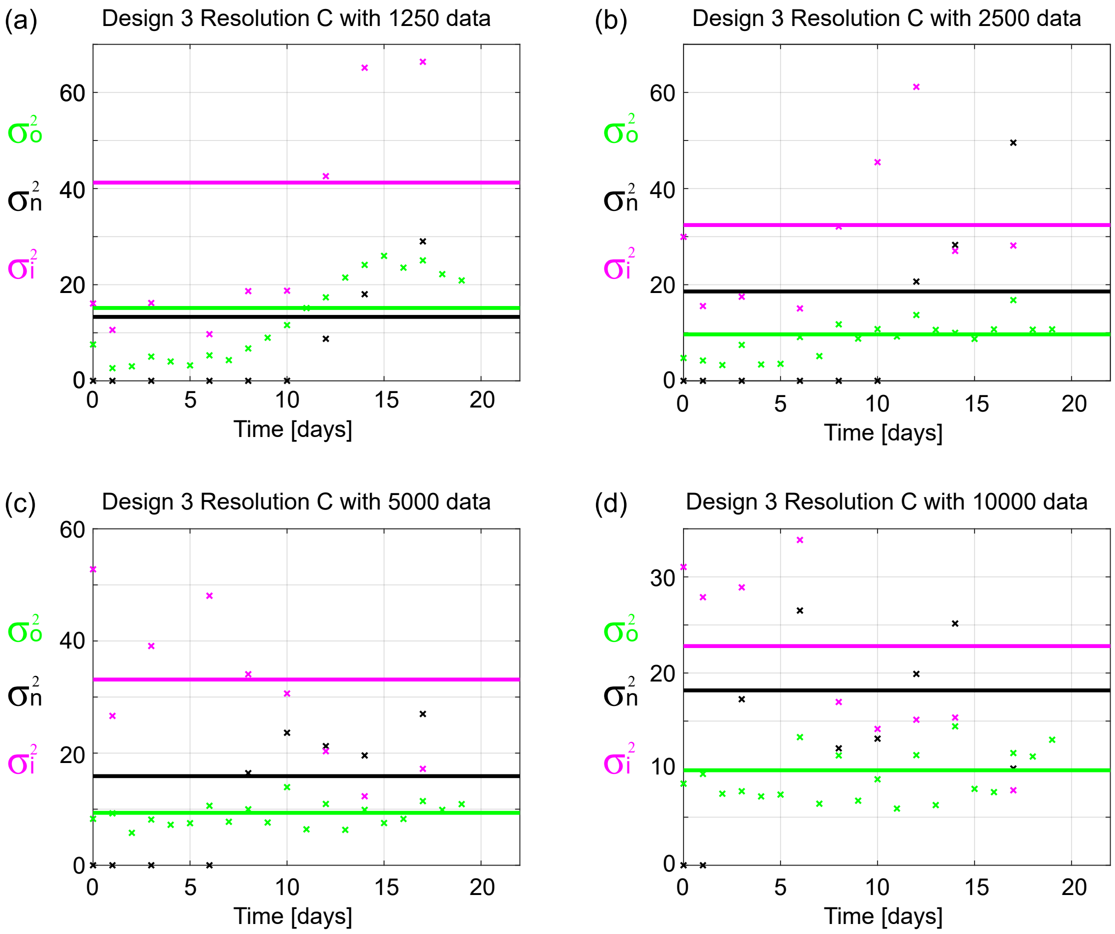


Figure S3: Variances $\sigma_{\mathrm{o}}^2$, $\sigma_{\mathrm{n}}^2$, and $\sigma_{\mathrm{i}}^2$, for outer, necrotic and inhibited radii, respectively. Results shown for WM793b spheroids, using Design 3 and Temporal Resolution C, formed with (a) 1250, (b) 2500, (c) 5000, (d) 10000 cells per spheroid, respectively.

### C Experimental data

#### C.1 Outer radius experimental measurements and images

The IncuCyte S3 live cell imaging system is a useful tool that we use to obtain many outer radius measurements. The other outer radius measurements are obtained from confocal microscopy.

We start with 24 spheroids in the IncuCyte S3 live cell imaging system for each cell line and initial spheroid size and image every 6 hours for the duration of the experiment. However, some measurements could not be obtained primarily due to blurring of the automated imaging, spheroids not forming properly, or spheroids losing their structural integrity at very late time. In Table S1 we show the total number of measurements obtained at 24 hour intervals starting from Day 0 which corresponds to the time that we determined as when spheroid formation ends and growth begins (Supplementary Material C.1.1). In Figures S4-S6 we present representative experimental images obtained from IncuCyte S3 live cell imaging system for different days and WM793b, WM983b, and WM164 cell lines, respectively.

| | WM793 | | | | WM983b | | | WM164 | | | |
|---|---|---|---|---|---|---|---|---|---|---|---|
| Day | 1250 | 2500 | 5000 | 10000 | 2500 | 5000 | 10000 | 1250 | 2500 | 5000 | 10000 |
| 0 | 20 | 24 | 24 | 23 | 24 | 22 | 23 | 23 | 22 | 23 | 21 |
| 1 | 20 | 23 | 24 | 23 | 24 | 22 | 23 | 24 | 22 | 24 | 21 |
| 2 | 20 | 23 | 24 | 23 | 24 | 22 | 23 | 18 | 22 | 24 | 20 |
| 3 | 21 | 23 | 24 | 23 | 24 | 22 | 23 | 19 | 23 | 24 | 20 |
| 4 | 21 | 24 | 24 | 23 | 24 | 22 | 23 | 18 | 23 | 24 | 20 |
| 5 | 21 | 23 | 24 | 23 | 24 | 22 | 22 | 19 | 20 | 19 | - |
| 6 | 21 | 24 | 24 | 23 | 24 | 22 | 21 | 19 | 19 | 19 | - |
| 7 | 21 | 24 | 24 | 23 | 23 | 22 | 20 | - | - | - | - |
| 8 | 21 | 24 | 24 | 23 | 23 | 22 | 20 | - | - | - | - |
| 9 | 21 | 24 | 24 | 23 | 23 | 22 | 20 | - | - | - | - |
| 10 | 21 | 24 | 24 | 23 | 23 | 22 | 20 | - | - | - | - |
| 11 | 21 | 24 | 24 | 23 | 23 | 22 | 20 | - | - | - | - |
| 12 | 21 | 24 | 24 | 23 | 22 | 22 | 20 | - | - | - | - |
| 13 | 21 | 24 | 24 | 23 | 22 | 22 | 20 | - | - | - | - |
| 14 | 21 | 24 | 24 | 23 | - | - | - | - | - | - | - |
| 15 | 20 | 24 | 22 | 23 | 22 | 22 | 20 | - | - | - | - |
| 16 | 18 | 24 | 22 | 23 | 22 | 22 | 20 | - | - | - | - |
| 17 | 18 | 24 | 22 | 23 | 22 | 22 | 20 | - | - | - | - |
| 18 | 19 | 24 | 22 | 23 | 22 | 22 | 20 | - | - | - | - |
| 19 | 19 | 24 | 22 | 23 | 22 | 22 | 20 | - | - | - | - |
| 20 | 13 | 24 | 13 | 11 | - | - | - | - | - | - | - |

Table S1: Number of outer radius measurements obtained from the IncuCyte S3 live cell imaging system for the cell lines WM793b, WM983b, and WM164. Day 0 corresponds to the time that we determined as when spheroid formation ends and growth begins, see supplementary material C.1.1.

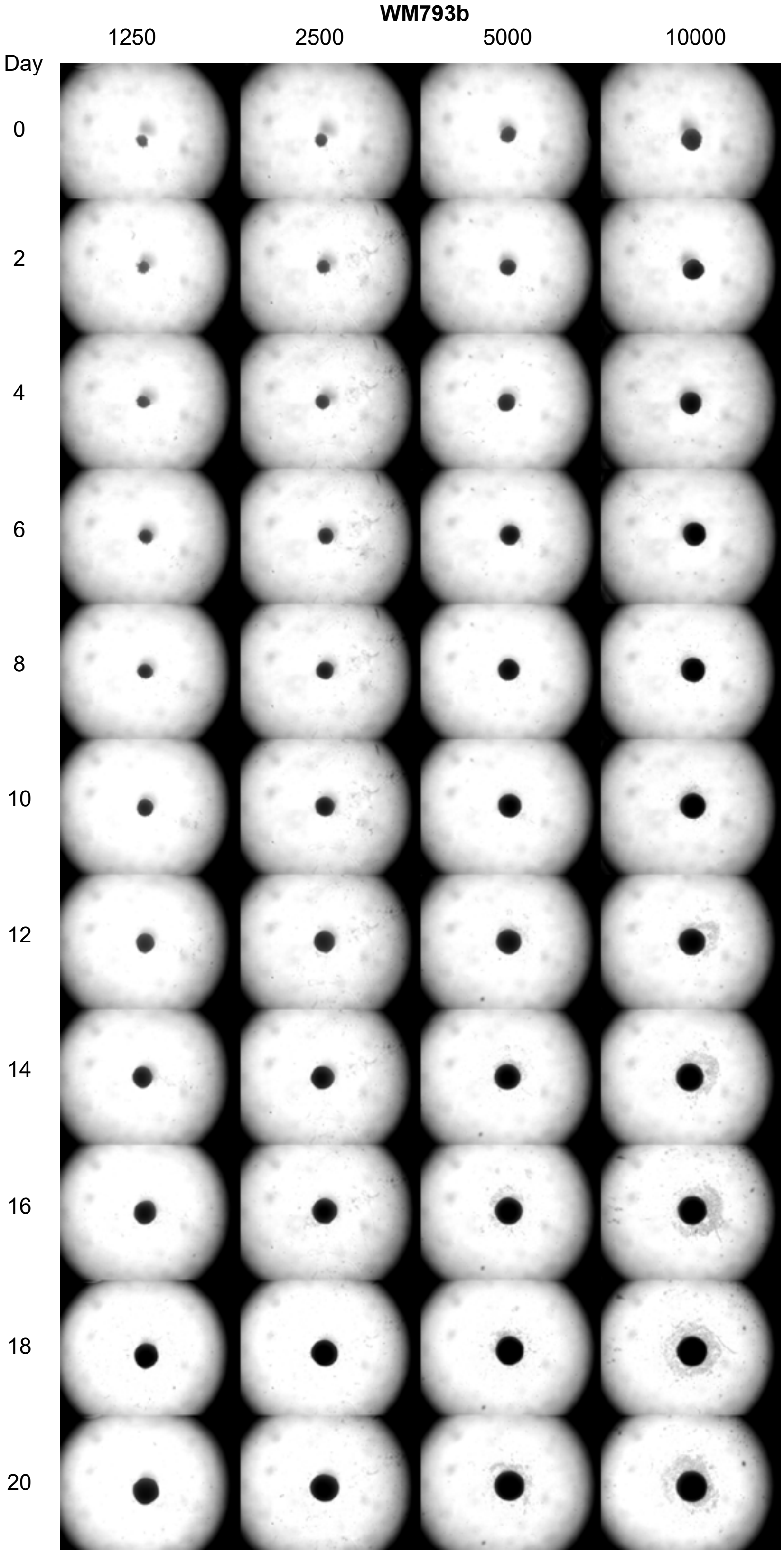


Figure S4: Snapshots of WM793b tumour spheroids from IncuCyte S3 live cell imaging system at 0, 2, 4, 6, 8, 10, 12, 14, 16, 18, and 20 days after formation for tumour spheroids formed with 1250, 2500, 5000, and 10000 cells per spheroid. Each image shows a 4.34 × 3.25 mm field of view.

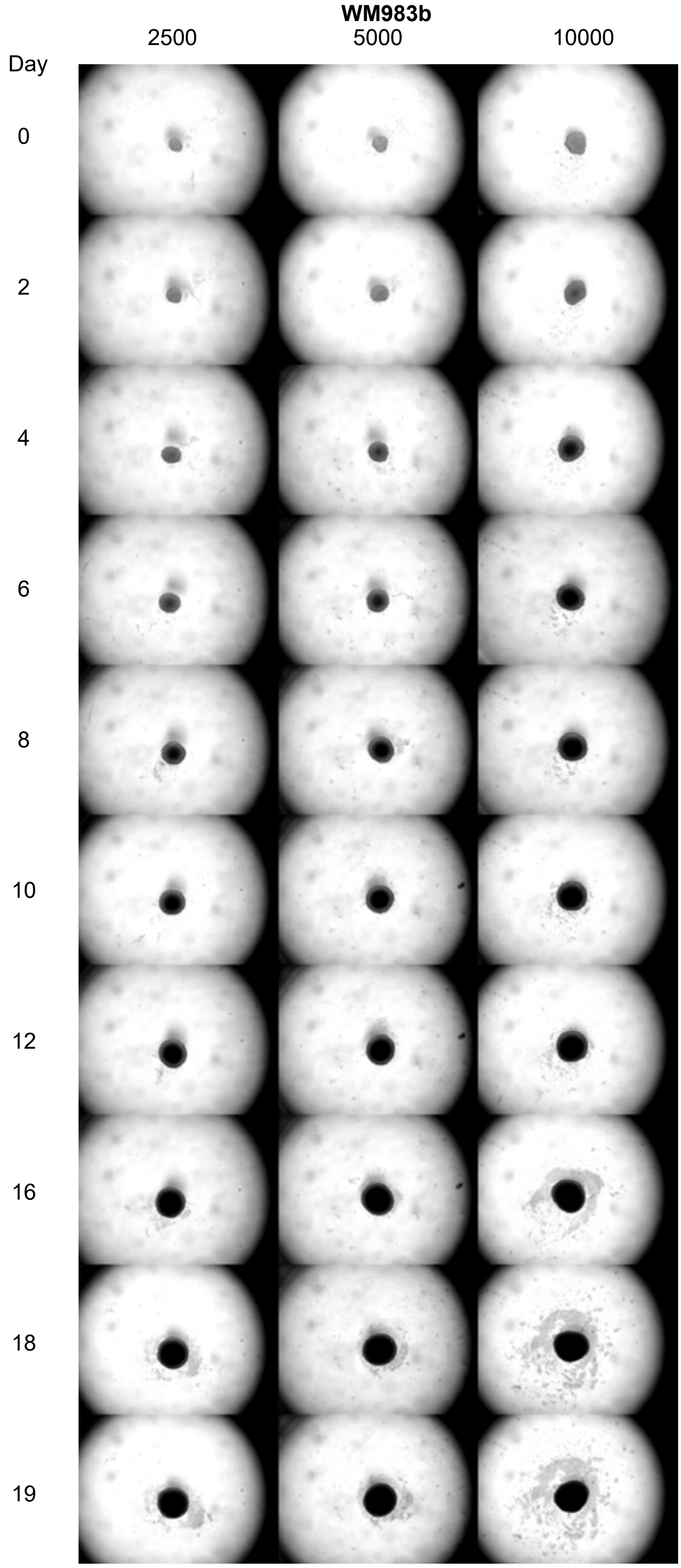


Figure S5: Snapshots of WM983b tumour spheroids from IncuCyte S3 live cell imaging system at 0, 2, 4, 6, 8, 10, 12, 16, 18, and 19 days after formation for tumour spheroids formed with 2500, 5000, and 10000 cells per spheroid. Each image shows a 4.34 × 3.25 mm field of view.

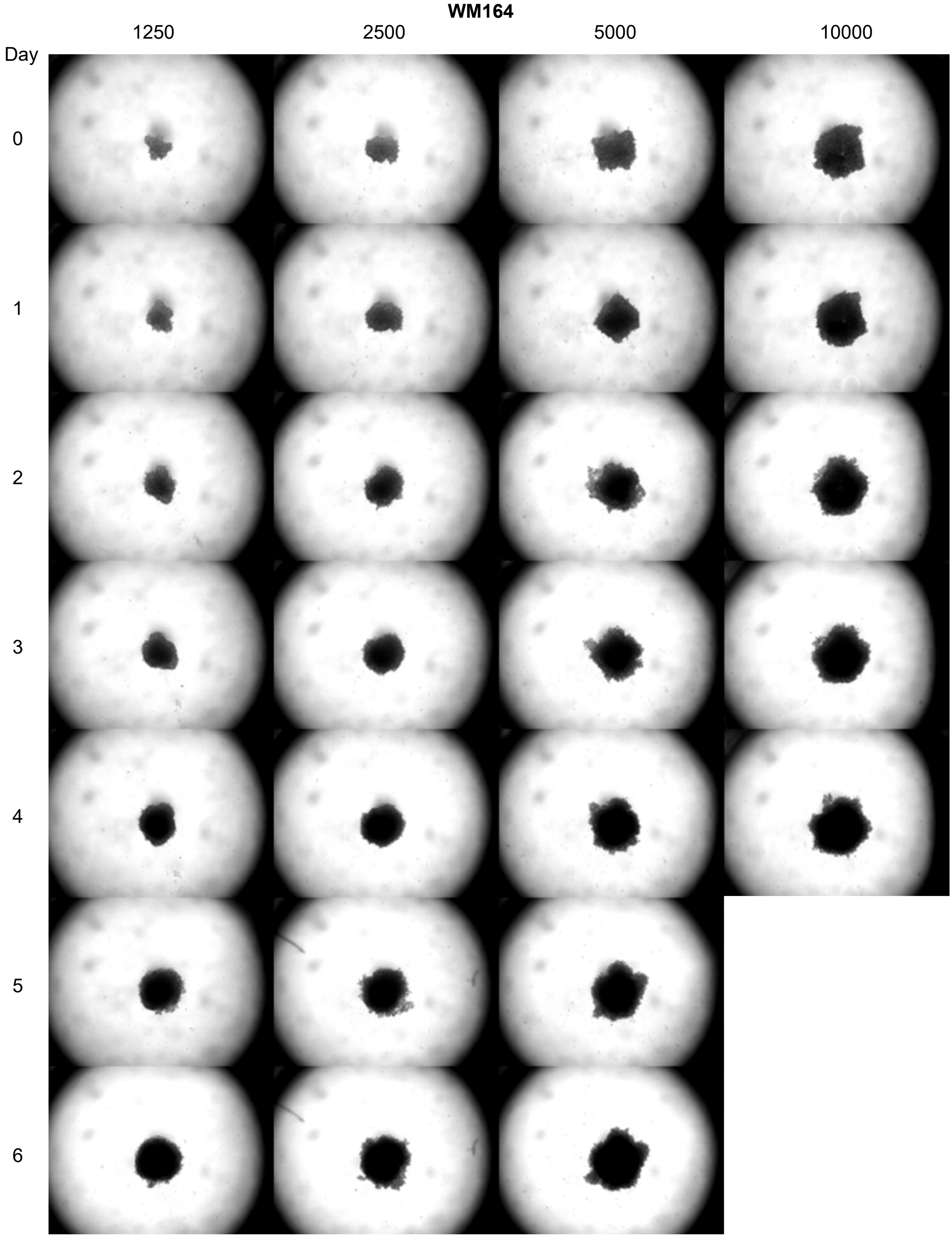


Figure S6: Snapshots of WM164 tumour spheroids from IncuCyte S3 live cell imaging system at 0, 1, 2, 3, 4, 5, and 6 days after formation for tumour spheroids formed with 1250, 2500, 5000, and 10000 cells per spheroid. Each image shows a 4.34 × 3.25 mm field of view.

##### C.1.1 Spheroid formation duration

In Figure S7 we show time snapshots of forming tumour spheroids obtained in the IncuCyte S3 live cell imaging system. These snapshots, alongside monitoring the evolution of the outer radius obtained from image processing, validate the assumption that the tumour spheroids have formed 4 days after seeding for WM793b. This method was also used to determine the duration of spheroid formation for the WM983b (Figure S8) and WM164 (Figure S9) cell lines.

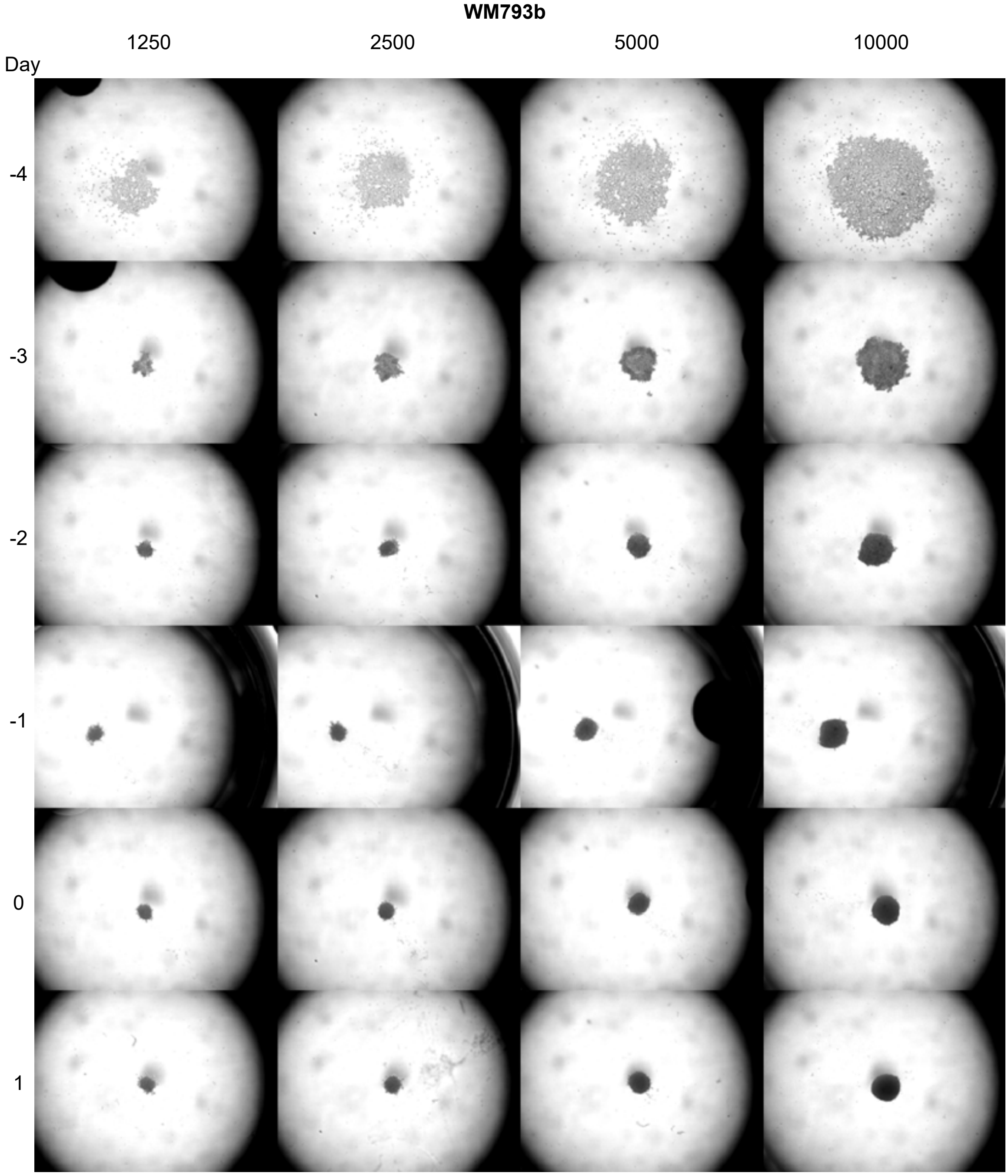


Figure S7: Spheroids are formed 4 days after seeding for WM793b. Snapshots from IncuCyte S3 live cell imaging system at -4, -3, -2, -1, 0, and 1 days after formation for tumour spheroids formed 1250, 2500, 5000, and 10000 cells per spheroid. Each image shows a 4.34 × 3.25 mm field of view.

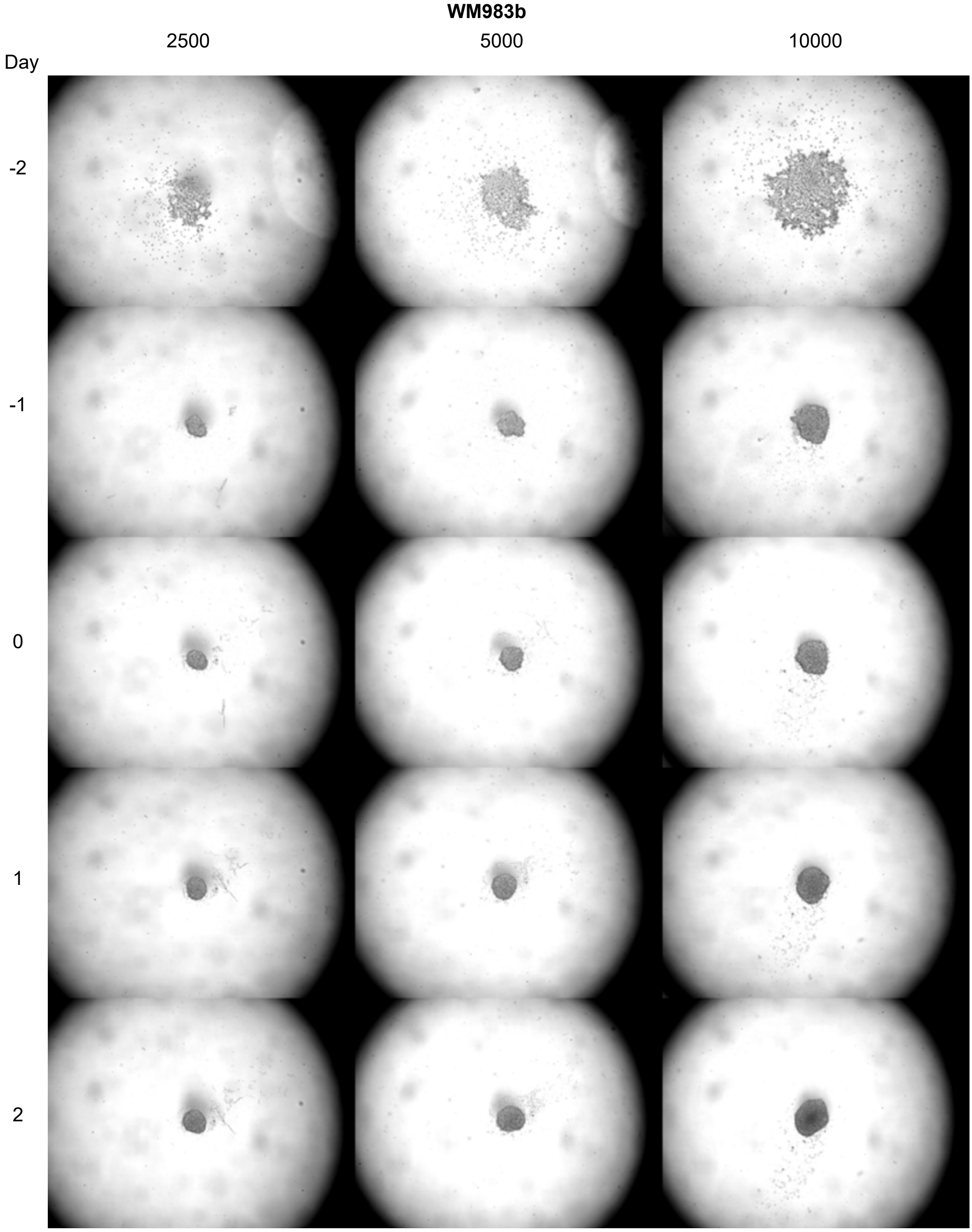


Figure S8: Spheroids are formed at 2 days after seeding for WM983b. Snapshots from IncuCyte S3 live cell imaging system at -2, -1, 0, 1, and 2 days after formation for tumour spheroids formed with 2500, 5000, and 10000 cells per spheroid. Each image shows a 4.34 × 3.25 mm field of view.

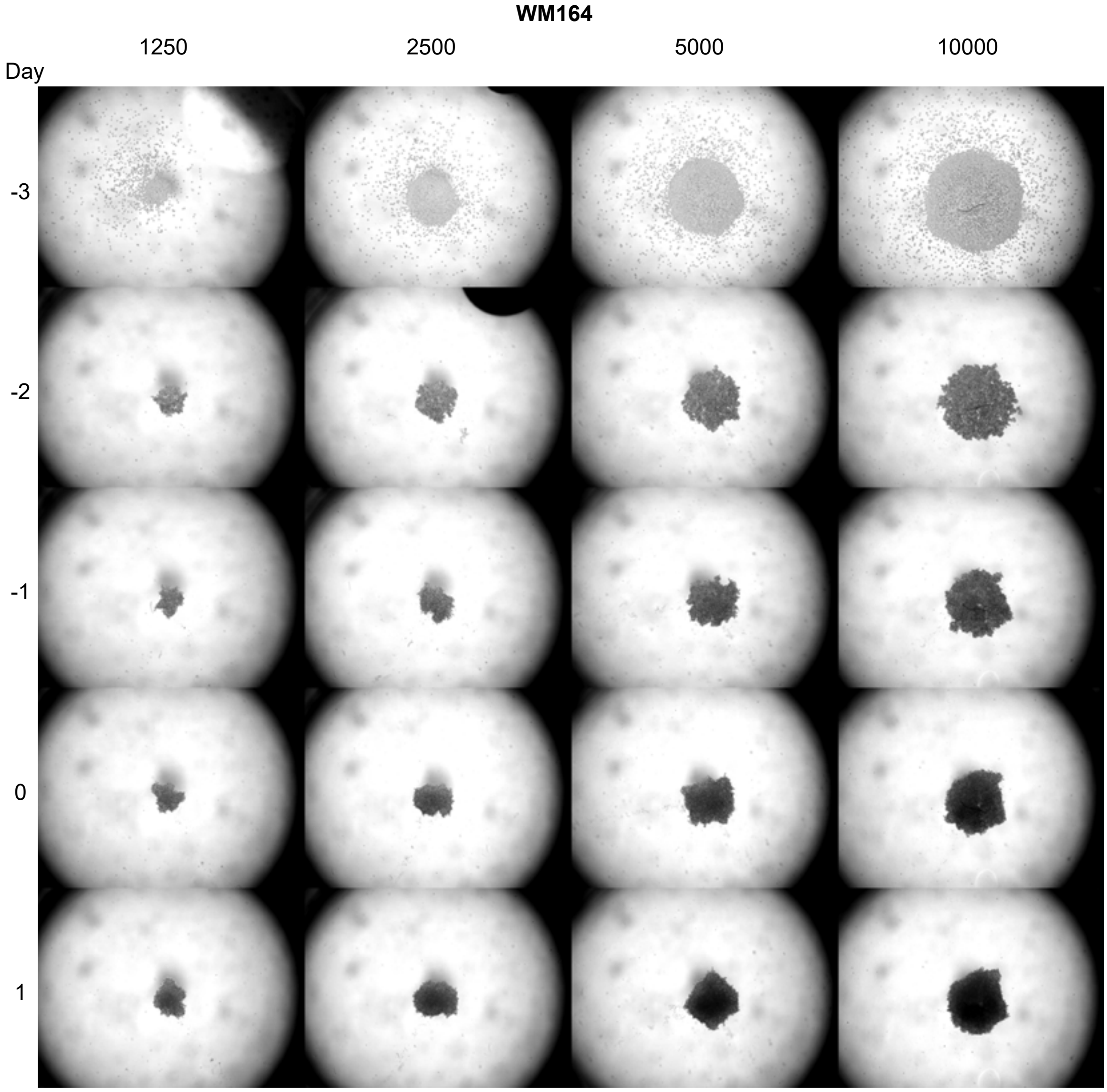


Figure S9: Spheroids are formed at 3 days after seeding for WM164. Snapshots from IncuCyte S3 live cell imaging system at -3, -2, -1, 0 and 1 days after formation for tumour spheroids formed with 1250, 2500, 5000, and 10000 cells per spheroid. Each image shows a 4.34 × 3.25 mm field of view.

#### C.2 Confocal microscopy

##### C.2.1 Measurements

In Table S2 we show the number of confocal measurements obtained. Spheroids damaged during harvesting and fixing procedures are not included.

| | WM793 | | | | WM983b | | | WM164 | | | |
|---|---|---|---|---|---|---|---|---|---|---|---|
| Day | 1250 | 2500 | 5000 | 10000 | 2500 | 5000 | 10000 | 1250 | 2500 | 5000 | 10000 |
| 0 | 5 | 5 | 12 | 7 | - | - | - | - | - | - | - |
| 1 | 4 | 10 | 11 | 12 | 6 | 9 | 6 | 6 | 4 | 14 | 6 |
| 2 | - | - | - | - | 12 | 9 | 10 | 13 | 10 | 10 | 9 |
| 3 | 5 | 22 | 23 | 18 | 12 | 10 | 9 | - | - | - | - |
| 4 | - | - | - | - | - | - | - | 13 | 8 | - | - |
| 5 | - | - | - | - | 20 | 15 | 18 | - | - | - | - |
| 6 | 7 | 28 | 25 | 25 | - | - | - | - | - | - | - |
| 7 | - | - | - | - | - | - | - | - | - | - | - |
| 8 | 12 | 27 | 20 | 23 | 16 | 13 | 15 | - | - | - | - |
| 9 | - | - | - | - | - | - | - | - | - | - | - |
| 10 | 8 | 19 | 21 | 15 | 16 | 17 | 21 | - | - | - | - |
| 11 | - | - | - | - | - | - | - | - | - | - | - |
| 12 | 12 | 18 | 19 | 17 | 13 | 14 | 13 | - | - | - | - |
| 13 | - | - | - | - | - | - | - | - | - | - | - |
| 14 | 15 | 19 | 22 | 21 | 17 | 21 | 19 | - | - | - | - |
| 15 | - | - | - | - | - | - | - | - | - | - | - |
| 16 | - | - | - | - | 11 | 20 | 19 | - | - | - | - |
| 17 | 11 | 15 | 14 | 5 | - | - | - | - | - | - | - |
| 18 | - | - | - | - | - | - | - | - | - | - | - |
| 19 | - | - | - | - | 25 | 31 | 16 | - | - | - | - |
| 20 | 22 | 23 | 21 | 20 | - | - | - | - | - | - | - |

Table S2: Number of spheroids imaged using confocal microscopy for the cell lines WM793b, WM983b, and WM164. For each imaged spheroid we obtain a measurement of the outer radius, inhibited radius, and necrotic radius. Day 0 corresponds to the time for the each cell line that we determined as when spheroid formation ends and growth begins. Measurements were taken on days days 3, 4, 5, 7, 10, 12, 14, 16, 18, 21 and 24 after seeding, and appear on different days in the table due to the different formation times.

#### C.3 Confocal microscopy supplementary experimental images

Here we present confocal microscopy images of spheroids formed with the WM793b, WM983b, and WM164 cell lines. In the images we outline each spheroids outer boundary, inhibited region, and necrotic region.

### C.3.1 WM793b

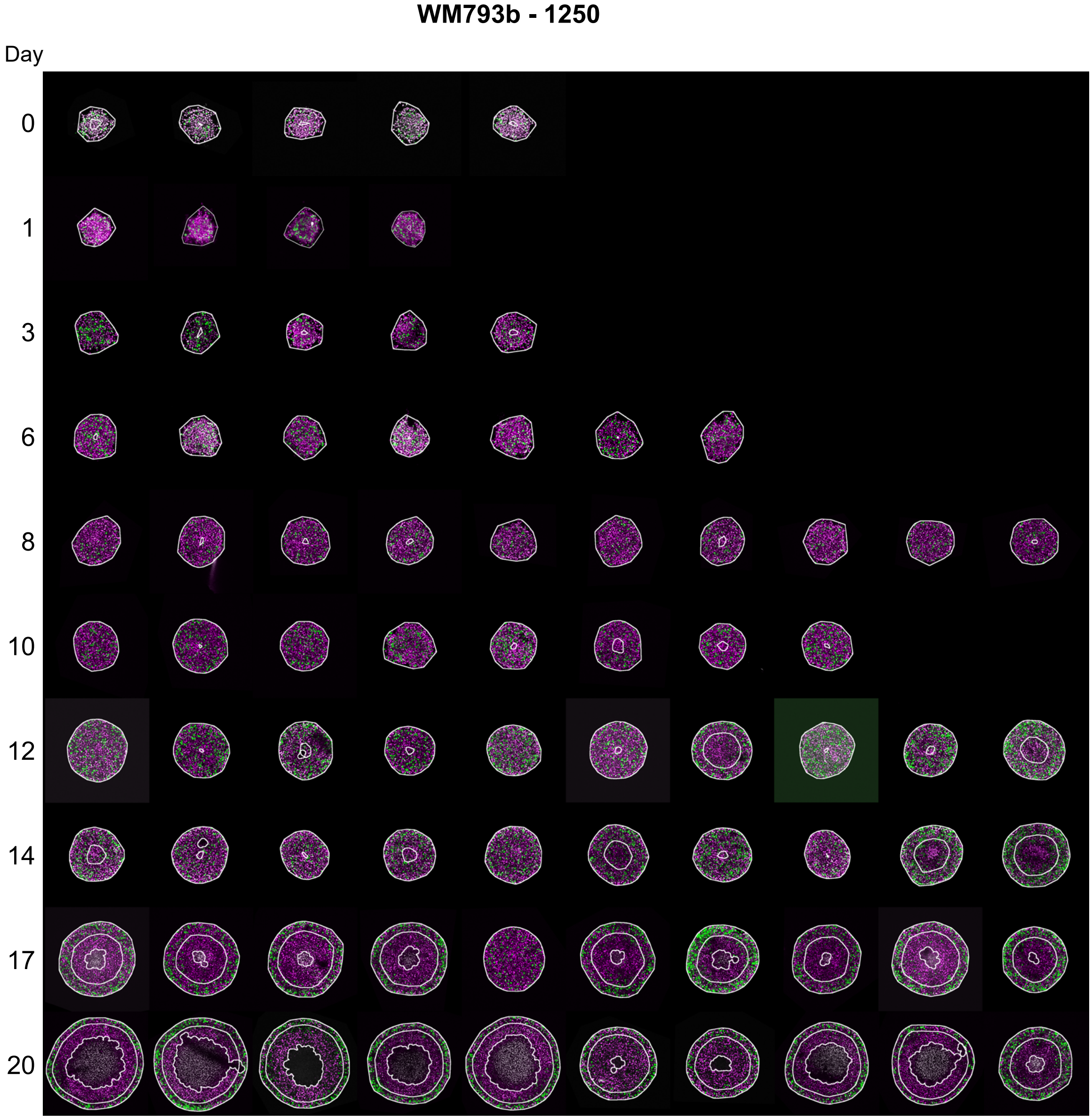


Figure S10: Experimental images of WM793b tumour spheroids formed with 1250 cells per spheroid. Each image shows a 800×800 µm field of view.

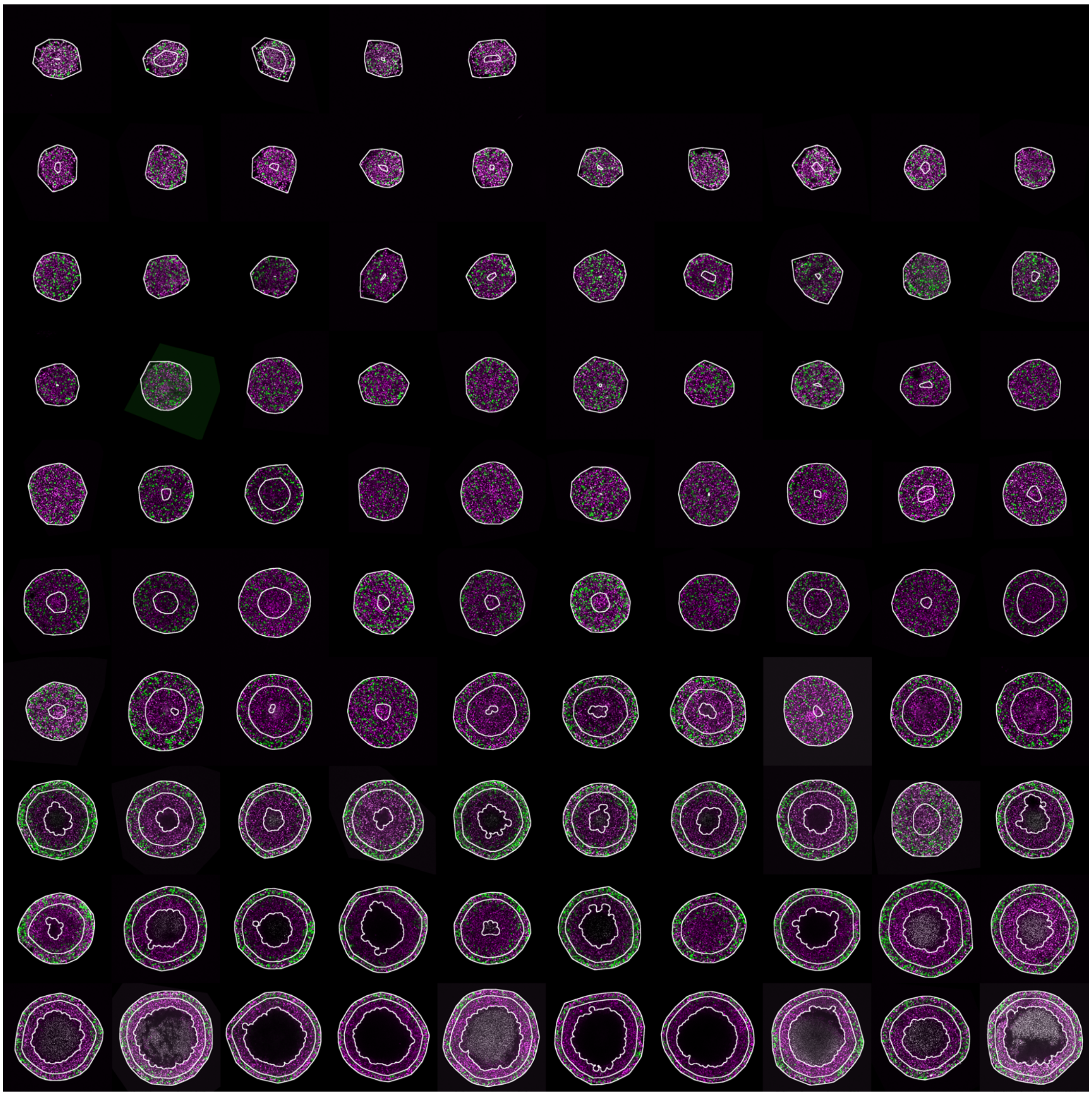


Figure S11: Experimental images of WM793b tumour spheroids formed with 2500 cells per spheroid. Each image shows a 800×800 µm field of view.

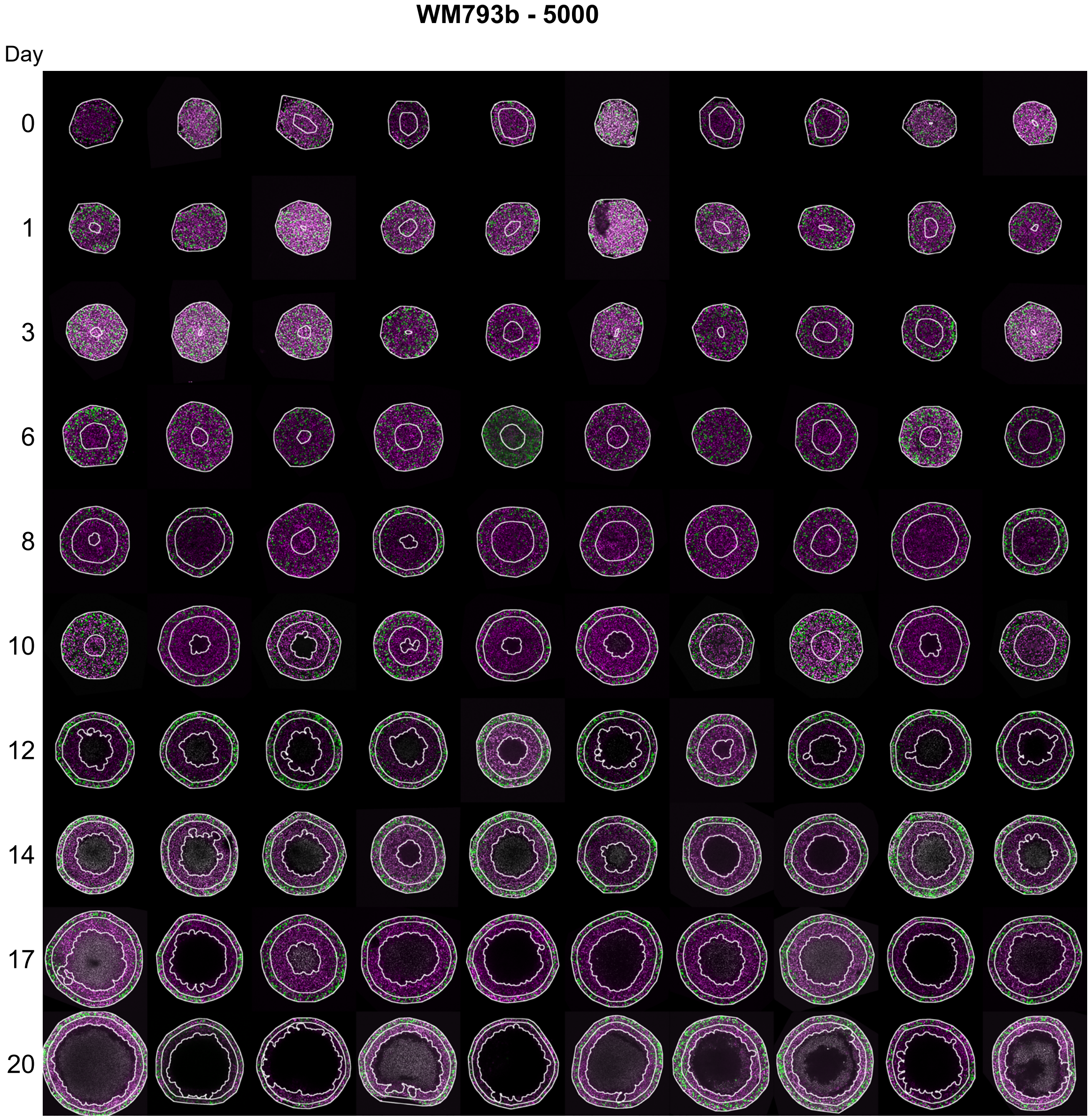


Figure S12: Experimental images of WM793b tumour spheroids formed with 5000 cells per spheroid. Each image shows a 800×800 μm field of view.

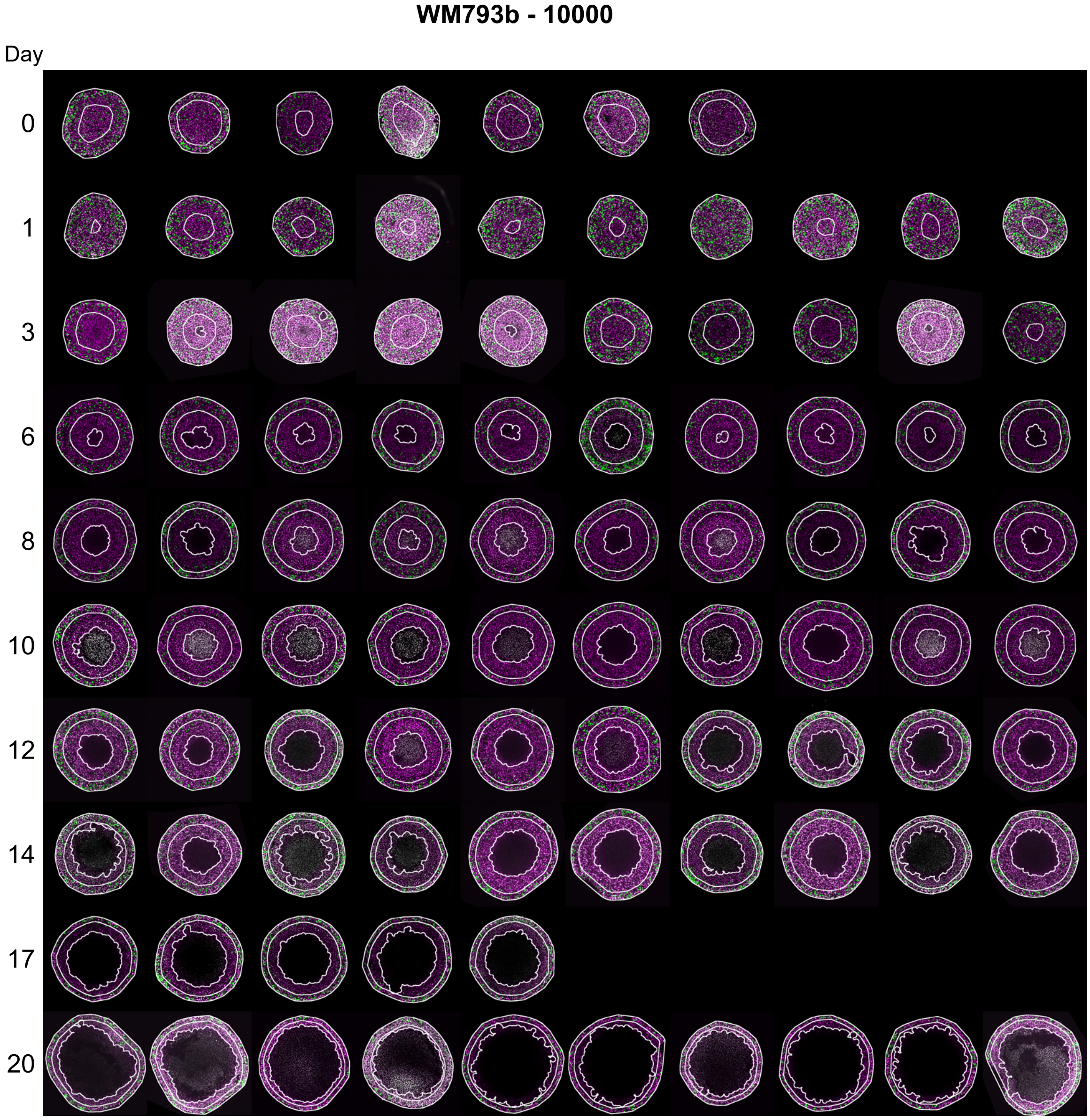


Figure S13: Experimental images of WM793b tumour spheroids formed with 10000 cells per spheroid. Each image shows a 800×800 µm field of view.

### C.3.2 WM983b

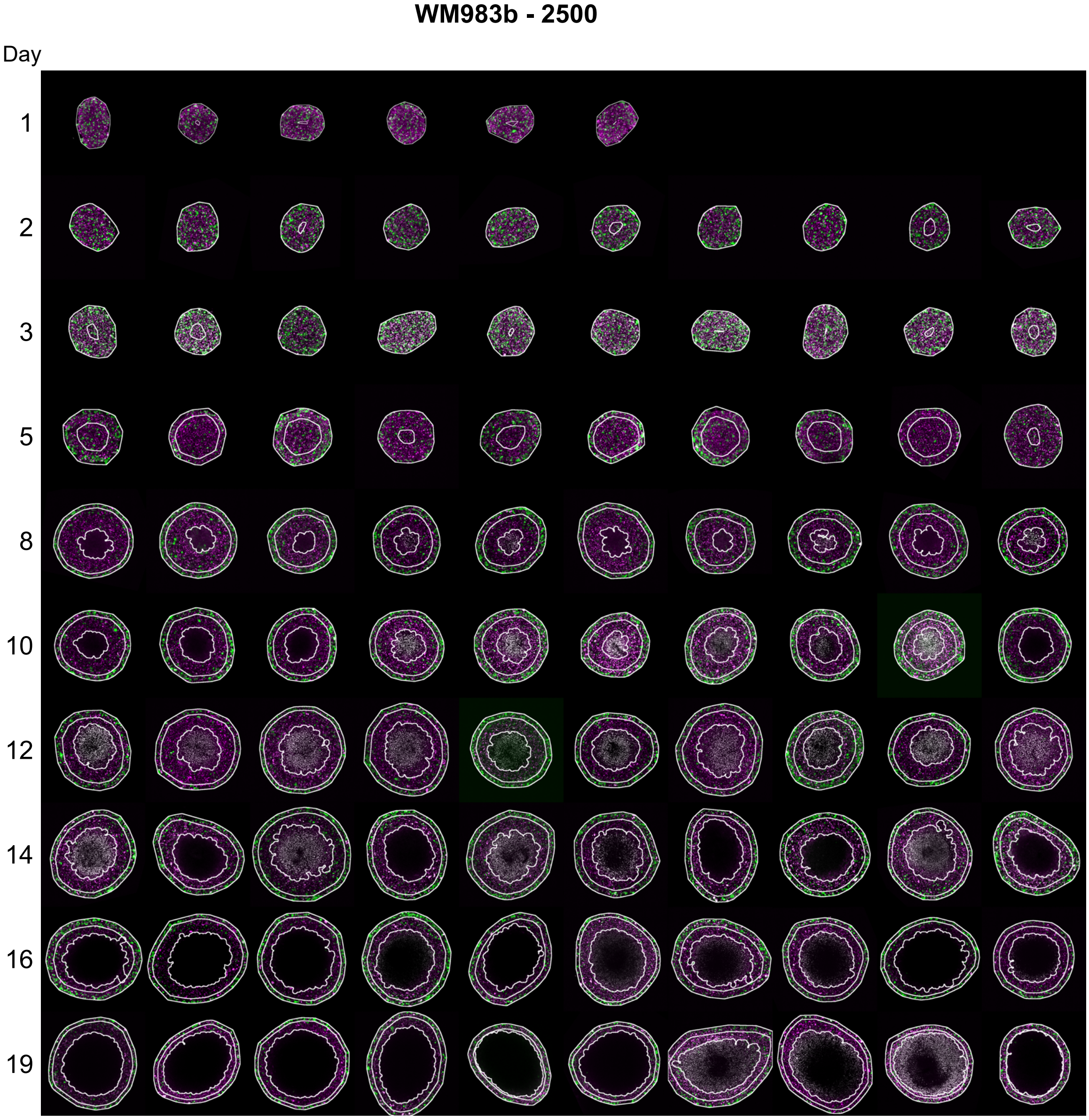


Figure S14: Experimental images of WM983b tumour spheroids formed with 2500 cells per spheroid. Each image shows a 800×800 µm field of view.

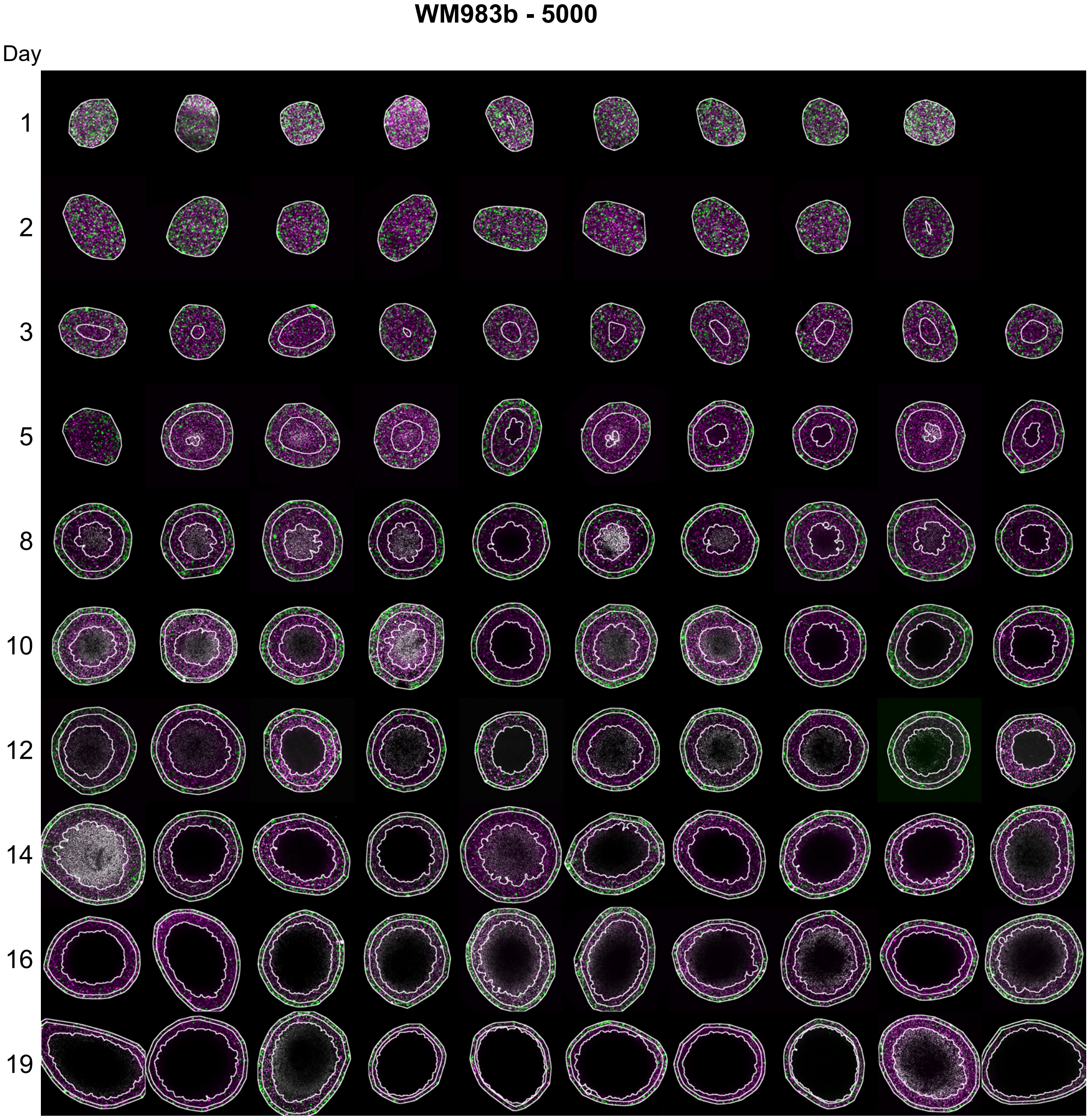


Figure S15: Experimental images of WM983b tumour spheroids formed with 5000 cells per spheroid. Each image shows a 800×800 µm field of view.

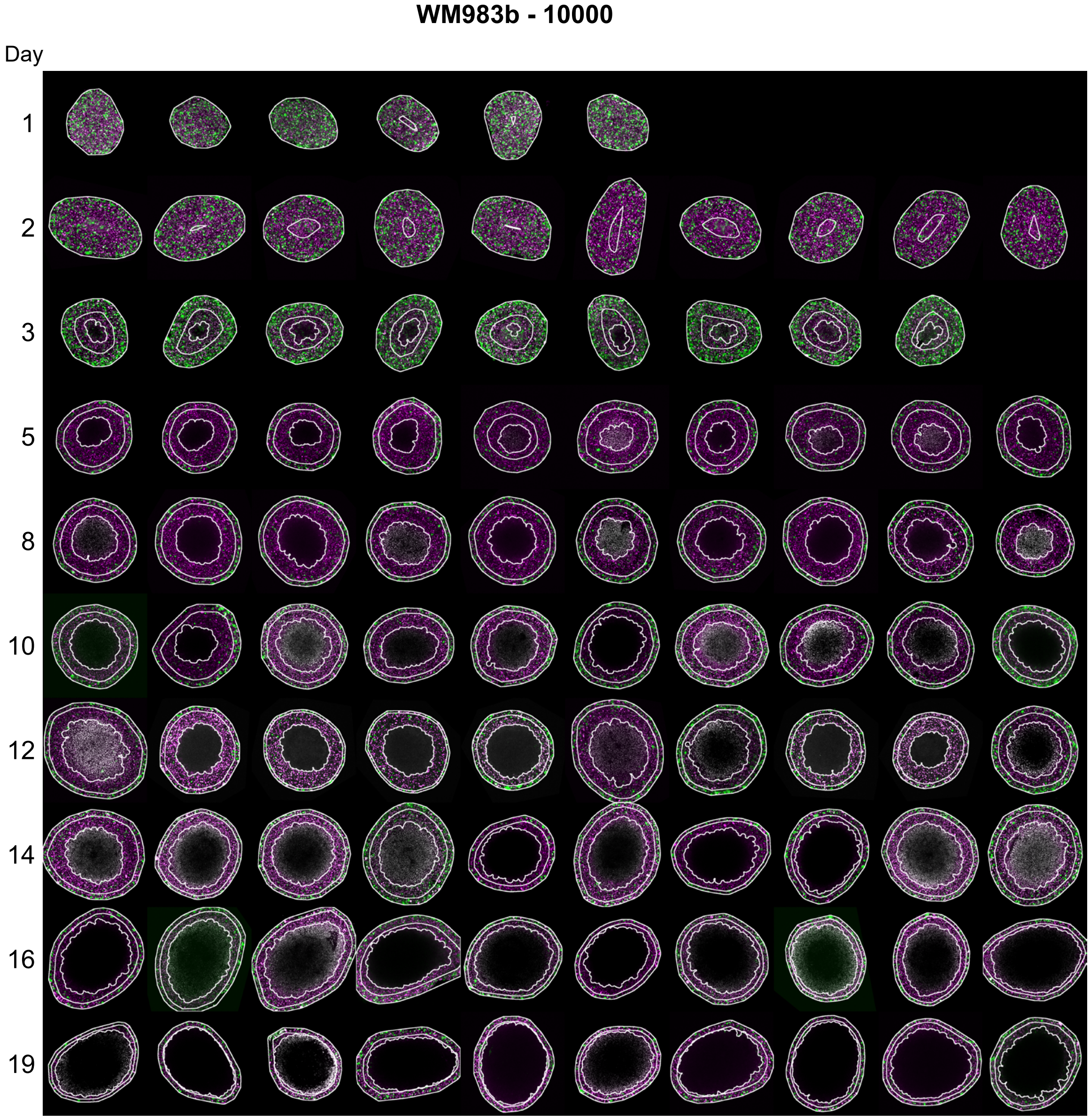


Figure S16: Experimental images of WM983b tumour spheroids formed with 10000 cells per spheroid. Each image shows a 800×800 μm field of view.

### C.3.3 WM164

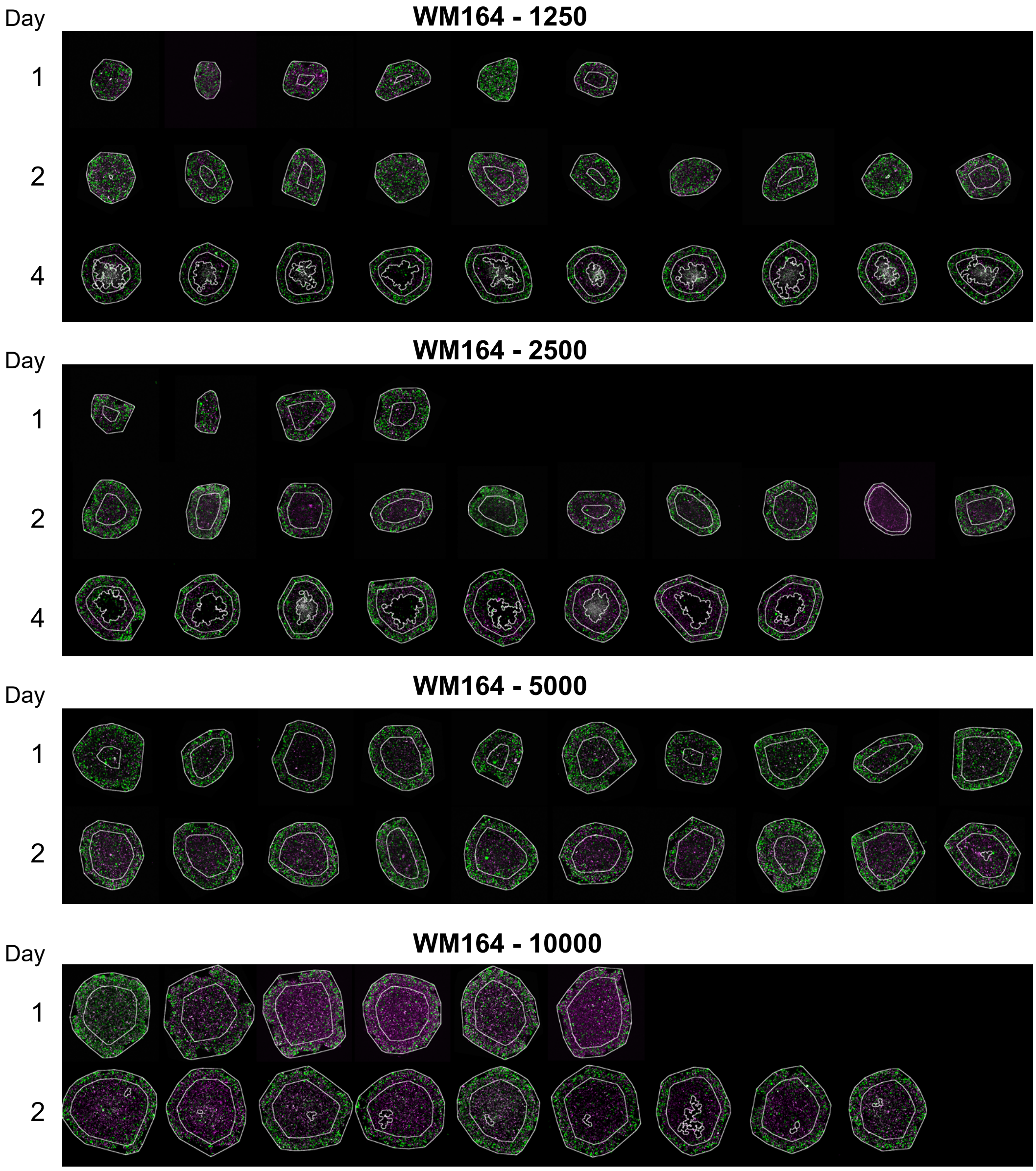


Figure S17: Experimental images of WM164 tumour spheroids formed with 1250, 2500, 5000, and 10000 cells per spheroid. Each image shows a 800×800 µm field of view.

##### C.3.4 3D rendering

Here we present a 3D rendering of a confocal microscopy image z-stack of half of a FUCCI-melanoma WM793b spheroid 17 days after formation with 5000 cells.

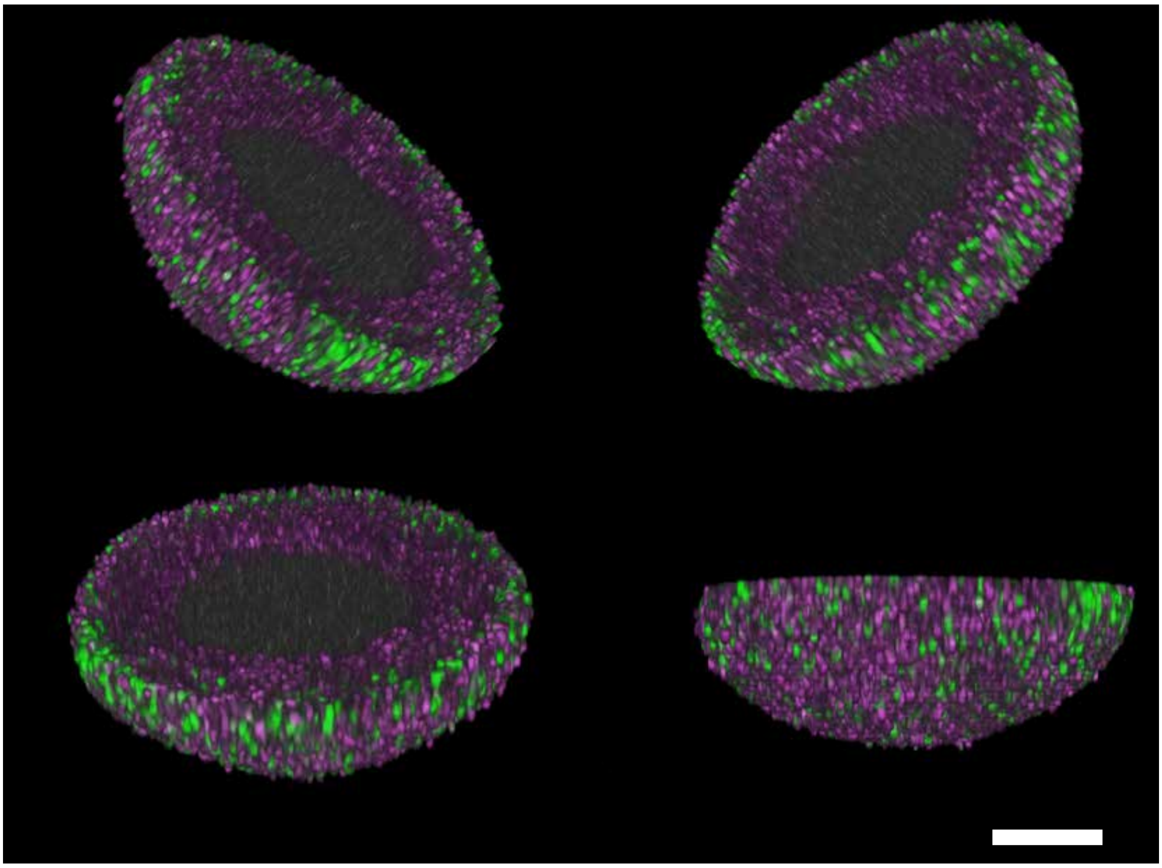

Figure S18: 3D rendering of half of a FUCCI-melanoma WM793b spheroid 17 days after formation with 5000 cells. Scale bar 200 µm.

### D WM793b additional results

#### D.1 Results in tables

In all figures with profile likelihoods we include a red-dashed horizontal line at 0.15 indicating the 95% confidence interval threshold value [11]. Here, in Table S3 we present the corresponding MLE's and approximate 95% confidence intervals for a range of experimental designs.

| Cell line | Figure | Experimental Design | $Q$ | $\gamma$ | $s$ [day$^{-1}$] | $R_{\mathrm{c}}$ [µm] | $R_o(0)$ [µm] |
|---|---|---|---|---|---|---|---|
| WM793b | 2 | 1 Res A | 0.711 (0.479, 1.000) | 0.020 (0.010, 6.000) | 0.141 (0.127, 1.000) | 299.980 (25.000, 350.000) | 183.509 (179.118, 186.825) |
| | | 1 Res B | 0.675 (0.550, 1.000) | 0.010 (0.010, 6.000) | 0.143 (0.128, 0.324) | 312.725 (85.574, 350.000) | 182.940 (178.565, 186.470) |
| | | 1 Res C | 0.885 (0.609, 1.000) | 0.484 (0.010, 6.000) | 0.134 (0.128, 0.144) | 258.100 (203.186, 350.000) | 183.415 (181.221, 185.260) |
| WM793b | 3 | 2 Res A | 1.000 (0.940, 1.000) | 0.924 (0.299, 1.262) | 0.126 (0.121, 0.133) | 257.056 (254.484, 259.373) | 184.543 (181.892, 187.082) |
| | | 3 Res A | 0.770 (0.753, 0.784) | 0.010 (0.010, 0.110) | 0.159 (0.153, 0.166) | 260.672 (257.408, 262.899) | 178.733 (176.178, 181.102) |
| WM793b | 4 | 3 Res A 1250 MLE | 0.847 (0.824, 0.869) | 0.010 (0.010, 6.000) | 0.134 (0.126, 0.141) | 250.352 (243.485, 256.425) | 126.100 (122.459, 129.875) |
| | | 3 Res A 2500 MLE | 0.829 (0.817, 0.838) | 0.010 (0.010, 0.641) | 0.150 (0.145, 0.157) | 256.818 (253.607, 259.141) | 143.797 (141.164, 146.444) |
| | | 3 Res A 5000 MLE | 0.770 (0.753, 0.784) | 0.010 (0.010, 0.110) | 0.159 (0.153, 0.166) | 260.672 (257.408, 262.899) | 178.733 (176.178, 181.102) |
| | | 3 Res A 10000 MLE | 0.832 (0.816, 0.846) | 1.130 (0.919, 1.333) | 0.149 (0.139, 0.161) | 253.692 (251.158, 255.826) | 223.118 (219.825, 226.245) |
| WM983b | S39 | 3 Res A 2500 MLE | 0.801 (0.779, 0.822) | 0.395 (0.334, 0.457) | 0.319 (0.304, 0.336) | 209.774 (205.679, 214.047) | 128.274 (125.128, 131.309) |
| | | 3 Res A 5000 MLE | 0.824 (0.815, 0.832) | 0.398 (0.366, 0.445) | 0.316 (0.300, 0.333) | 210.220 (208.168, 212.156) | 155.949 (152.684, 159.075) |
| | | 3 Res A 10000 MLE | 0.841 (0.818, 0.862) | 0.517 (0.475, 0.562) | 0.368 (0.341, 0.398) | 209.470 (205.893, 212.974) | 187.514 (183.013, 191.669) |
| WM164 | S40 | 3 Res A 1250 MLE | 0.891 (0.856, 0.923) | 0.010 (0.010, 0.408) | 0.430 (0.398, 0.465) | 325.842 (315.384, 335.715) | 230.322 (222.762, 238.128) |
| | | 3 Res A 2500 MLE | 0.813 (0.779, 0.842) | 0.010 (0.010, 1.006) | 0.419 (0.383, 0.487) | 357.305 (351.641, 363.356) | 275.862 (263.881, 285.260) |
| | | 3 Res A 5000 MLE | 0.735 (0.530, 0.770) | 1.365 (0.010, 6.000) | 0.380 (0.298, 0.478) | 434.753 (428.901, 600.000) | 355.357 (339.002, 370.630) |
| | | 3 Res A 10000 MLE | 0.701 (0.665, 0.736) | 0.127 (0.010, 6.000) | 0.287 (0.226, 0.394) | 528.532 (519.431, 538.034) | 471.672 (450.601, 488.117) |

Table S3: Most likely estimates and approximate 95% confidence intervals for a range of experimental designs. Results shown to three decimal places.

#### D.2 Measurement times and experimental duration

Figure 1 shows that varying the temporal resolution in Design 1 is not sufficient to predict necrotic and inhibited radii. Here, in Figures S19 and S20, we show that varying the temporal resolution using Designs 2 and 3, respectively, gives consistent results across temporal resolutions A, B, and C.

Next we consider four additional experimental designs that use different temporal measurements

- Temporal Resolution D: the first 4 days (Day 1, 2, 3),
- Temporal Resolution E: the first 10 days (Day 1, 2, 3, 4, 5, 6, 7, 8, 9, 10),
- Temporal Resolution F: the last 10 days (Day 10, 11, 12, 13, 14, 15, 16, 17, 18, 19),
- Temporal Resolution G: the last 4 days (Day 16, 17, 18, 19),

In Figures S21 and S22, we present results for the WM793b cell line for spheroids formed with 1250 and 5000 cells, respectively. These results show that using Temporal Resolution D is not sufficient to predict late time behaviour (Figure S21e, Figure S22e) and Temporal Resolution E can also not be sufficient to predict late time behaviour (Figure S21f). Similarly, using late time experimental measurements, as in Temporal Resolutions F and G, is insufficient to determine tumour spheroid structure at early times (Figure S22g-h).

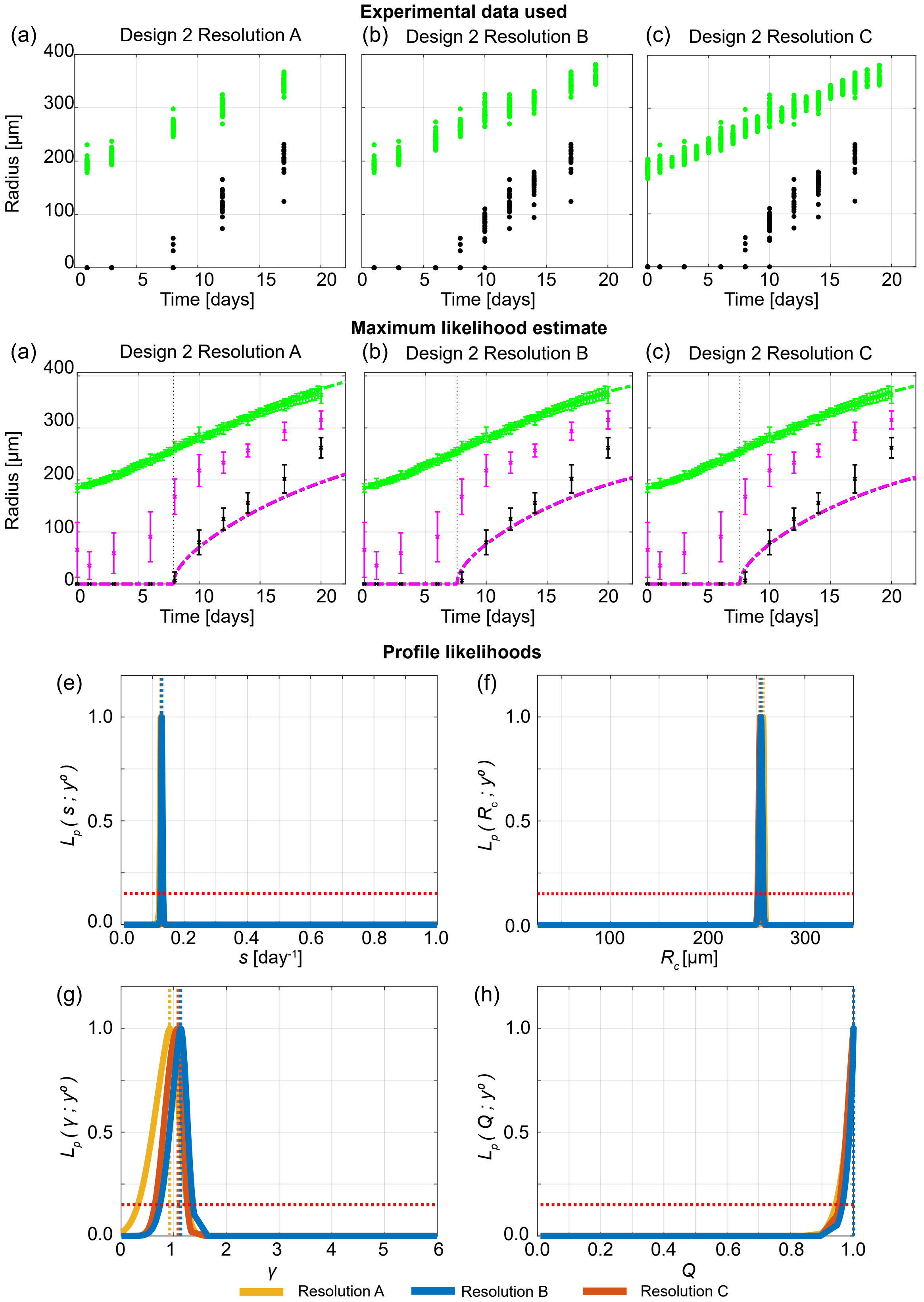


Figure S19: Increasing times when outer and necrotic radius is measured gives consistent information. (a)-(c) Experimental data used in Design 2 with Temporal Resolutions A, B, and C. Profile likelihoods for (e) $s$, (f) $R_c$, (g) $\gamma$, (h) $Q$. Yellow, blue, and orange lines in (e)-(h) represent profile likelihoods from Design 2 with Temporal Resolutions A, B, and C, respectively. (i)-(k) Comparison of Greenspan model simulated with maximum likelihood estimate compared to full experimental data set for Design 2 with Temporal Resolutions A, B, and C, where error bars show standard deviation.

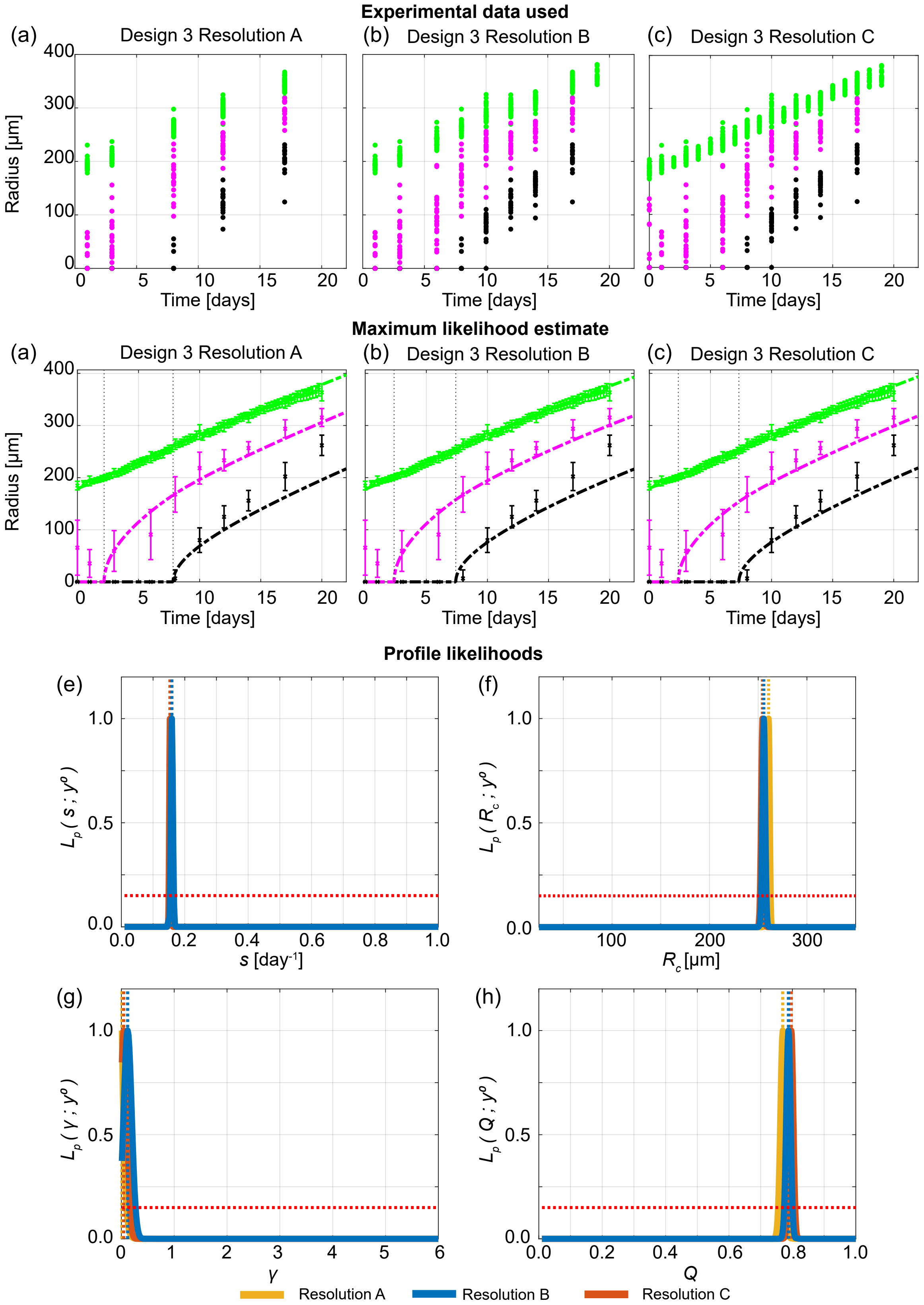


Figure S20: Increasing times when outer, necrotic and inhibited radius and necrotic is measured gives consistent information. (a)-(c) Experimental data used in Design 3 with Temporal Resolutions A, B, and C. Profile likelihoods for (e) $s$, (f) $R_c$, (g) $\gamma$, (h) $Q$. Yellow, blue, and orange lines in (e)-(h) represent profile likelihoods from Design 3 with Temporal Resolutions A, B, and C, respectively. (i)-(k) Comparison of Greenspan model simulated with maximum likelihood estimate compared to full experimental data set for Design 3 with Temporal Resolutions A, B, and C, where error bars show standard deviation.

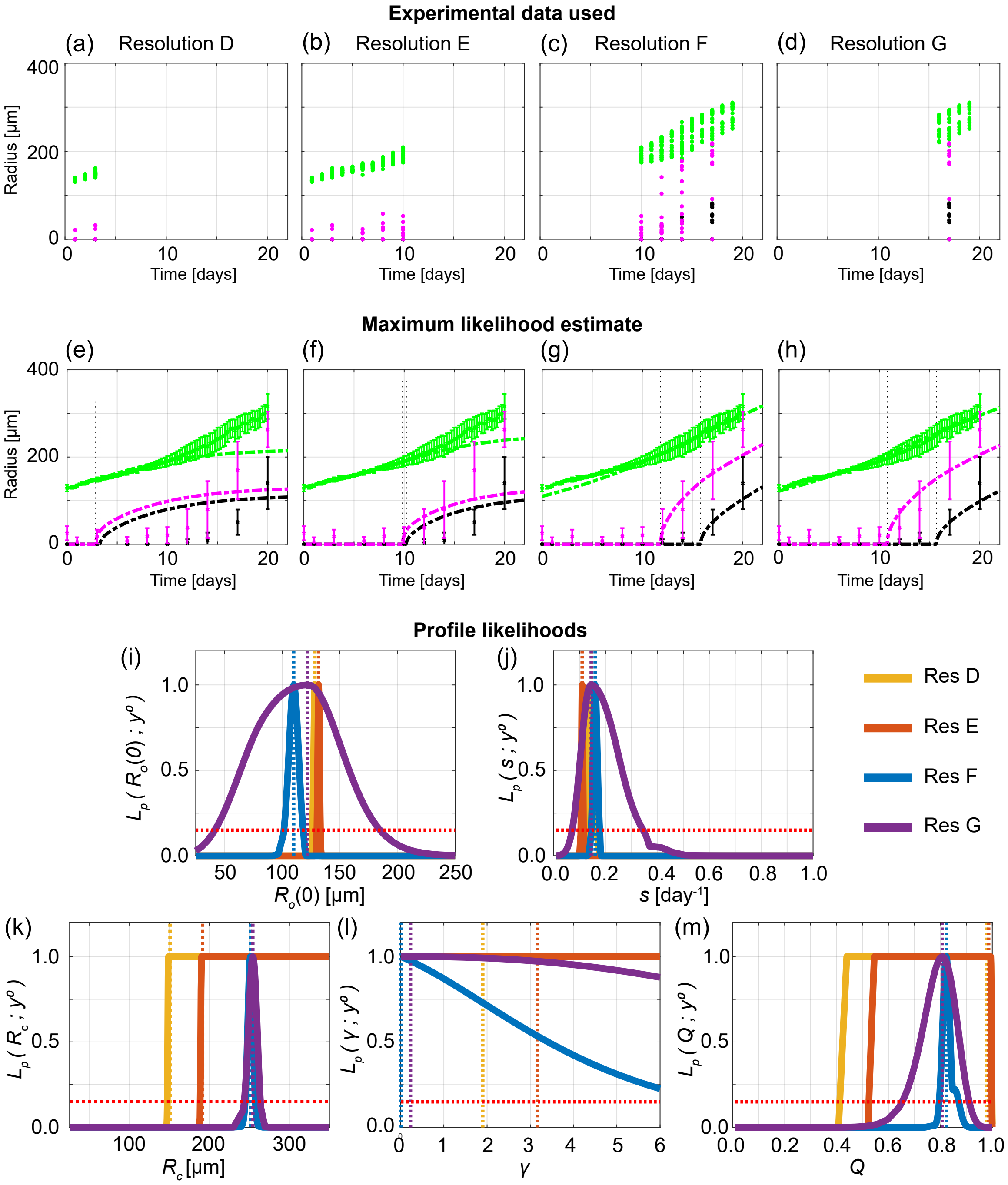


Figure S21: Design 3 with temporal Resolutions D, E, F, and G for WM793b tumour spheroids formed with 1250 cells per spheroid. (a)-(d) Experimental data used in Design 3 Temporal Resolutions D, E, F, and G, respectively. (e)-(h) Comparison of Greenspan model simulated with maximum likelihood estimate compared to full experimental data set for Design 3 Temporal Resolutions D, E, F, and G, respectively, where error bars show standard deviation. Profile likelihoods for (i) $R_\mathrm{o}(0)$, (j) $s$, (k) $R_\mathrm{c}$, (l) $\gamma$, (m) $Q$. Yellow, orange, blue, and purple lines in (e)-(h) represent profile likelihoods from Designs 3 Temporal Resolutions D, E, F, and G, respectively.

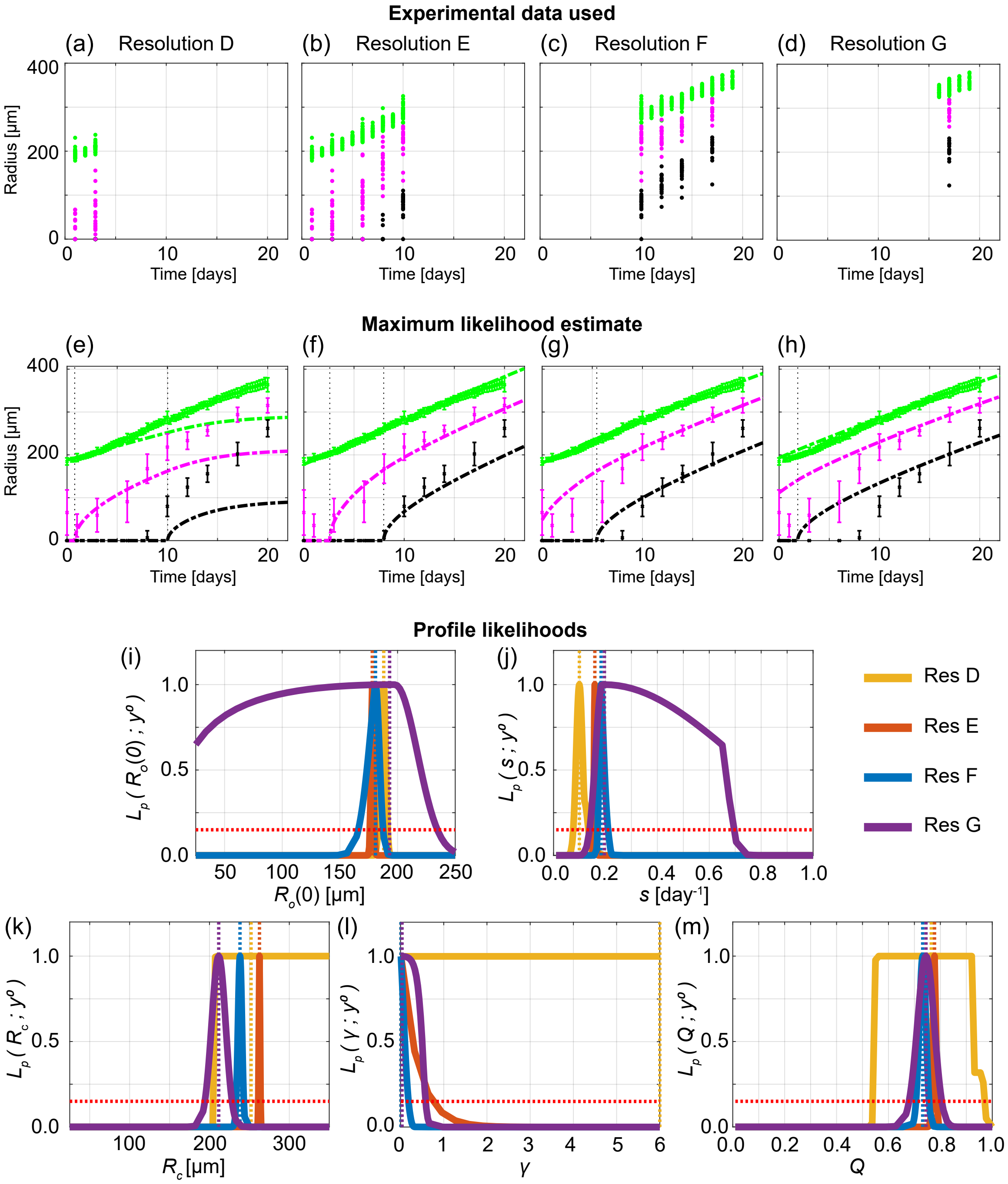


Figure S22: Design 3 with temporal Resolutions D, E, F, and G for WM793b tumour spheroids formed with 5000 cells per spheroid. (a)-(d) Experimental data used in Design 3 Temporal Resolutions D, E, F, and G, respectively. (e)-(h) Comparison of Greenspan model simulated with maximum likelihood estimate compared to full experimental data set for Design 3 Temporal Resolutions D, E, F, and G, respectively, where error bars show standard deviation. Profile likelihoods for (i) $s$, (j) $R_{\mathrm{o}}(0)$, (k) $R_{\mathrm{c}}$, (l) $\gamma$, (m) $Q$. Yellow, orange, blue, and purple lines in (e)-(h) represent profile likelihoods from Designs 3 Temporal Resolutions D, E, F, and G, respectively.

#### D.3 Profile likelihoods for $R_\mathrm{o}(0)$

To perform statistical identifiability analysis we we treat the initial outer radius, $R_\mathrm{o}(0)$, as a parameter. Here, in Figure S23 we show that profile likelihoods for $R_\mathrm{o}(0)$ are consistent across temporal resolutions and experimental designs.

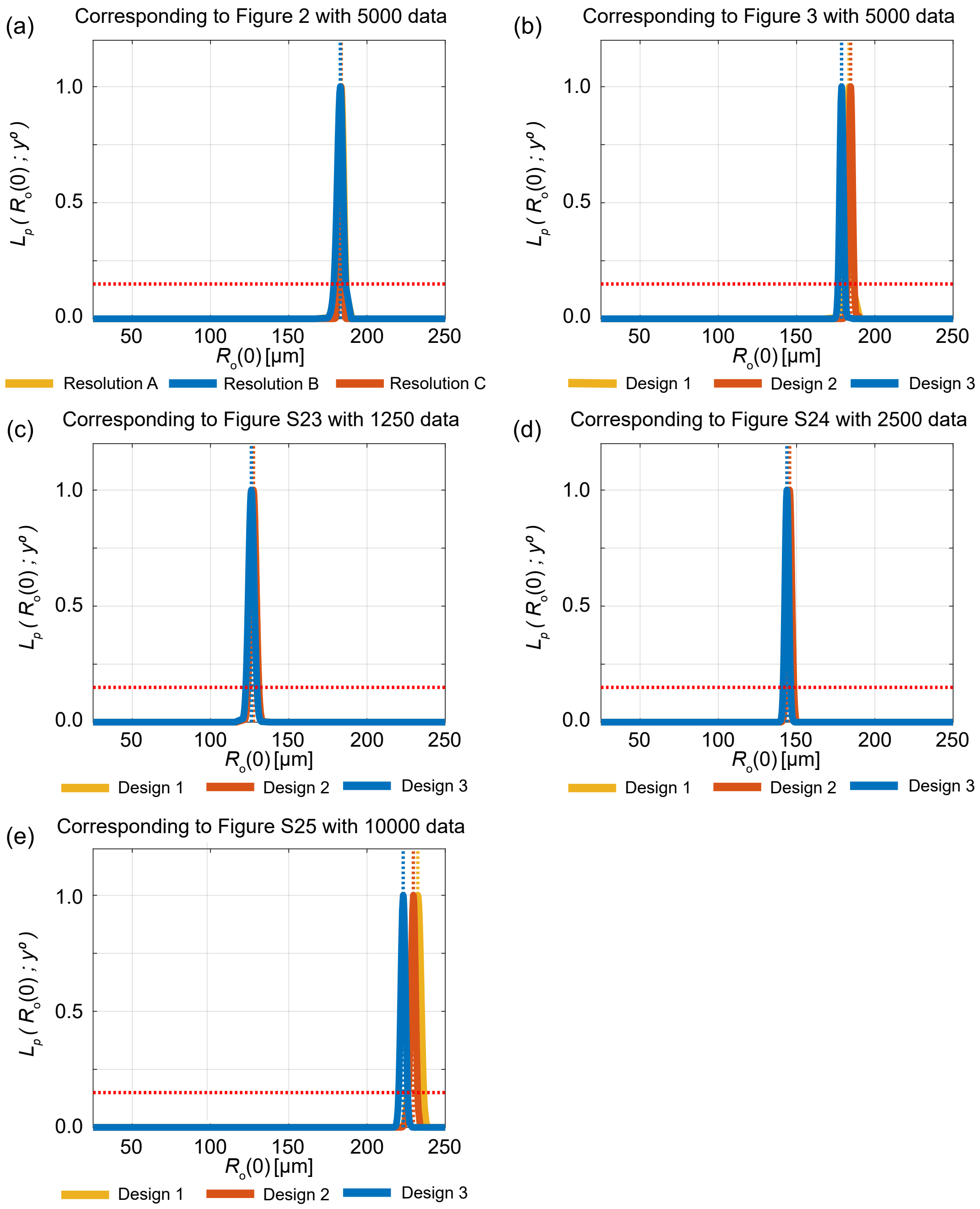


Figure S23: Profile likelihoods for $R_o(0)$

#### D.4 Outer radius measurements are not sufficient to predict inhibited and necrotic radii

In Figure 2 we compare Design 1 with Temporal Resolutions A, B, and C for the WM793b cell line formed with 5000 cells. Here, in Figures S24, S25, and S26, we compare Design 1 with Temporal Resolutions A, B, and C for the WM793b spheroids formed with 1250, 2500, and 10000 cells, respectively. These results show that Design 1 is not a reliable design and that outer radius measurements are not sufficient to predict inhibited and necrotic radii.

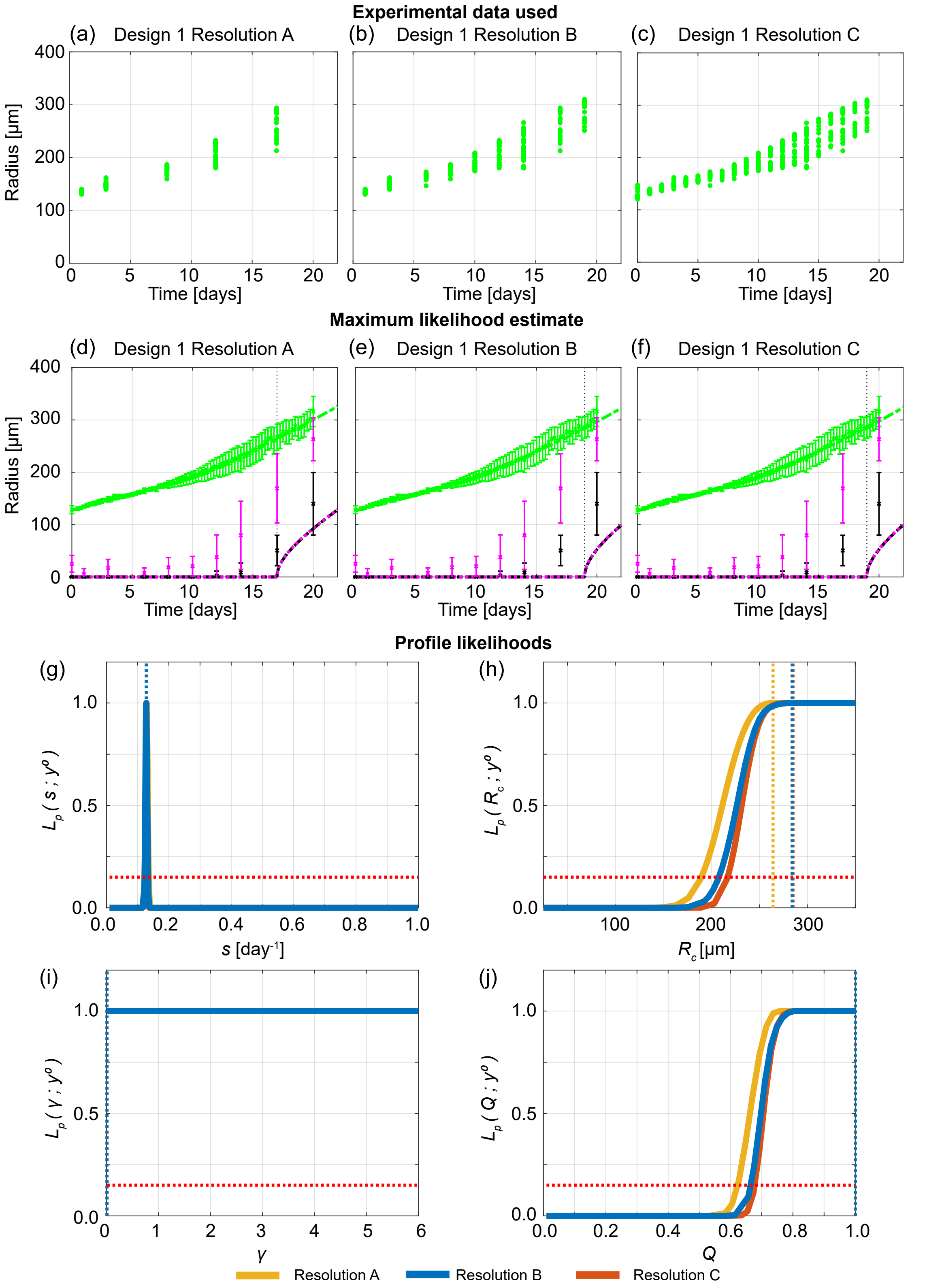


Figure S24: Increasing the temporal resolution when the outer radius is measured does not provide accurate information on internal structure. (a)-(c) Experimental data used in Design 1 with temporal resolutions A, B, and C. (d)-(f) Comparison of Greenspan model simulated with maximum likelihood estimate compared to full experimental data set, where error bars show standard deviation. Profile likelihoods for (g) $s$, (h) $R_c$, (i) $\gamma$, (j) $Q$. Yellow, blue, orange lines in (g)-(j) represent profile likelihoods from Design 1 with temporal resolutions A, B, and C, respectively, and the red-dashed line shows the approximate 95% confidence interval threshold. Results shown for WM793b spheroids formed with 1250 cells per spheroid.

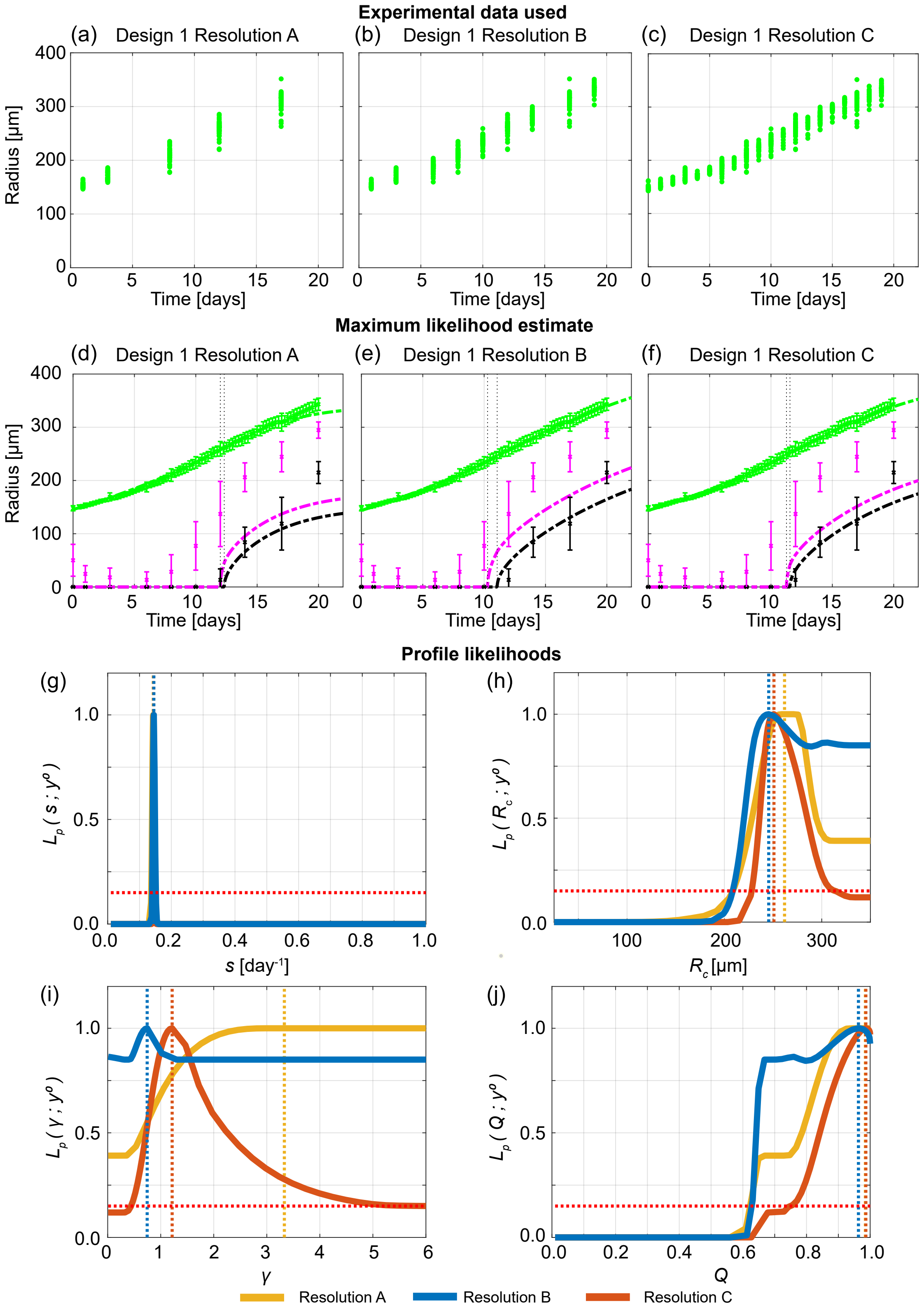


Figure S25: Increasing the temporal resolution when the outer radius is measured does not provide accurate information on internal structure. (a)-(c) Experimental data used in Design 1 with temporal resolutions A, B, and C. (d)-(f) Comparison of Greenspan model simulated with maximum likelihood estimate compared to full experimental data set, where error bars show standard deviation. Profile likelihoods for (g) $s$, (h) $R_c$, (i) $\gamma$, (j) $Q$. Yellow, blue, orange lines in (g)-(j) represent profile likelihoods from Design 1 with temporal resolutions A, B, and C, respectively, and the red-dashed line shows the approximate 95% confidence interval threshold. Results shown for WM793b spheroids formed with 2500 cells per spheroid.

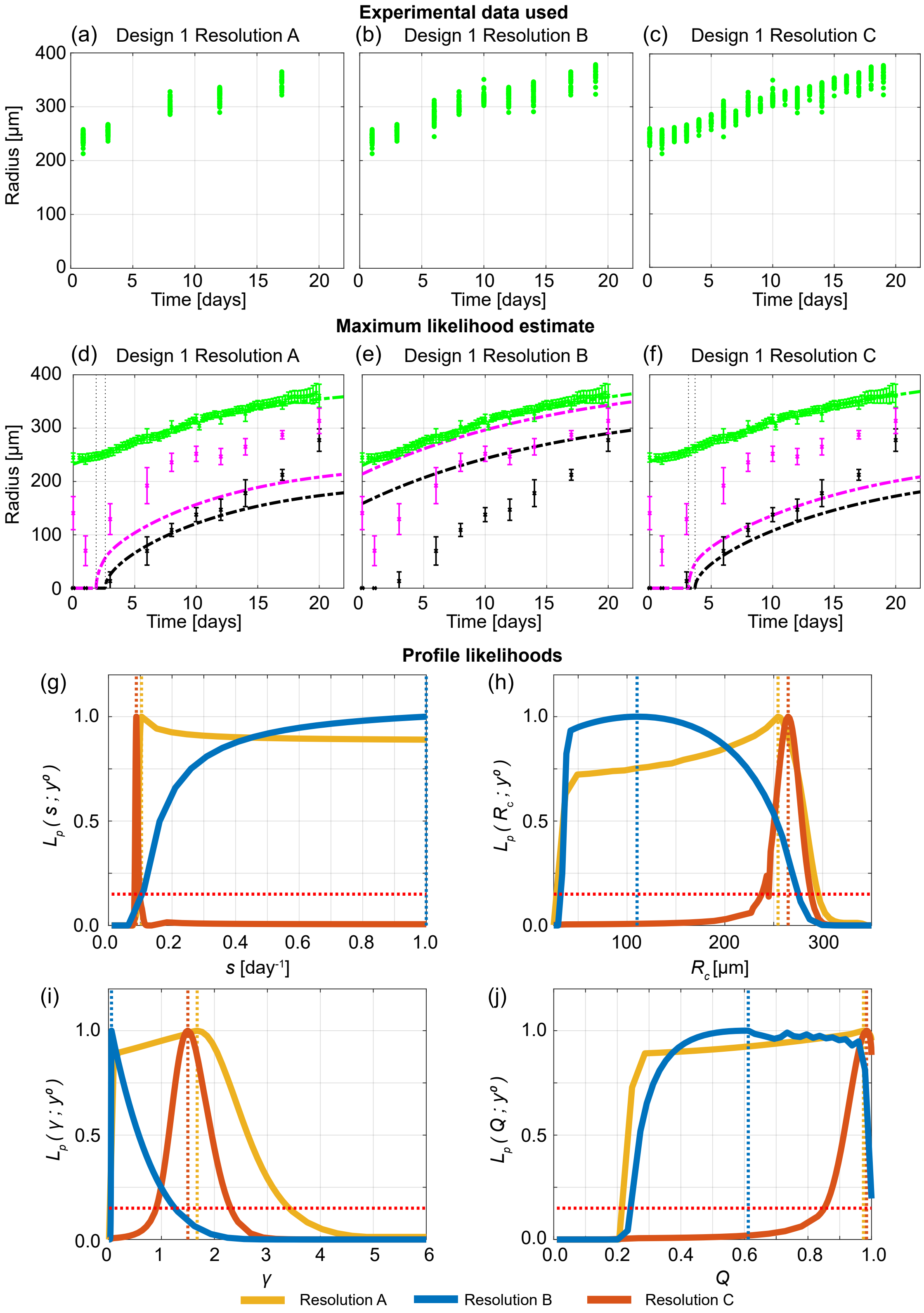


Figure S26: Increasing the temporal resolution when the outer radius is measured does not provide accurate information on internal structure. (a)-(c) Experimental data used in Design 1 with temporal resolutions A, B, and C. (d)-(f) Comparison of Greenspan model simulated with maximum likelihood estimate compared to full experimental data set, where error bars show standard deviation. Profile likelihoods for (g) $s$, (h) $R_c$, (i) $\gamma$, (j) $Q$. Yellow, blue, orange lines in (g)-(j) represent profile likelihoods from Design 1 with temporal resolutions A, B, and C, respectively, and the red-dashed line shows the approximate 95% confidence interval threshold. Results shown for WM793b spheroids formed with 10000 cells per spheroid.

#### D.5 Cell cycle and necrotic core measurements reveal time evolution of internal spheroid structure

In Figure 3 we compare Designs 1, 2, and 3 for the WM793b cell line formed with 5000 cells. Here, in Figures S27, S28, and S29, we compare Designs 1, 2, and 3 for the WM793b spheroids formed with 1250, 2500, and 10000 cells, respectively. These results also show that Design 3 provides most insight.

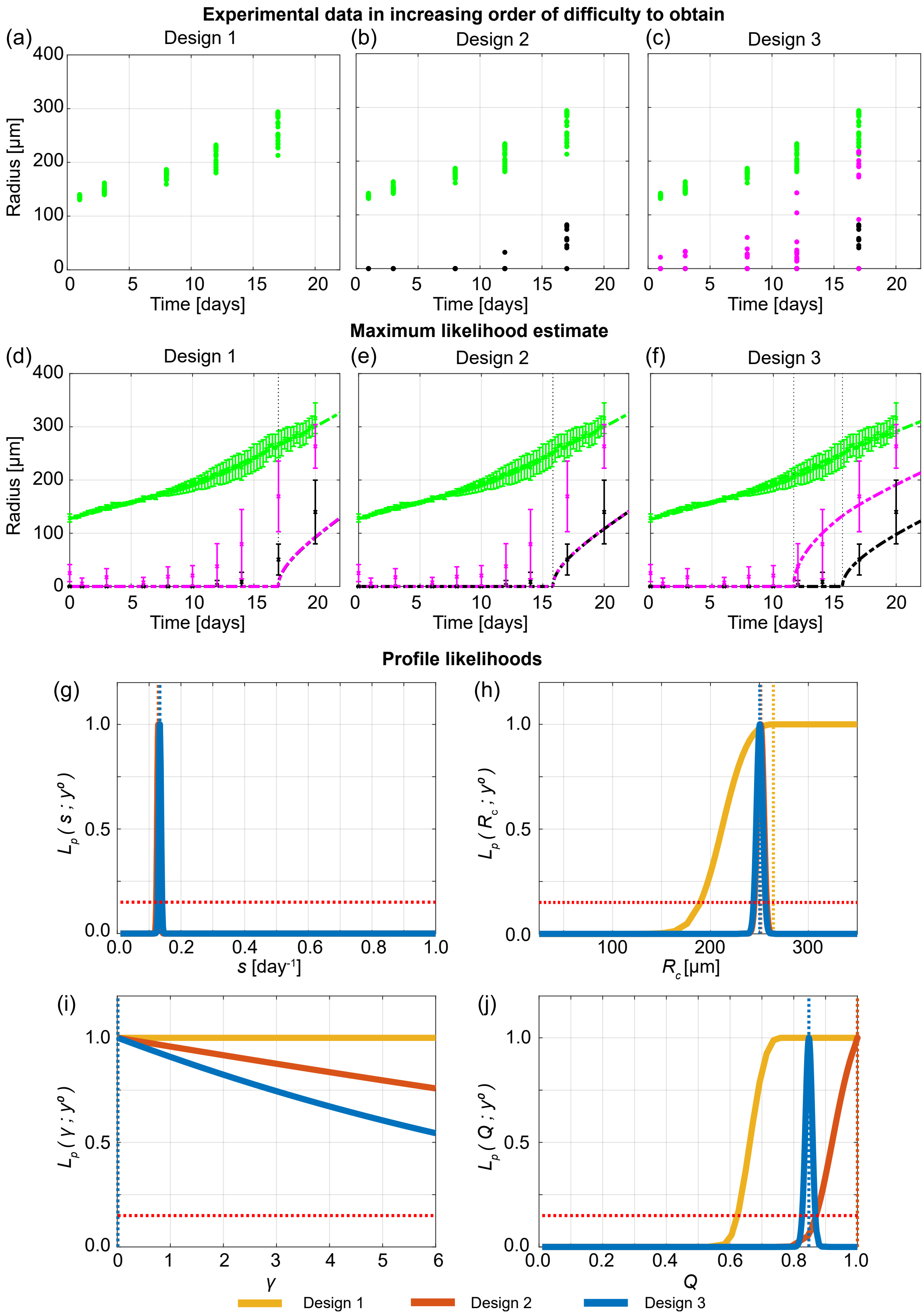


Figure S27: Measuring the necrotic and inhibited radius provides valuable information. (a)-(c) Experimental data used in Designs 1, 2 and 3. (d)-(f) Comparison of Greenspan model simulated with maximum likelihood estimate compared to full experimental data set for Designs 1, 2 and 3, where error bars show standard deviation. Profile likelihoods for (g) $s$, (h) $R_c$, (i) $\gamma$, (j) $Q$. Yellow, orange, blue lines in (g)-(j) represent profile likelihoods from Designs 1, 2, and 3, respectively, and the red-dashed line shows the approximate 95% confidence interval threshold. Results shown for WM793b spheroids formed with 1250 cells per spheroid.

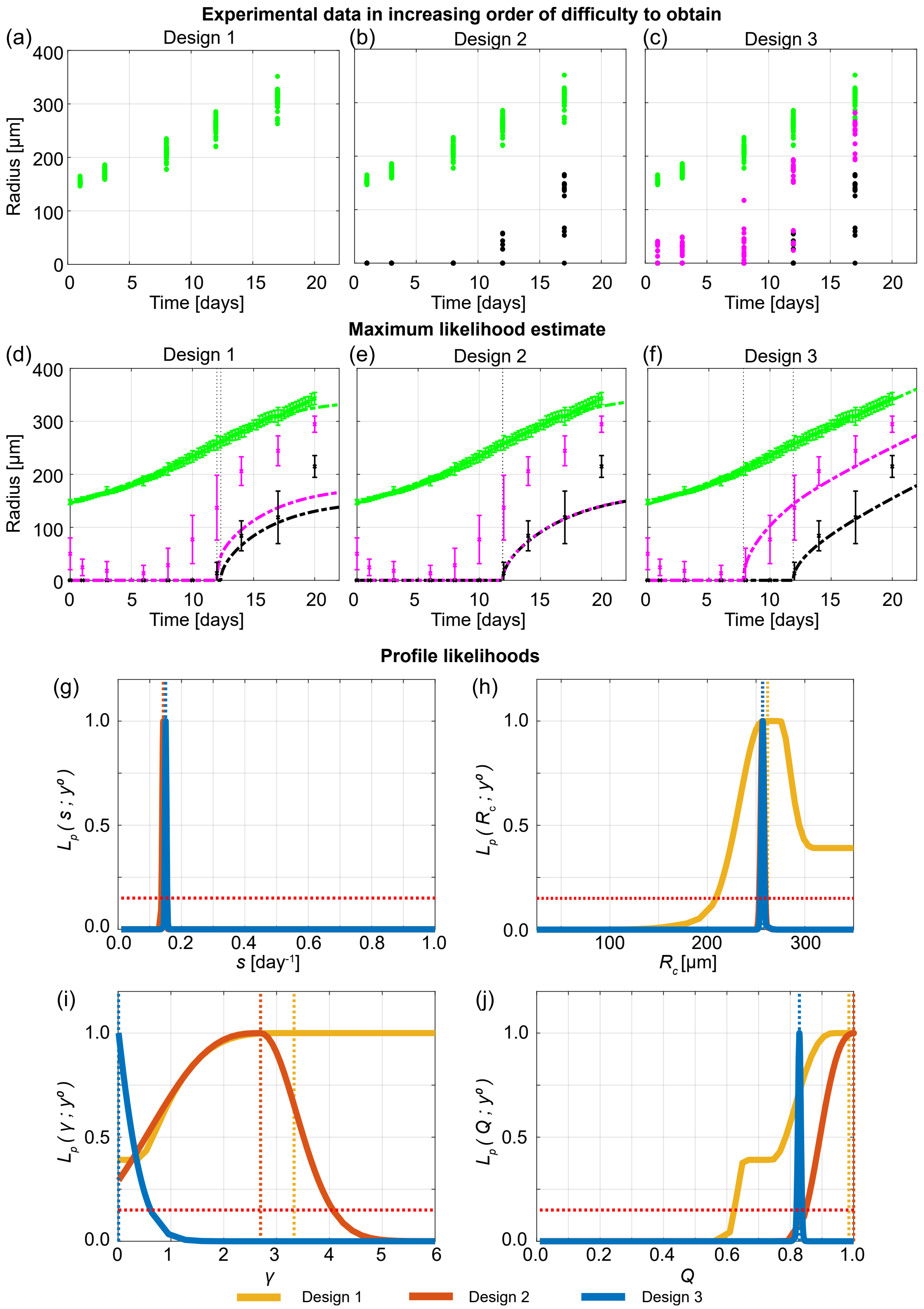


Figure S28: Measuring the necrotic and inhibited radius provides valuable information. (a)-(c) Experimental data used in Designs 1, 2 and 3. (d)-(f) Comparison of Greenspan model simulated with maximum likelihood estimate compared to full experimental data set for Designs 1, 2 and 3, where error bars show standard deviation. Profile likelihoods for (g) $s$, (h) $R_c$, (i) $\gamma$, (j) $Q$. Yellow, orange, blue lines in (g)-(j) represent profile likelihoods from Designs 1, 2, and 3, respectively, and the red-dashed line shows the approximate 95% confidence interval threshold. Results shown for WM793b spheroids formed with 2500 cells per spheroid.

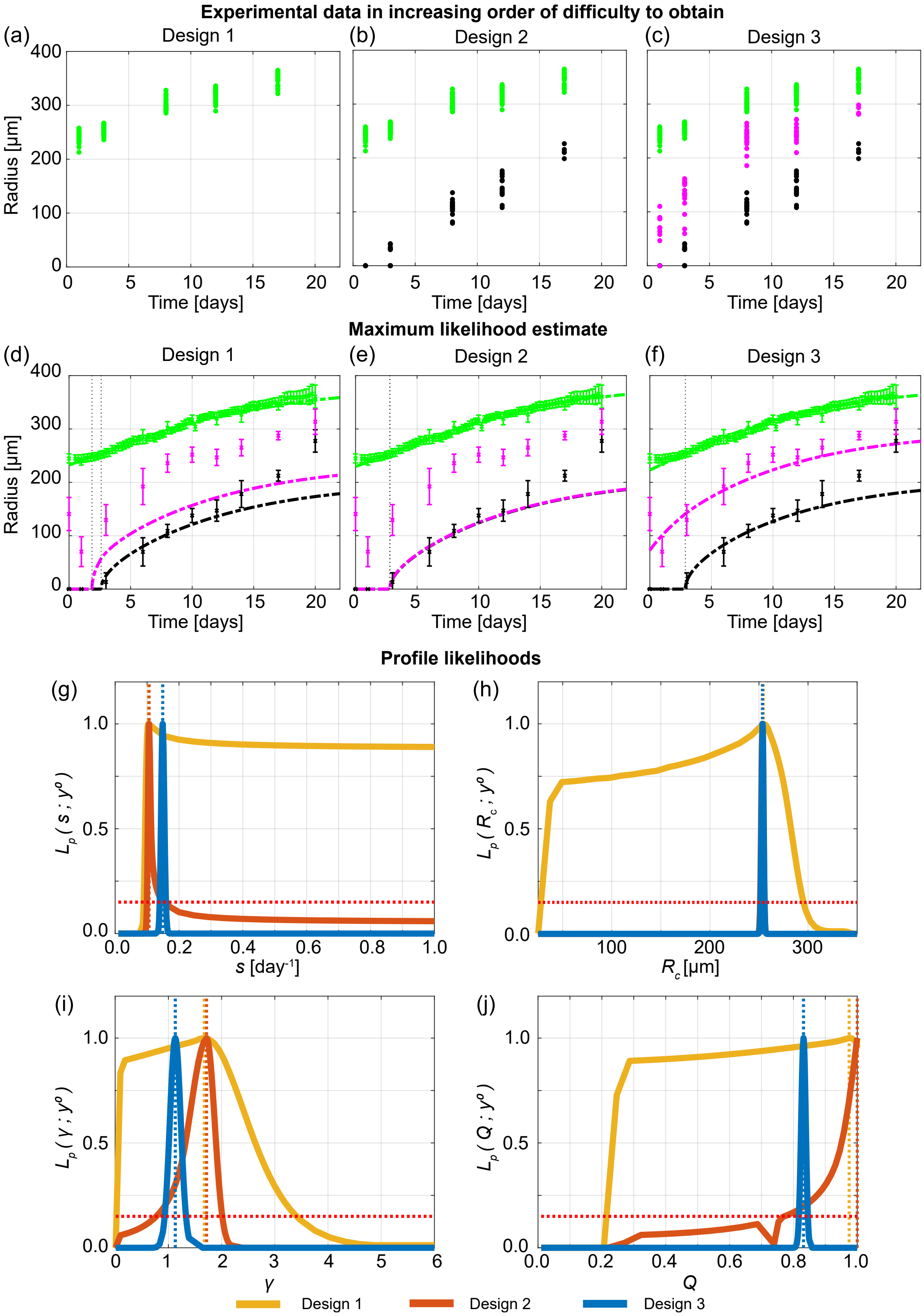


Figure S29: Measuring the necrotic and inhibited radius provides valuable information. (a)-(c) Experimental data used in Designs 1, 2 and 3. (d)-(f) Comparison of Greenspan model simulated with maximum likelihood estimate compared to full experimental data set for Designs 1, 2 and 3, where error bars show standard deviation. Profile likelihoods for (g) $s$, (h) $R_c$, (i) $\gamma$, (j) $Q$. Yellow, orange, blue lines in (g)-(j) represent profile likelihoods from Designs 1, 2, and 3, respectively, and the red-dashed line shows the approximate 95% confidence interval threshold. Results shown for WM793b spheroids formed with 10000 cells per spheroid.

### E Synthetic data: WM793b

To confirm that profile likelihood analysis works as expected, we generate synthetic data from Greenspan's mathematical model using known parameters. We then explore when these known parameters are recovered using the varying experimental designs considered in the main manuscript: Design 1 with varying temporal resolutions (Supplementary Material E.1); comparing Design 1, Design 2 and Design 3 (Supplementary Material E.2), and exploring the role of initial spheroid size and also here experimental duration (Supplementary Material E.3). Since Greenspan's model may be misspecified, and may not capture all of the biological details of tumour spheroid growth, the fact that these results for synthetic data are consistent with those from experimental data enhances our confidence that key biological features are captured in Greenspan's model. Furthermore, when generating synthetic data there is additional flexibility so we also explore what may happen if we were to spend significantly more time collecting measurements (Supplementary Material E.4).

To generate synthetic data, we use the MLE from Design 3 Resolution C applied to experimental data obtained from WM793b spheroids each formed with 5000 cells: $(R_\mathrm{c}, s, \gamma, Q, R_\mathrm{o}(0)) = (254.366, 0.1532, 0.045, 0.797, 179.550)$. First, we simulate Greenspan's deterministic mathematical model with these known parameters. Next, to obtain one noisy synthetic outer radius measurement we record the outer radius from Greenspan's model generated from the known parameters at one time point. Next, we sample a normal distribution with zero mean and variance given by experimentally obtained outer radius pooled sample variance $s_\mathrm{o}^2 = 9.35$. We add this sampled noise to the recorded outer radius measurement. We repeat this process to obtain additional outer radius measurements. Similarly, we repeat this process to obtain necrotic and inhibited radius measurements, using experimentally obtained pooled sample variances $s_\mathrm{n}^2 = 15.89$, and $s_\mathrm{i}^2 = 33.12$, respectively. We generate 10 measurements, or 48 when exploring the role of additional measurements in supplementary material E.4, of the outer radius, inhibited radius and necrotic radius every half day from day 0 to day 20.

#### E.1 Outer radius measurements are not sufficient to predict inhibited and necrotic radii

Similarly to Figure 2, we observe in Figure S30 that outer radius measurements are not sufficient to predict inhibited and necrotic radii. Simulating Greenspan's model with the MLE from Design 1 Time Resolution A (Figure S30d) and with Design 1 Time Resolution B (Figure S30e) shows the time evolution of the outer radius is captured but the time evolution of the inhibited and necrotic region are not. However, simulating Greenspan's model with the MLE from Design 1 Time Resolution C (Figure S30f) appears to capture the time evolution of the outer, inhibited, and necrotic radii. However, inspecting the profile likelihoods in Figures S30g-j shows that, while the known parameters are captured, the profiles are wide suggesting that parameters are non-identifiable. This means that many parameter values give a similar match to the outer radius experimental data and these parameters do not necessarily agree with the inhibited and necrotic radii measurements.

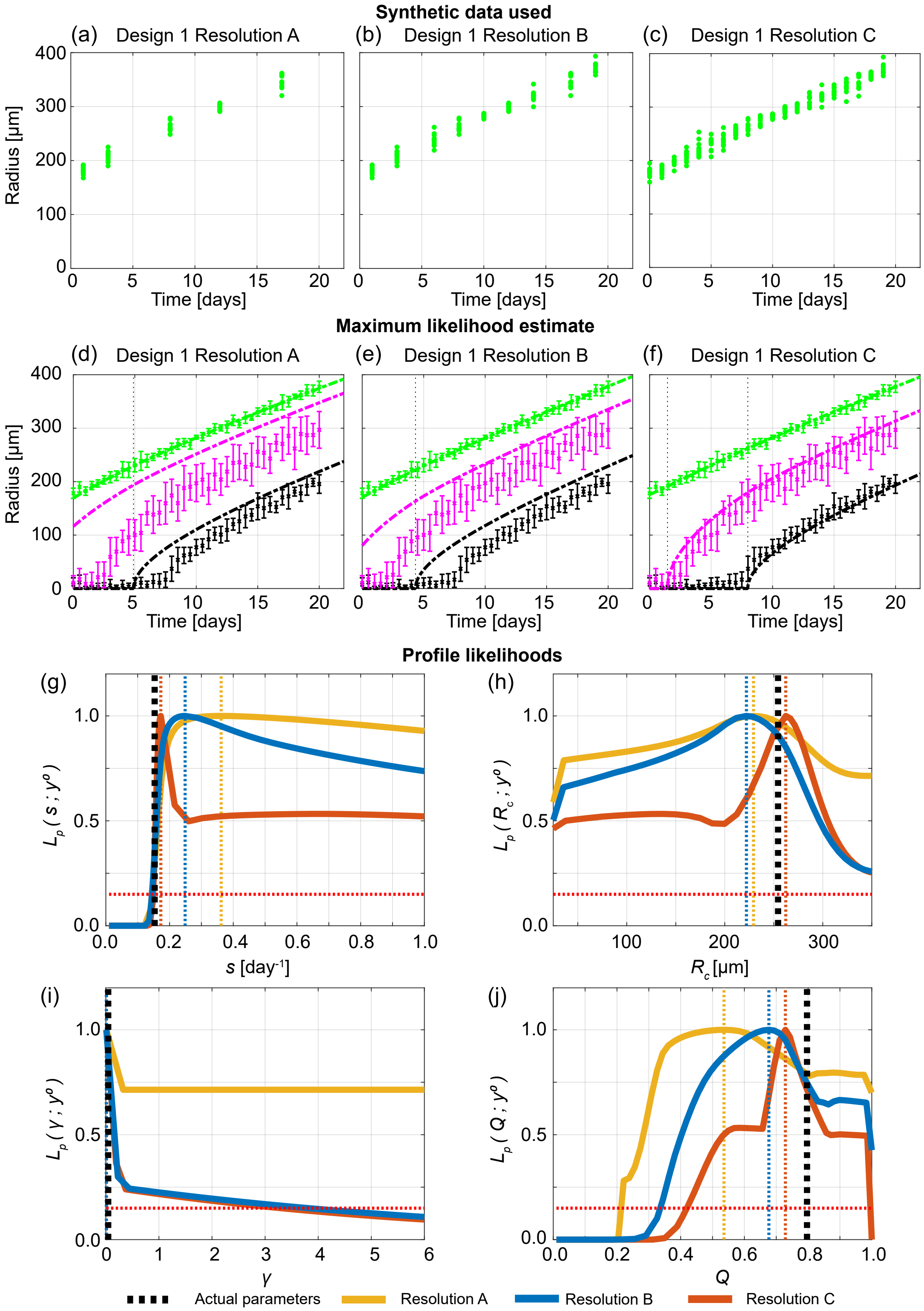


Figure S30: Synthetic data shows that outer radius measurements are not sufficient to predict inhibited and necrotic radii. (a)-(c) Synthetic data used in Designs 1 with Temporal Resolutions A, B, and C. (d)-(f) Comparison of Greenspan model simulated with maximum likelihood estimate compared to full synthetic data set. Profile likelihoods for (g) $s$, (h) $R_c$, (i) $\gamma$, (j) $Q$. Yellow, blue, and orange lines in (g)-(j) represent profile likelihoods from Design 1 with Temporal Resolutions A, B, and C, respectively. Black dashed lines in (g)-(j) show known parameters used to generate the synthetic data.

#### E.2 Cell cycle and necrotic core measurements reveal time evolution of internal spheroid structure

Similarly to Figure 3, Design 3 provides most insight and best captures the known parameter values used to generate the synthetic data (Figure S31).

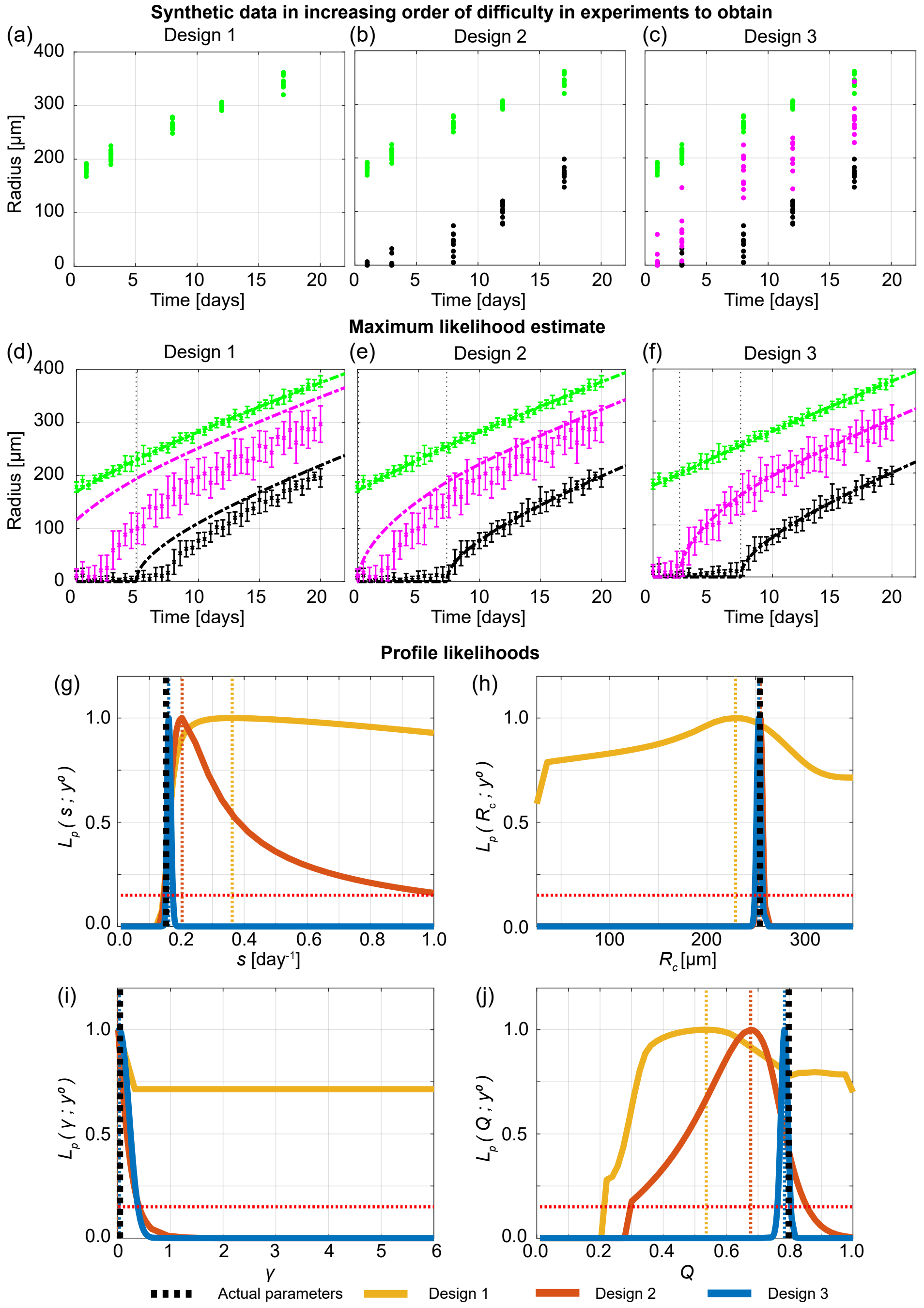


Figure S31: Synthetic data shows that measuring the necrotic and inhibited radius provides valuable information. (a)-(c) Synthetic data used in Designs 1, 2 and 3. (d)-(f) Comparison of Greenspan model simulated with maximum likelihood estimate compared to full synthetic data set for Designs 1, 2 and 3. Profile likelihoods for (g) $s$, (h) $R_c$, (i) $\gamma$, (j) $Q$. Yellow, orange, blue lines in (g)-(j) represent profile likelihoods from Designs 1 low temporal resolution, 2, and 3, respectively. Black dashed lines in (e)-(h) show known parameters used to generate the synthetic data.

#### E.3 Role of initial spheroid size and experiment duration

In Greenspan's model a change in the initial radius, $R_\mathrm{o}(0)$, corresponds to a shift in time (Figure S32a). We now consider the role of initial spheroid size and experiment duration. As before, we use the MLE from spheroids formed with 5000 cells per spheroid to generate synthetic data. To generate synthetic data for spheroids formed with 1250, 2500, and 10000 cells per spheroid we use the MLE obtained from spheroids formed with 5000 cells per spheroid and only update $R_\mathrm{o}(0)$. To update $R_\mathrm{o}(0)$ we use the MLE from Design 3 applied to experimental data obtained from WM793b spheroids formed with 1250, 2500, and 10000 cells per spheroid, respectively.

We assume that each experiment is performed to Day 6 after formation, and use Design 3 with 10 measurements obtained on Day 1, 2, 3, 4, 5, and 6 (Figure S32b-e). Note that during this experimental duration only spheroids formed with 10000 cells per spheroid form a necrotic core with the known parameters, while only spheroids formed with 5000 and 10000 cells per spheroid form an inhibited region with the known parameters. Therefore, we expect that most insight will be gained from the experiment formed with spheroids formed with 10000 cells per spheroid.

Simulating Greenspan's model with the MLE obtained from each of those data sets (dashed lines in Figures S32f-i) we observe good agreement to the first six days of synthetic data for each initial spheroid size. However, simulating Greenspan's model with the MLE obtained from each of those data sets (dashed lines in Figures S32f-i) and comparing to Greenspan's model simulated over 20 days with the known parameters used to generate the synthetic data (solid lines in Figures S32f-i)) this is not the case. We only observe excellent agreement for the experiment with spheroids formed with 10000 cells, since this experiment has measurements in phase (iii). Profile likelihoods for the parameters also show that only the experiment performed with spheroids formed with 10000 cells accurately captures the known parameters (Figure S32j-n).

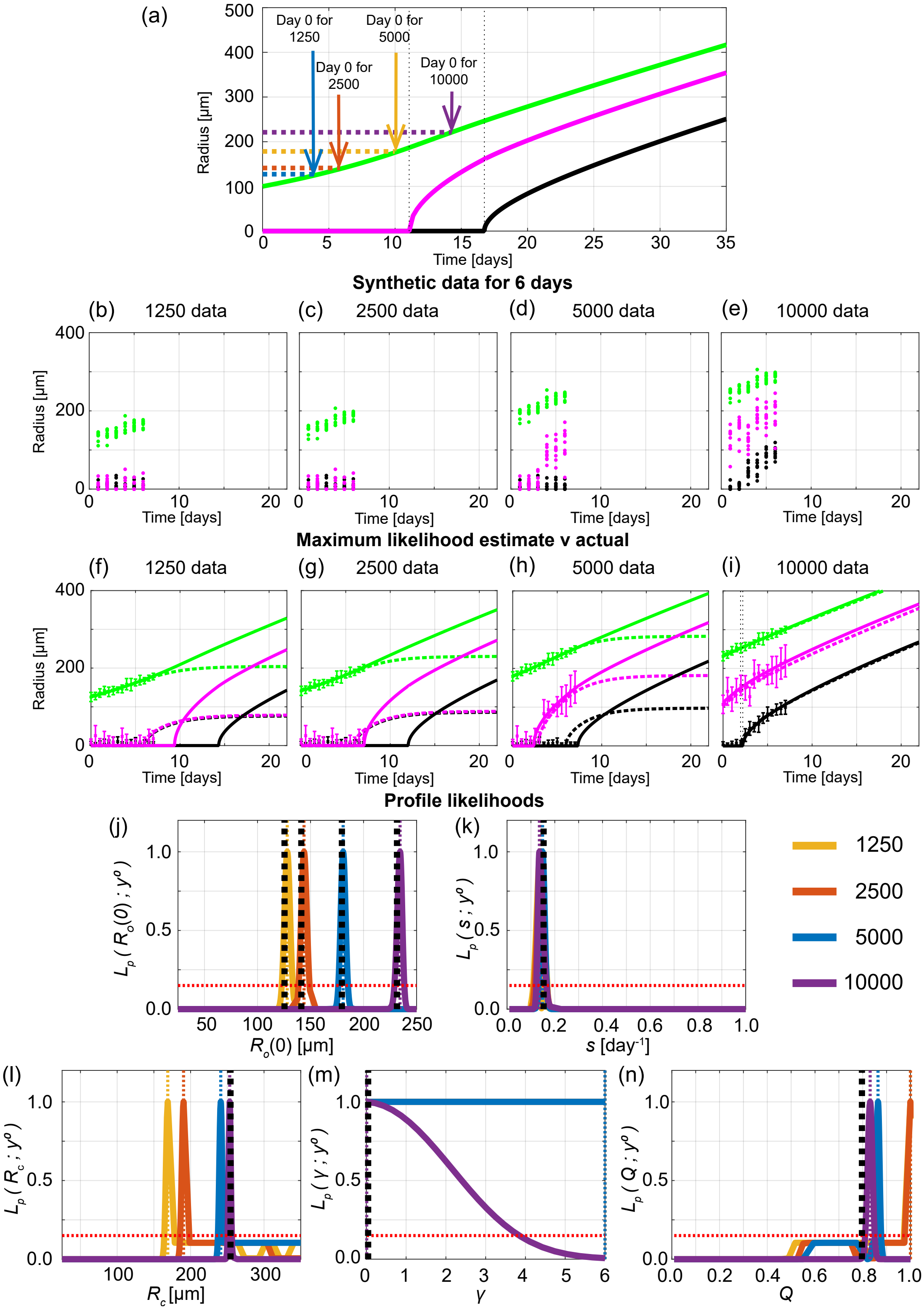


Figure S32: Synthetic data exploring role of initial spheroid size and experimental duration. (a) In Greenspan's model a change in $R_{\mathrm{o}}(0)$ corresponds to a shift in time. (b)-(e) Synthetic generated for the first six days after formation for spheroids formed with (b) 1250, (c) 2500, (d) 5000, (e) 10000 cells per spheroid. (f)-(i) Comparison of Greenspan model simulated with maximum likelihood estimate (dashed lines) compared to synthetic data for first 6 days compared to Greenspan model simulated with known parameters used to generate the synthetic data (solid lines). Profile likelihoods for (j) $R_{\mathrm{o}}(0)$, (k) $s$, (l) $R_{\mathrm{c}}$, (m) $\gamma$, (n) $Q$. Yellow, orange, blue, and purple lines in (j)-(n) represent profile likelihoods from spheroids formed with 1250, 2500, 5000, and 10000 cells per spheroid, respectively. Black dashed lines in (j)-(n) show known parameters used to generate the synthetic data.

#### E.4 Increasing number of measurements

In biological experiments it is time consuming and expensive to increase the number of measurements obtained. However, by generating synthetic data we can easily simulate additional measurements. We generate 48 measurements of the outer radius, inhibited radius, and necrotic radius every half day from Day 0 to Day 20. We choose 48 measurements since this corresponds to half of a 96-well plate and is extremely large in comparison to typical experiments. These results show that many measurements of Design 2 may provide good insight in this extreme scenario but that Design 3 still provides most insight.

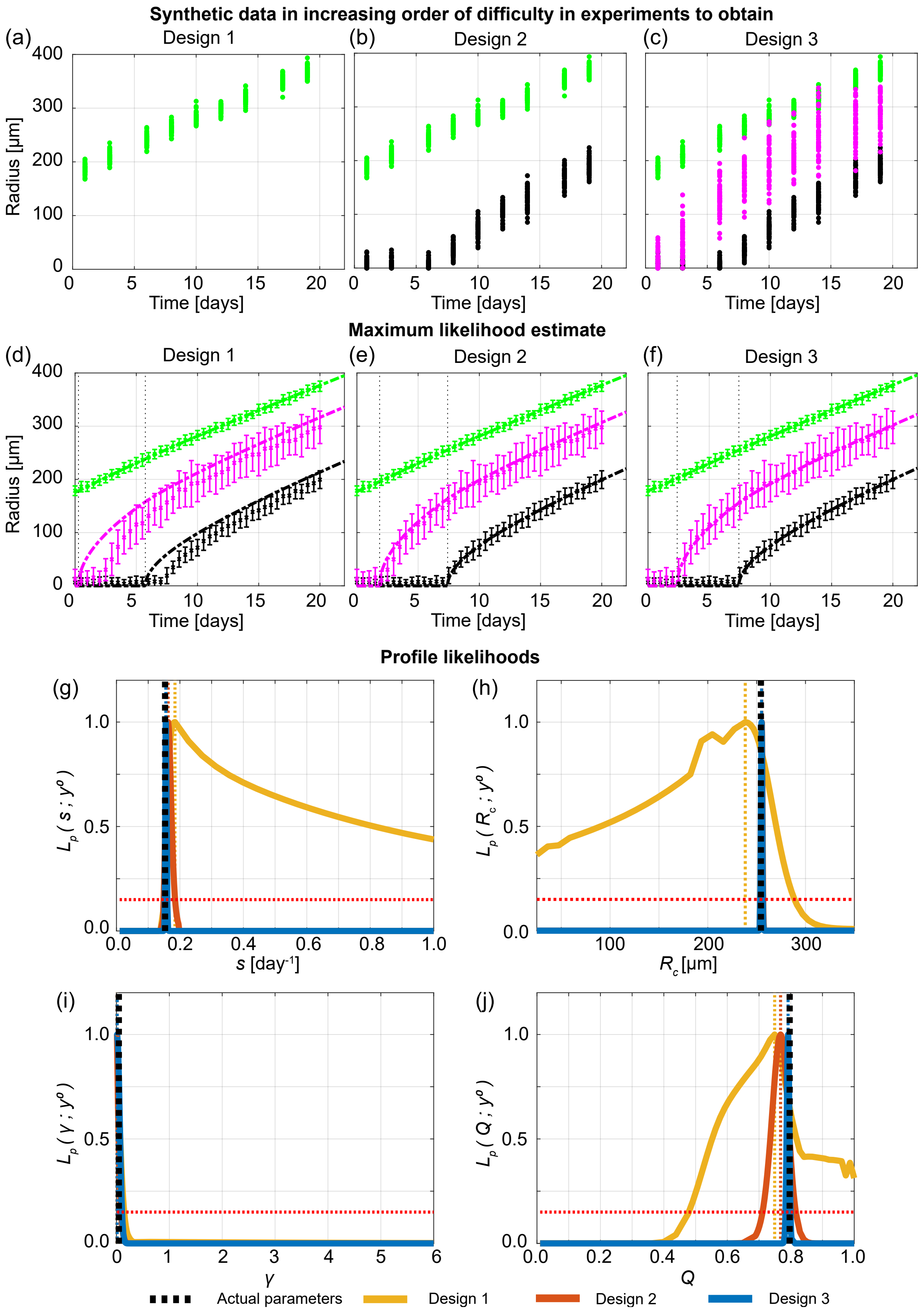


Figure S33: Synthetic data shows that more measurements of the necrotic and inhibited radius provides valuable information. (a)-(c) Synthetic data used in Designs 1, 2 and 3. (d)-(f) Comparison of Greenspan model simulated with maximum likelihood estimate compared to full synthetic data set for Designs 1, 2 and 3. Profile likelihoods for (g) $s$, (h) $R_c$, (i) $\gamma$, (j) $Q$. Yellow, orange, blue lines in (e)-(h) represent profile likelihoods from Designs 1, 2, and 3, with low temporal resolution, respectively. Black dashed lines in (g)-(j) show known parameters used to generate the synthetic data.

### F Parameter identifiability analysis for WM983b

The main manuscript focuses on results for the human melanoma WM793b cell line. Here, we present the corresponding results for the human melanoma WM983b spheroids formed with 2500, 5000, and 10000 cells.

All key observations made in reference to the WM793b cell line hold for the WM983b cell line. Specifically, in Figures S34, S35 and S36, for spheroids formed with 2500, 5000, and 10000 cells, respectively, we show that varying the temporal resolution using only Design 1 is insufficient to determine necrotic and inhibited radii. In Figures S37, S38, and S39 for spheroids formed with 2500, 5000, and 10000 cells, respectively, we show that Design 3 provides most insight. In Figure S40 we show that information gained across experiments with different initial spheroid sizes is consistent. Minor modifications were applied to the experimental designs as the WM983b tumour spheroids form after 3 days, which is 1 day earlier than the WM793b tumour spheroids, and the experiment was terminated after 19 days so the updated temporal resolutions are for this cell line are chosen as: Resolution A using Days 1, 3, 8, 12, 16; Resolution B using Days 1, 3, 5, 8, 10, 12, 14, 16, 18; Resolution C using Days 0, 1, 2, 3, 4, 5, 6, 7, 8, 9, 10, 11, 12, 13, 14, 15, 16, 17, 18; where Day 0 corresponds to the time that we determined as when spheroid formation ends and growth begins.

#### F.1 Outer radius measurements are not sufficient to predict inhibited and necrotic radii

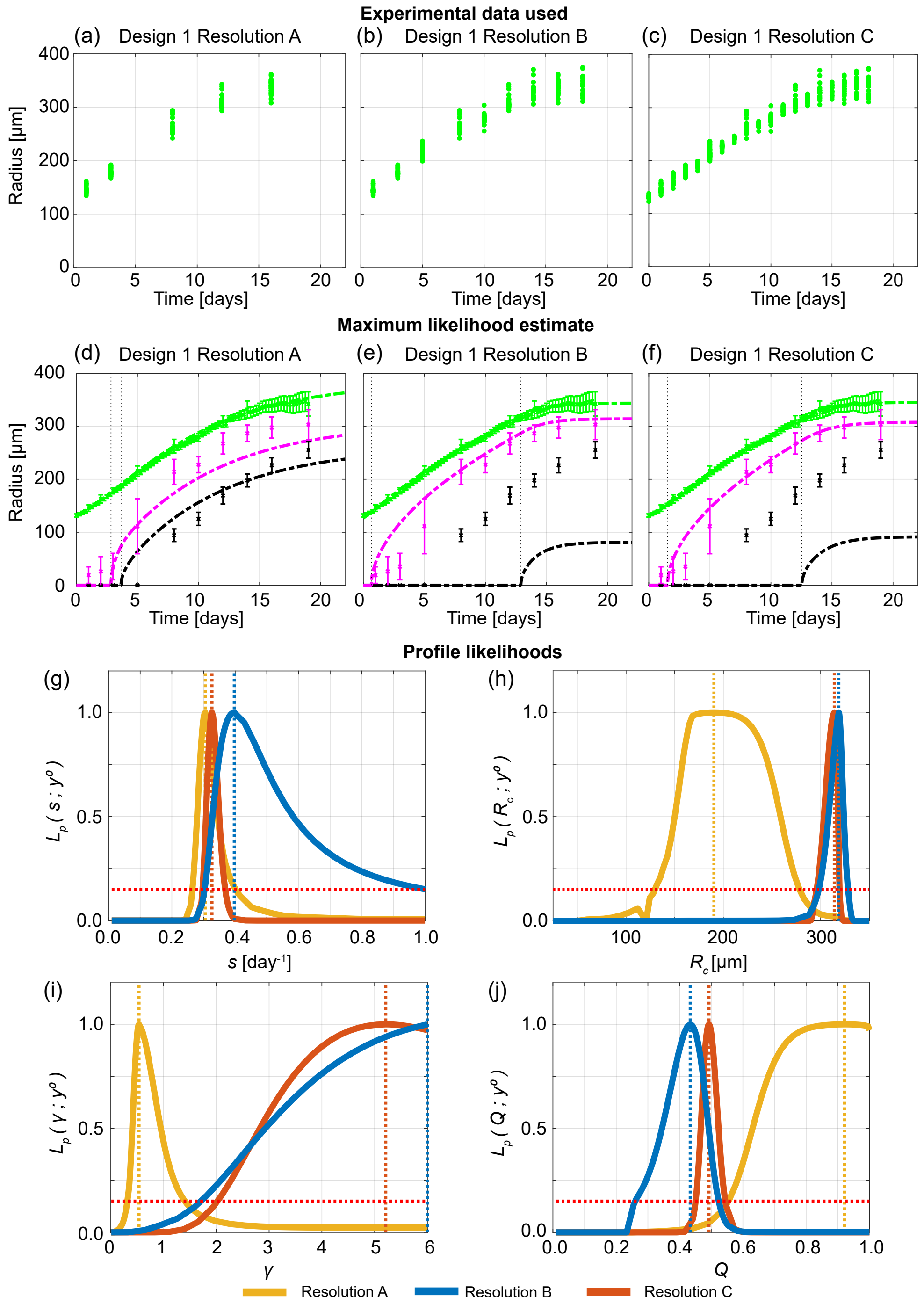


Figure S34: Increasing the temporal resolution when the outer radius is measured does not provide accurate information on internal structure. (a)-(c) Experimental data used in Design 1 with temporal resolutions A, B, and C. (d)-(f) Comparison of Greenspan model simulated with maximum likelihood estimate compared to full experimental data set, where error bars show standard deviation. Profile likelihoods for (g) $s$, (h) $R_c$, (i) $\gamma$, (j) $Q$. Yellow, blue, orange lines in (g)-(j) represent profile likelihoods from Design 1 with temporal resolutions A, B, and C, respectively, and the red-dashed line shows the approximate 95% confidence interval threshold. Results shown for WM983b spheroids formed with 2500 cells per spheroid.

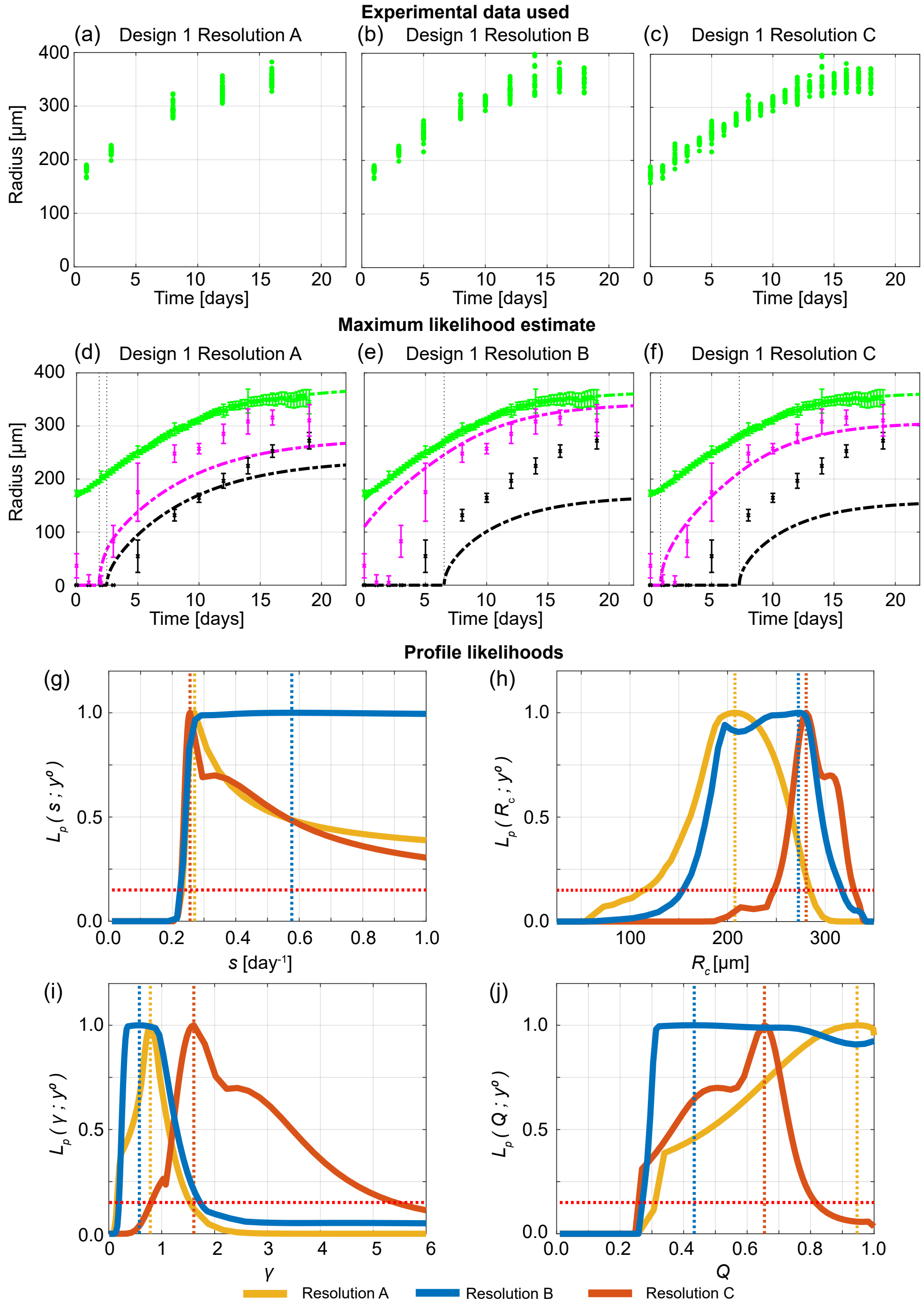


Figure S35: Increasing the temporal resolution when the outer radius is measured does not provide accurate information on internal structure. (a)-(c) Experimental data used in Design 1 with temporal resolutions A, B, and C. (d)-(f) Comparison of Greenspan model simulated with maximum likelihood estimate compared to full experimental data set, where error bars show standard deviation. Profile likelihoods for (g) $s$, (h) $R_c$, (i) $\gamma$, (j) $Q$. Yellow, blue, orange lines in (g)-(j) represent profile likelihoods from Design 1 with temporal resolutions A, B, and C, respectively, and the red-dashed line shows the approximate 95% confidence interval threshold. Results shown for WM983b spheroids formed with 5000 cells per spheroid.

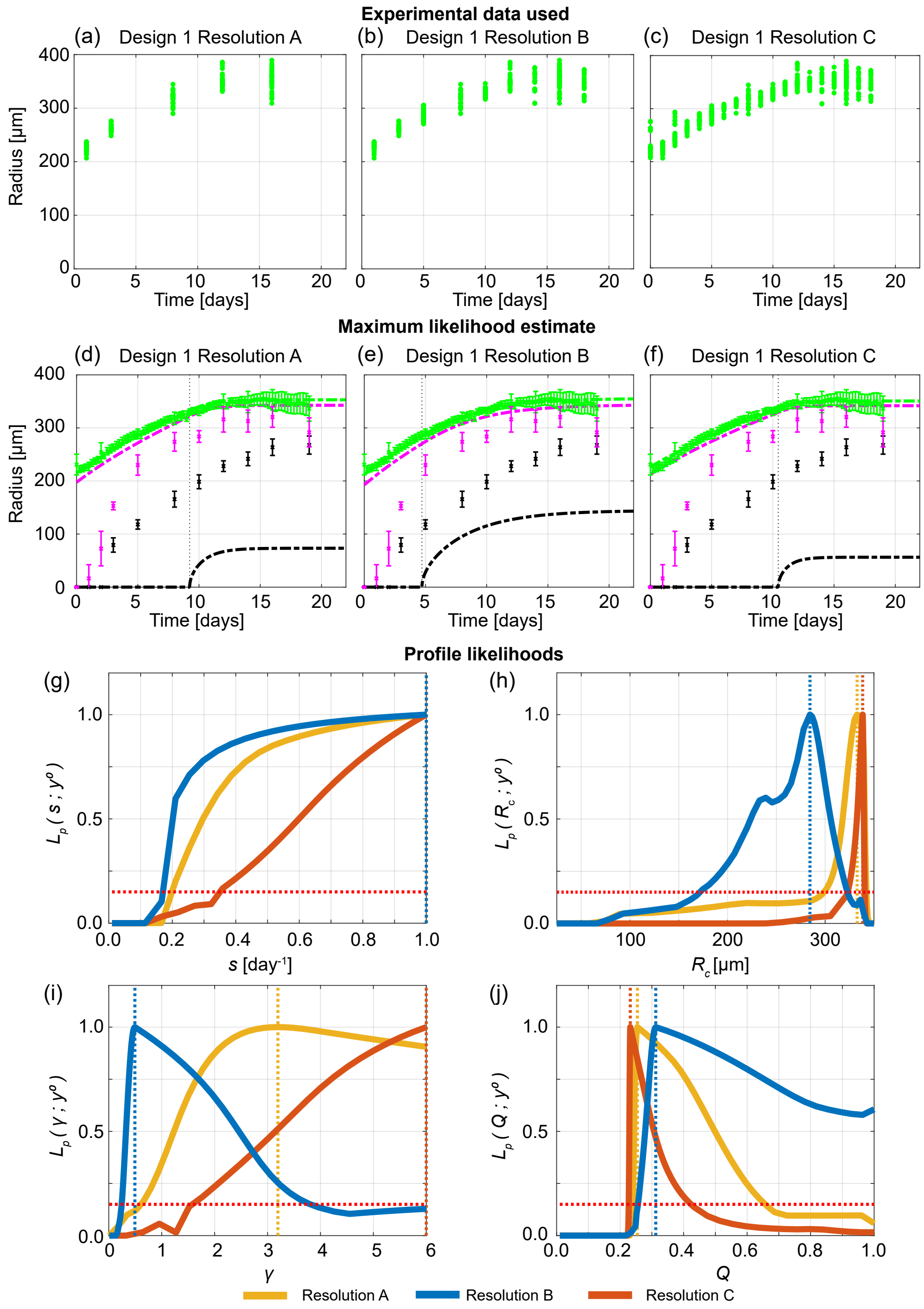


Figure S36: Increasing the temporal resolution when the outer radius is measured does not provide accurate information on internal structure. (a)-(c) Experimental data used in Design 1 with temporal resolutions A, B, and C. (d)-(f) Comparison of Greenspan model simulated with maximum likelihood estimate compared to full experimental data set, where error bars show standard deviation. Profile likelihoods for (g) $s$, (h) $R_c$, (i) $\gamma$, (j) $Q$. Yellow, blue, orange lines in (g)-(j) represent profile likelihoods from Design 1 with temporal resolutions A, B, and C, respectively, and the red-dashed line shows the approximate 95% confidence interval threshold. Results shown for WM983b spheroids formed with 10000 cells per spheroid.

#### F.2 Cell cycle and necrotic core measurements reveal time evolution of internal spheroid structure

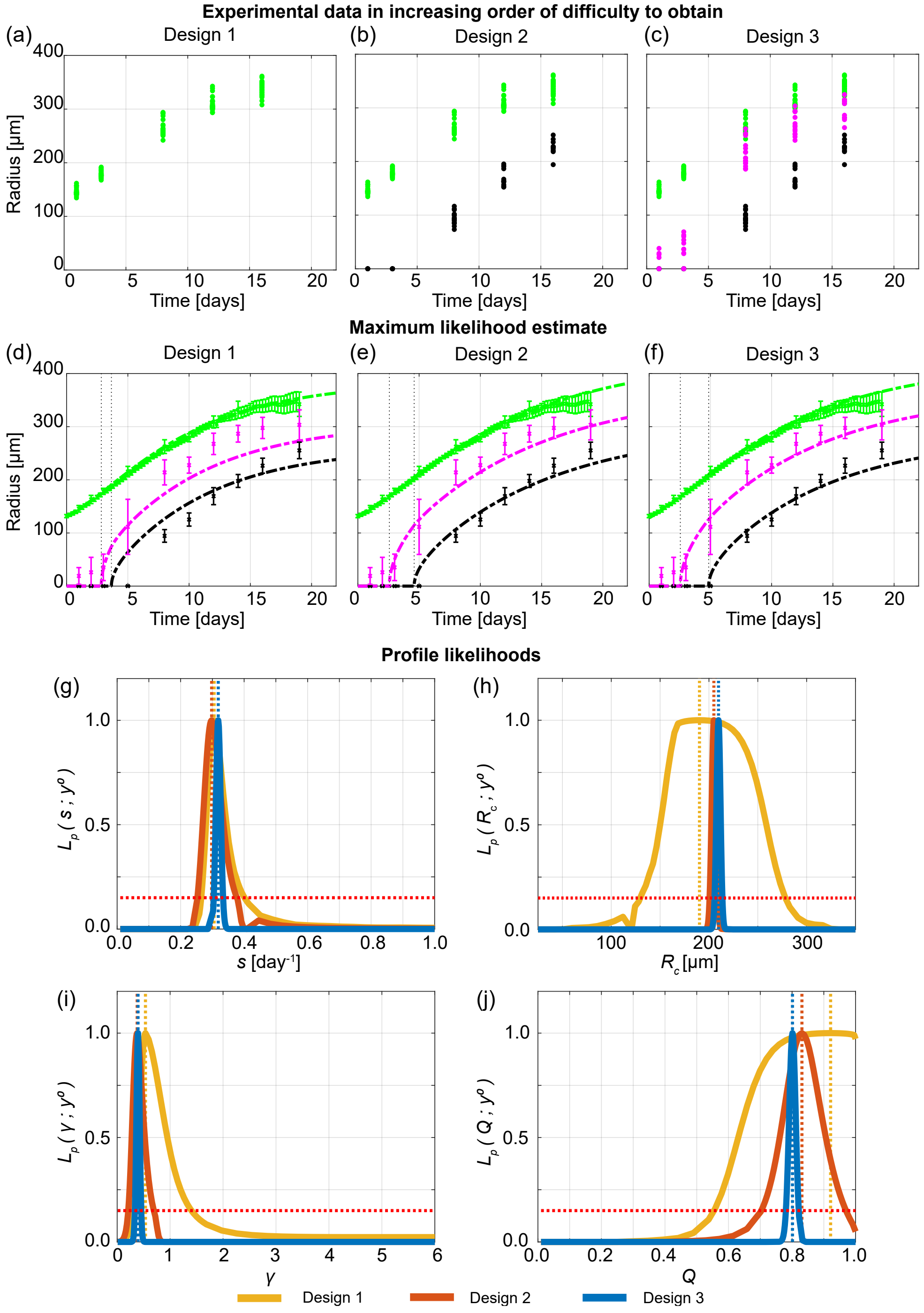


Figure S37: Measuring the necrotic and inhibited radius provides valuable information. (a)-(c) Experimental data used in Designs 1, 2 and 3. (d)-(f) Comparison of Greenspan model simulated with maximum likelihood estimate compared to full experimental data set for Designs 1, 2 and 3, where error bars show standard deviation. Profile likelihoods for (g) $s$, (h) $R_c$, (i) $\gamma$, (j) $Q$. Yellow, orange, blue lines in (g)-(j) represent profile likelihoods from Designs 1, 2, and 3, respectively, and the red-dashed line shows the approximate 95% confidence interval threshold. Results shown for WM983b spheroids formed with 2500 cells per spheroid.

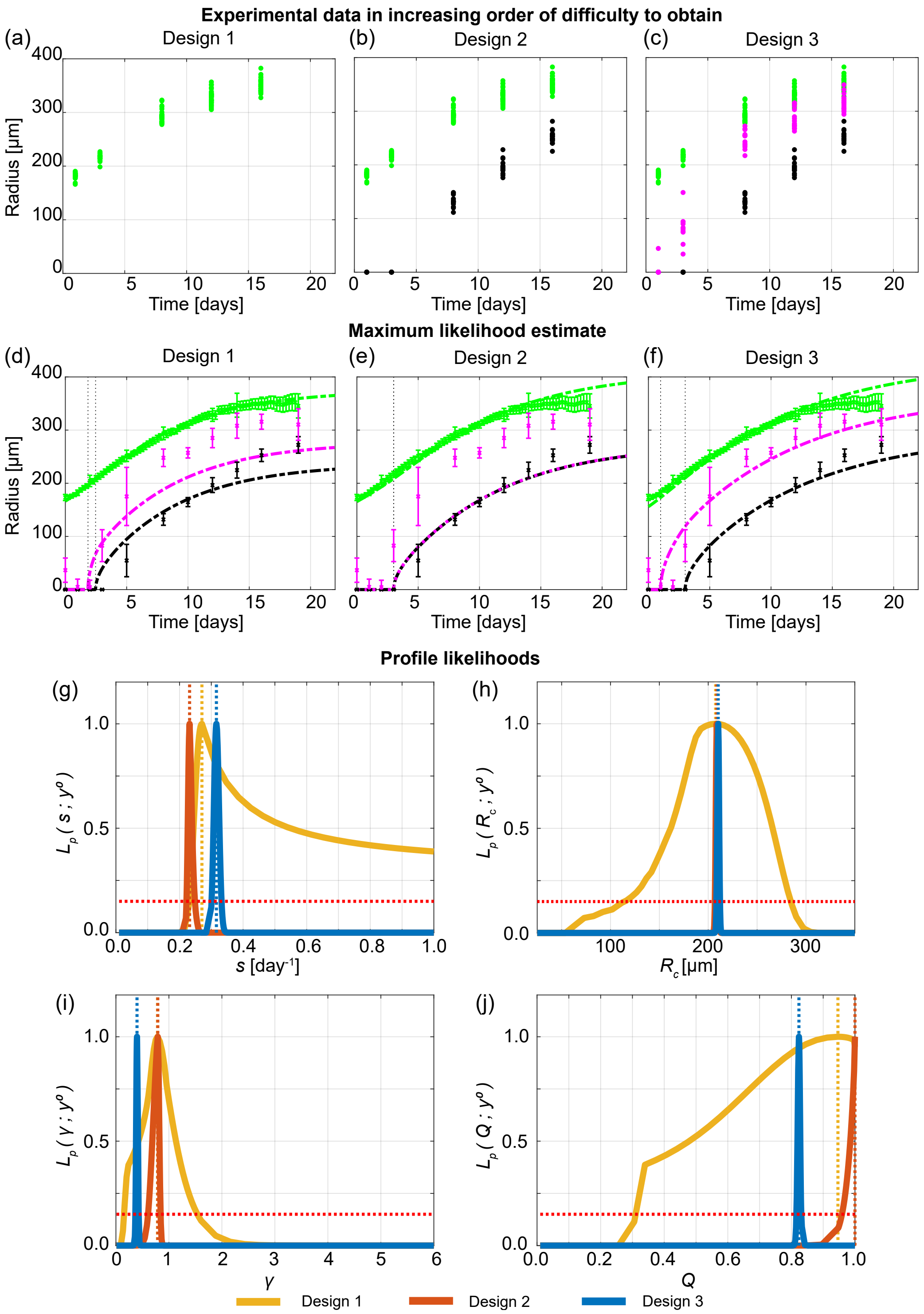


Figure S38: Measuring the necrotic and inhibited radius provides valuable information. (a)-(c) Experimental data used in Designs 1, 2 and 3. (d)-(f) Comparison of Greenspan model simulated with maximum likelihood estimate compared to full experimental data set for Designs 1, 2 and 3, where error bars show standard deviation. Profile likelihoods for (g) $s$, (h) $R_c$, (i) $\gamma$, (j) $Q$. Yellow, orange, blue lines in (g)-(j) represent profile likelihoods from Designs 1, 2, and 3, respectively, and the red-dashed line shows the approximate 95% confidence interval threshold. Results shown for WM983b spheroids formed with 5000 cells per spheroid.

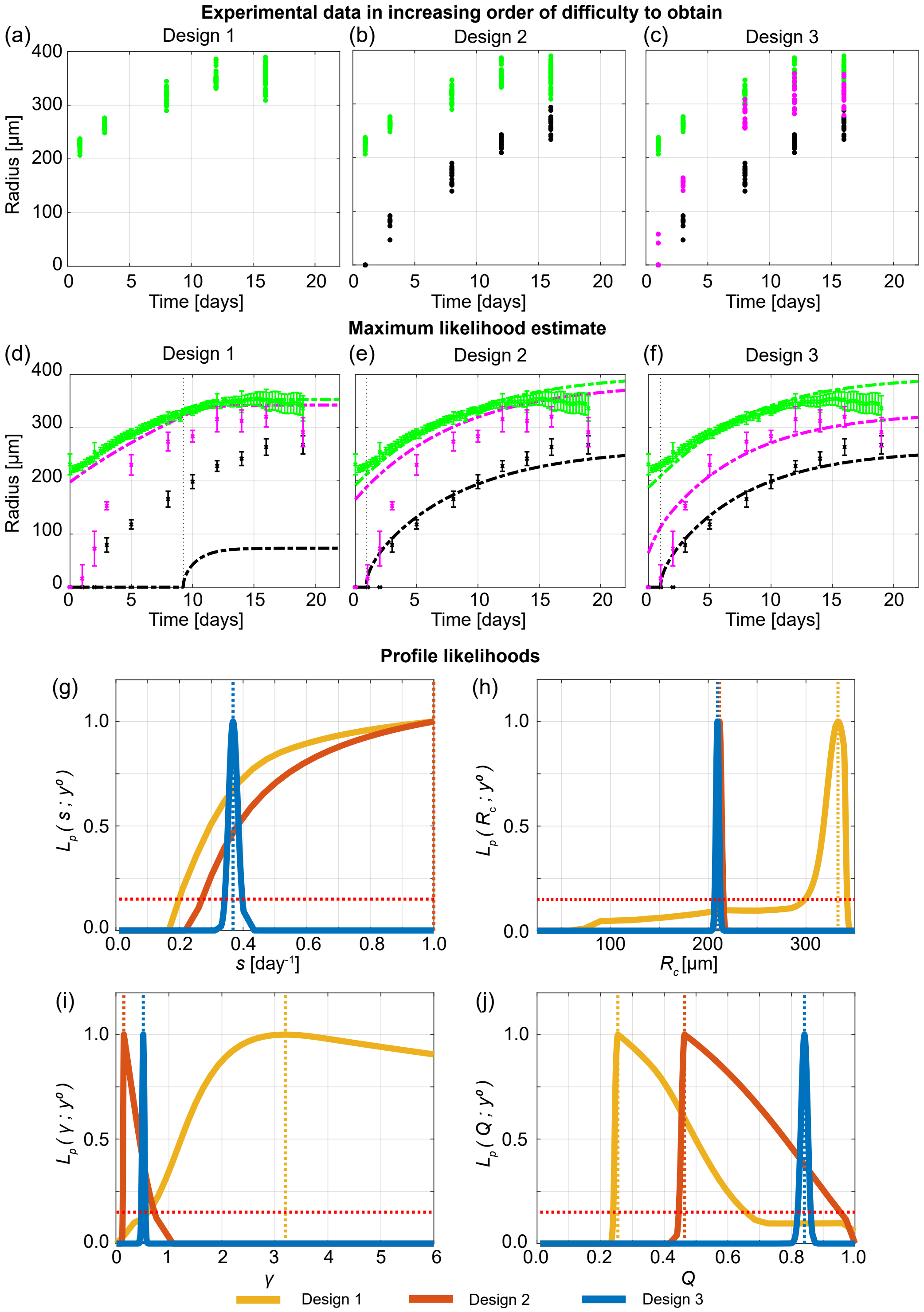


Figure S39: Measuring the necrotic and inhibited radius provides valuable information. (a)-(c) Experimental data used in Designs 1, 2 and 3. (d)-(f) Comparison of Greenspan model simulated with maximum likelihood estimate compared to full experimental data set for Designs 1, 2 and 3, where error bars show standard deviation. Profile likelihoods for (g) $s$, (h) $R_c$, (i) $\gamma$, (j) $Q$. Yellow, orange, blue lines in (g)-(j) represent profile likelihoods from Designs 1, 2, and 3, respectively, and the red-dashed line shows the approximate 95% confidence interval threshold. Results shown for WM983b spheroids formed with 10000 cells per spheroid.

#### F.3 Information gained across spheroid sizes is consistent

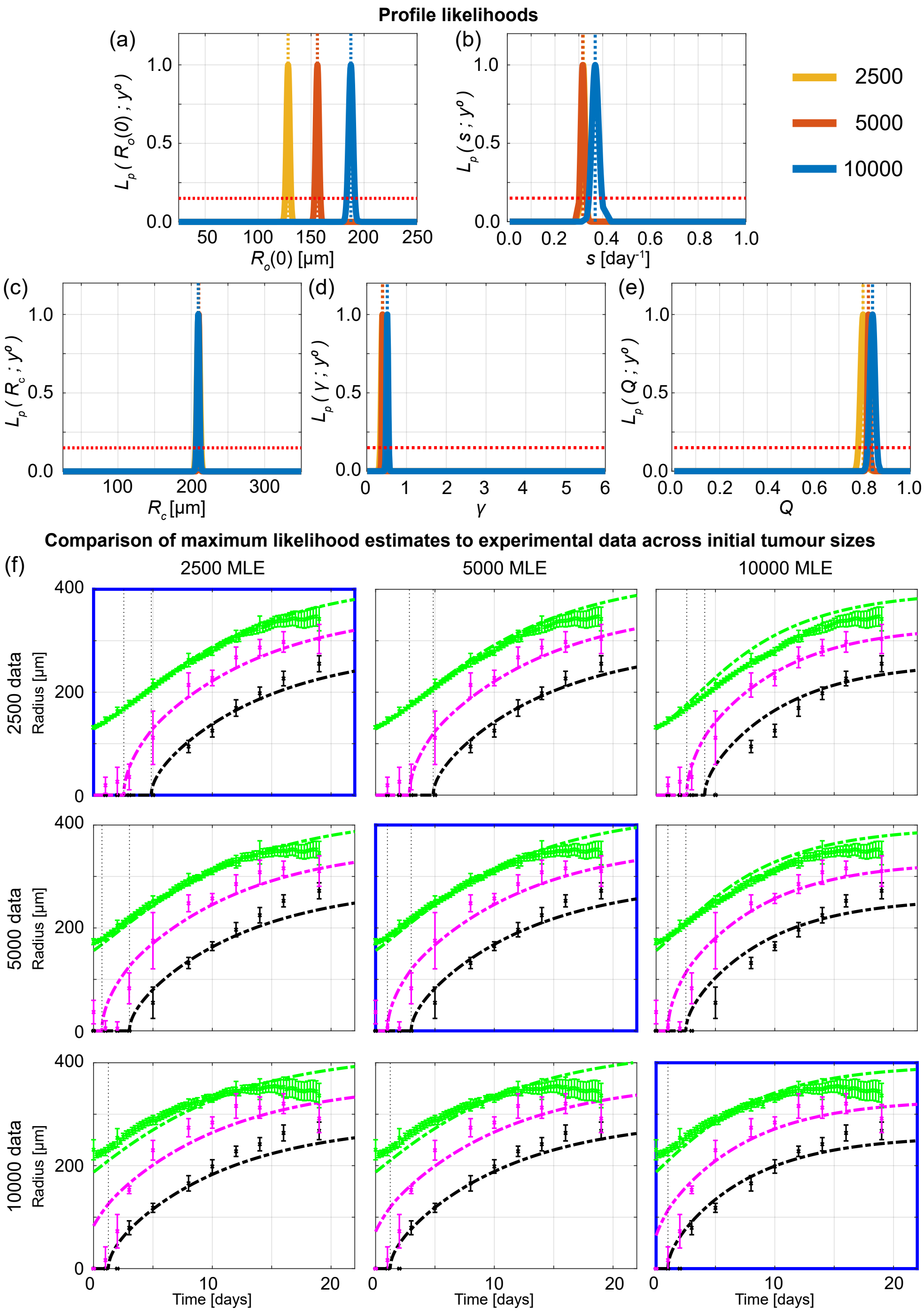


Figure S40: Information gained from experiments across different initial tumour spheroid sizes is mostly consistent. Profile likelihoods for (a) $R_o$, (b) $s$, (c) $R_c$, (d) $\gamma$, (e) $Q$. Yellow, orange, and blue lines in (a)-(e) represent profile likelihoods from tumour spheroids initially with approximately 2500, 5000, and 10000 cells per spheroid, respectively, and the red-dashed line shows the approximate 95% confidence interval threshold. (f) Comparison of Greenspan model simulated with maximum likelihood estimates compared to full experimental data sets across initial tumour spheroid size, where error bars show standard deviation. Results shown for WM983b cell line.

### G Parameter identifiability analysis for WM164

The main manuscript focuses on results for the human melanoma WM793b cell line. Here, we present analogous results for the human melanoma WM164 spheroids formed with 1250, 2500, 5000, and 10000 cells per spheroid. These spheroids are more challenging to interpret as we will now explain.

In experiments WM164 spheroids formed after 3 days. These spheroids were larger in size than other spheroids considered in this work, with the initial radius of WM164 spheroids formed with 1250 cells per spheroid larger than and similar size to WM983b and WM793b spheroids formed with 10000 cells per spheroid, respectively. The WM164 spheroids had relatively poor spherical symmetry [14], grew rapidly and many spheroids lost their structural integrity nine days after seeding formed with 1250, 2500, and 5000 cells per spheroid, and seven days after seeding for spheroids formed with 10000 cells spheroid. In addition, confocal microscopy could not be performed on day 7 after seeding for spheroids formed with 5000 and 10000 cells per spheroid due loss of structural integrity during harvesting. Identification of the necrotic region using image processing was more challenging, than for other cell lines, as a well-defined necrotic region did not form prior to the termination of the experiment. Therefore, necrotic region measurements for these spheroids are more subjective and uncertain. Spheroid boundaries were less smooth, so settings on the IncuCyte S3 live cell imaging system were updated to measure the largest brightfield object area with max eccentricity to 0.75 and sensitivity 20. These outer radius measurements were then manually reviewed to confirm accuracy.

We perform analysis for WM164 spheroids using Days 1, 2, 3, 4 and 5 after formation, where measurements could be obtained. This means that we do not include the last day of outer radius measurements for spheroids formed with 1250, 2500, and 5000 cells per spheroid. This allows us to compare the final outer radius measurement to Greenspan's model simulated with the MLE as a predictive test. However, for spheroids formed with 10000 cells per spheroid we include all data points so cannot form a predictive test, but this is because we seek to obtain as much information as possible in the shorter experimental duration. While the experimental duration for WM164 spheroids is relatively short in comparison to the WM793b and WM983b experiments, these experiments are still performed for multiple days longer than previous WM164 spheroid experiments [15].

To perform the analysis we update initial parameter bounds, used for practical parameter identifiability analysis, for as $200 < R_\mathrm{o}(0) < 600\,[\mu\mathrm{m}]$ and $200 < R_\mathrm{c} < 700\,[\mu\mathrm{m}]$. We update `FirstGuess` to $(Q, \gamma, s, R_\mathrm{c}, R_\mathrm{o}(0)) = (0.8, 0.1, 0.5, 400, 210)$ for spheroids formed with 1250 and 2500 cells per spheroid, and to $(Q, \gamma, s, R_\mathrm{c}, R_\mathrm{o}(0)) = (0.8, 0.1, 0.4, 400, 350)$ for spheroids formed with 5000 and 10000 cells per spheroid. Due to the reduced experimental duration for WM164 spheroids, and as we have already demonstrated with two other cell lines and synthetic data that Design 3 provides most insight, here we compare results obtained from spheroids with different initial sizes using Design 3.

In Figure S41a, we observe four distinct narrow peaks for $R_\mathrm{o}(0)$ corresponding to the four initial spheroid sizes, which is expected. For $s$, we observe that profile likelihoods overlap showing information obtained for $s$ is relatively consistent for different initial spheroid sizes (Figure S41b).

Interestingly and importantly, we observe four distinct peaks for $R_c$ (Figure S41c). This lack of consistency is different to the other two cell lines considered and strongly suggests information gained across initial spheroid sizes is not consistent. This is supported by direct observation of the experimental data where spheroids formed with 2500 cells have necrotic cores on day 4, whereas similar sized spheroids on day 2 formed with 5000 and 10000 cells per spheroid do not. Profile likelihoods for $\gamma$ are wide for spheroids formed with 5000 and 10000 cells per spheroid, and narrow and overlapping for spheroids formed with 1250 and 2500 cell per spheroid, showing that $\gamma$ requires more necrotic core measurements to be identified (Figure S41d). Profile likelihoods for $Q$ suggest that $Q$ decreases as the initial spheroid size increases (Figure S41e). This result for $Q$ is less consistent and in constrast to results from other cell lines, where the profiles for $Q$ overlapped for all spheroid sizes. Overall, we conclude that, possibly due to their lack of spherical symmetry, WM164 spheroids are more challenging to interpret and information gained using spheroids of different sizes is not consistent.

To support these results, we show along the diagonal of Figure S41f the solution of the mathematical model evaulated at the MLE associated with each initial spheroid size compared to the experimental measurements. In doing so we demonstrate that we accurately predict the last outer radius measurement using previous days measurements for spheroids formed with 1250, 2500, and 5000 cells per spheroid. However, on the off-diagonals of Figure 4f, we compare how the Greenspan model simulated with the MLE from one initial spheroid size predicts data from different initial spheroid sizes by only changing the initial radius. These off-diagonal results show that using information from one spheroid size to predict the behaviour of a different spheroid size is not always accurate. For example, using information gained from spheroids formed with 10000 cell per spheroid poorly predicts the behaviour of spheroids formed with 1250 cell per spheroid, as the time evolution of the outer radius is not accurately predicted at late time and the inhibited and necrotic regions form much earlier than predicted (top right of Figure S41f).

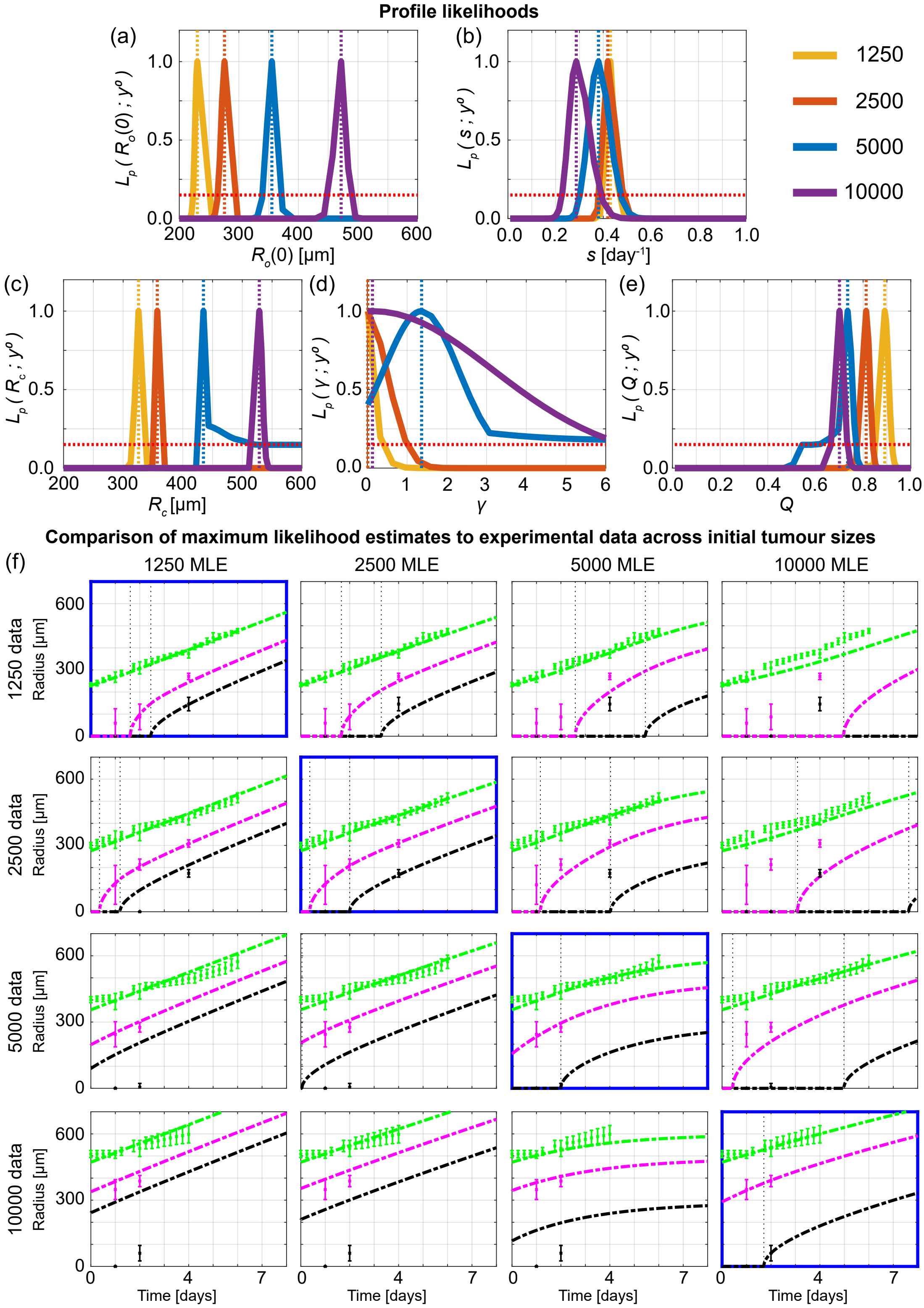


Figure S41: Information gained from WM164 experiments across different initial tumour spheroid sizes is inconsistent. Profile likelihoods for (a) $R_\mathrm{o}(0)$, (b) $s$, (c) $R_\mathrm{c}$, (d) $\gamma$, (e) $Q$. Yellow, orange, blue, and purple lines in (a)-(e) represent profile likelihoods from WM164 spheroids formed with 1250, 2500, 5000, 10000 cells per spheroid, respectively, and the red-dashed line shows the approximate 95% confidence interval threshold. (f) Comparison of Greenspan model simulated with maximum likelihood estimates compared to full experimental data sets across initial tumour spheroid size, where error bars show standard deviation.
